# *In vivo* functional profiling and structural characterisation of the human *Glp1r* A316T variant

**DOI:** 10.1101/2024.10.19.619191

**Authors:** Liliane El Eid, Kieran Deane-Alder, Roxana-Maria Rujan, Zamara Mariam, Affiong I. Oqua, Yusman Manchanda, Matthew J. Belousoff, Jorge Bernardino de la Serna, Kyle W. Sloop, Guy A. Rutter, Alex Montoya, Dominic J. Withers, Steven J. Millership, Karim Bouzakri, Ben Jones, Christopher A. Reynolds, Patrick M. Sexton, Denise Wootten, Giuseppe Deganutti, Alejandra Tomas

## Abstract

Glucagon-like peptide-1 receptor agonists (GLP-1RAs) are effective therapies for type 2 diabetes (T2D) and obesity, yet patient responses are variable. Variation in the human *Glp1r* gene might be directly linked to therapeutic responses. A naturally occurring missense variant, A316T, protects against T2D and cardiovascular disease. Here, we have generated and characterised a human *Glp1r* A316T mouse model. Human *Glp1r*^A316T/A316T^ mice displayed lower fasting blood glucose *versus* wildtype littermates, even under metabolic stress, and exhibited alterations in islet cytoarchitecture and α/β-cell identity under a high-fat, high-sucrose diet. This was however associated with blunted responses to GLP-1RAs *in vivo*. Further investigations in rodent and human β-cell models demonstrated that human *Glp1r* A316T exhibits characteristics of constitutive activation but dampened GLP-1RA responses. Results are further supported by cryo-EM analyses and molecular dynamics simulations of GLP-1R A316T structure, collectively demonstrating that the A316T variant governs basal GLP-1R activity and pharmacological responses to GLP-1R-targeting therapies.

**Teaser:** The *Glp1r* A316T missense variant displays improved glucose tolerance but impaired pharmacological incretin responses *in vivo*.

## Introduction

The glucagon-like peptide-1 receptor (GLP-1R), an important drug target for the treatment of type 2 diabetes (T2D), obesity, and cardiovascular disease, is a class B1 secretin-like G protein-coupled receptor (GPCR) predominantly expressed in the pancreas, but also present in other organs such as the brain, heart, stomach, intestine, and kidneys (*1–3*). The endogenous ligand for GLP-1R, GLP-1, is a 30 or 31 amino acid peptide incretin hormone secreted from intestinal L-cells upon food intake (*4*), unsuitable for pharmacological use due to its short half-life (∼2 minutes). Several pharmacological GLP-1 analogues, developed to bypass rapid degradation by dipeptidyl peptidase 4 (DPP-4) and, together with lipid modifications to reduce renal elimination, prolong pharmacokinetics (*5*), have been proven highly efficacious in controlling blood glucose levels and reducing body weight (*6, 7*).

Akin to other GPCRs, the GLP-1R undergoes a conformational change upon agonist binding, pleiotropically coupling with heterotrimeric G proteins (*8, 9*), with preferential engagement of Gα_s_, leading to activation of adenylate cyclase, generation of cyclic adenosine monophosphate (cAMP), potentiation of intracellular calcium, insulin granule mobilisation and exocytosis in pancreatic β-cells (*4, 10*). The GLP-1R can additionally inhibit glucagon secretion, delay gastric emptying, and suppress appetite (*4, 11*), with further physiological functions, including blood pressure regulation (*12, 13*), maintenance of renal function (*14, 15*), and neuroprotection (*16–19*), revealed by recent studies.

Despite the clinical success of GLP-1R agonists (GLP-1RAs), there is considerable heterogeneity amongst patient responses, with some exhibiting marked therapeutic improvements while others show little to no beneficial effect, and/or prominent adverse side effects (*20*). Natural genetic variants of human GPCRs can be linked to changes in receptor function, e.g., alterations in receptor structure, basal and ligand-dependent activity, and/or level of expression (*21*). Several *Glp1r* missense variants have been identified, some of which are associated with T2D risk and related glycaemic traits in genome-wide association studies (GWAS) (*22–25*). Large-scale sequencing and functional genetics have revealed a *Glp1r* missense variant (A316T; rs10305492; MAF=1.4%), associated with lower fasting glucose and reduced T2D and cardiovascular disease risk (*22–24*). The consequences of this variant on receptor function remain poorly defined, with conflicting results reported (*26, 27*). Previous analyses of variant receptor signalling *in vitro* suggest that it confers a gain-of-function (GoF) phenotype (*24, 28, 29*), but in-depth characterisation in primary tissues and *in vivo* has not been performed to date.

Here we present a detailed comparison between the wildtype (WT) and A316T GLP-1R, including functional characterisation in pancreatic β-cell lines and primary murine and human islets, as well as *in vivo* analysis of the A316T variant using a newly generated mouse model bearing a homozygous A316T (c.946G>A) substitution in the human *Glp1r* gene (*hGlp1r*^A316T/A316T^), both under physiological conditions and following diabetes induction by prolonged mouse feeding with a high-fat, high-sucrose (HFHS) diet. We additionally present the cryo-EM structure of the A316T GLP-1R bound to GLP-1, together with computational analyses, based on molecular dynamics (MD) simulations of WT *versus* A316T receptor, which elucidate the structural changes that underlie the functional effects of the A316T variant.

## Results

### Human Glp1r^A316T/A316T^ mice exhibit enhanced glucose homeostasis but reduced pharmacological responses to GLP-1RAs under different nutritional states

Previous GWAS results have led to the hypothesis that the A316T variant constitutively activates the GLP-1R, causing increased insulin secretion at lower ambient glucose levels, which in turn might trigger receptor down-regulation over time, potentially resulting in incretin resistance (*30*). To test this hypothesis *in vivo*, a mouse line expressing human GLP-1R from the murine *Glp1r* locus (*31*) was used to generate a global knock-in model harbouring the A316T single nucleotide polymorphism (SNP) (*hGlp1r*^A316T/A316T^), with WT *hGlp1r*^+/+^ littermates used as controls (fig. S1A and B). Mouse genotypes were confirmed by Sanger sequencing, and glucose homeostasis analysed by crossover studies in a mixed sex cohort.

Despite no discernible weight differences between *hGlp1r*^A316T/A316T^ and WT littermates on a chow diet (Fig. 1A), glucose tolerance was improved following an oral gavage (Fig. 1B), and fasting blood glucose levels were significantly lower in *hGlp1r*^A316T/A316T^ mice (Fig. 1C), the later result in agreement with GWAS data (*22–24*). To examine the acute *versus* sustained pharmacogenetic effects of the GLP-1R A316T variant, we performed intraperitoneal glucose tolerance tests (IPGTTs) immediately after (acute) and six hours post-administration (sustained) of vehicle or the GLP-1RAs exendin-4 or exendin-F1, which have differing biased agonism profiles when assessed *in vitro* at the WT GLP-1R (*32*). We observed significantly improved acute glucose responses to vehicle administration in *hGlp1r*^A316T/A316T^ *versus hGlp1r*^+/+^ littermates, with a similar trend at the 6-hour time-point (Fig. 1D and E). Conversely, *hGlp1r*^A316T/A316T^ and *hGlp1r*^+/+^ mice exhibited similar glucose responses to exendin-4 both acutely and 6-hours post-agonist administration (Fig. 1F and G), with comparably attenuated glucose effects at the 6-hour time-point, associated with exendin-4-induced receptor desensitisation (*32*). A similar trend could be observed with exendin-F1, which, as expected by its bias profile away from β-arrestin recruitment, displayed reduced desensitisation at the 6-hour IPGTT for both mouse genotypes *versus* exendin-4 (*32*), but with prolonged glucose levels slightly improved for *hGlp1r*^A316T/A316T^ mice (Fig. 1H), without reaching statistical significance when measured as areas under the curve (AUCs) *versus hGlp1r*^+/+^ mice (Fig. 1I). Direct comparison between AUCs for the different conditions and genotypes per time-point is shown in figs. S2A and B. Agonist-specific glycaemic responses (ΔAUC, corresponding to agonist-minus vehicle-treated AUCs, measurable due to the crossover design of the study) revealed significantly reduced acute exendin-4 responses in *hGlp1r*^A316T/A316T^ *versus hGlp1r*^+/+^ littermates, a difference that was maintained at the 6-hour time-point, when exendin-4 treatment resulted in a paradoxical increase in blood glucose levels compared to vehicle in *hGlp1r*^A316T/A316T^ mice (Fig. 1J). Interestingly, while *hGlp1r*^A316T/A316T^ mice displayed similarly reduced acute responses to exendin-F1 when compared to *hGlp1r*^+/+^ mice, this difference was absent after 6 hours, suggesting a beneficial impact of the reduced desensitisation afforded by this agonist (*32*) for the mice expressing the A316T variant (Fig. 1K). These effects were also apparent when calculated as vehicle fold changes (fig. S2C). No significant differences were detected per genotype at prolonged minus acute time-points (fig. S2D and E), indicating that the loss of effect in response to GLP-1RAs is already present acutely. Similar experiments in heterozygous *hGlp1r*^A316T/+^ mice showed no clear differences *versus* WT *hGlp1r*^+/+^ littermates under a chow diet (fig. S2, F to P).

**Fig. 1.**
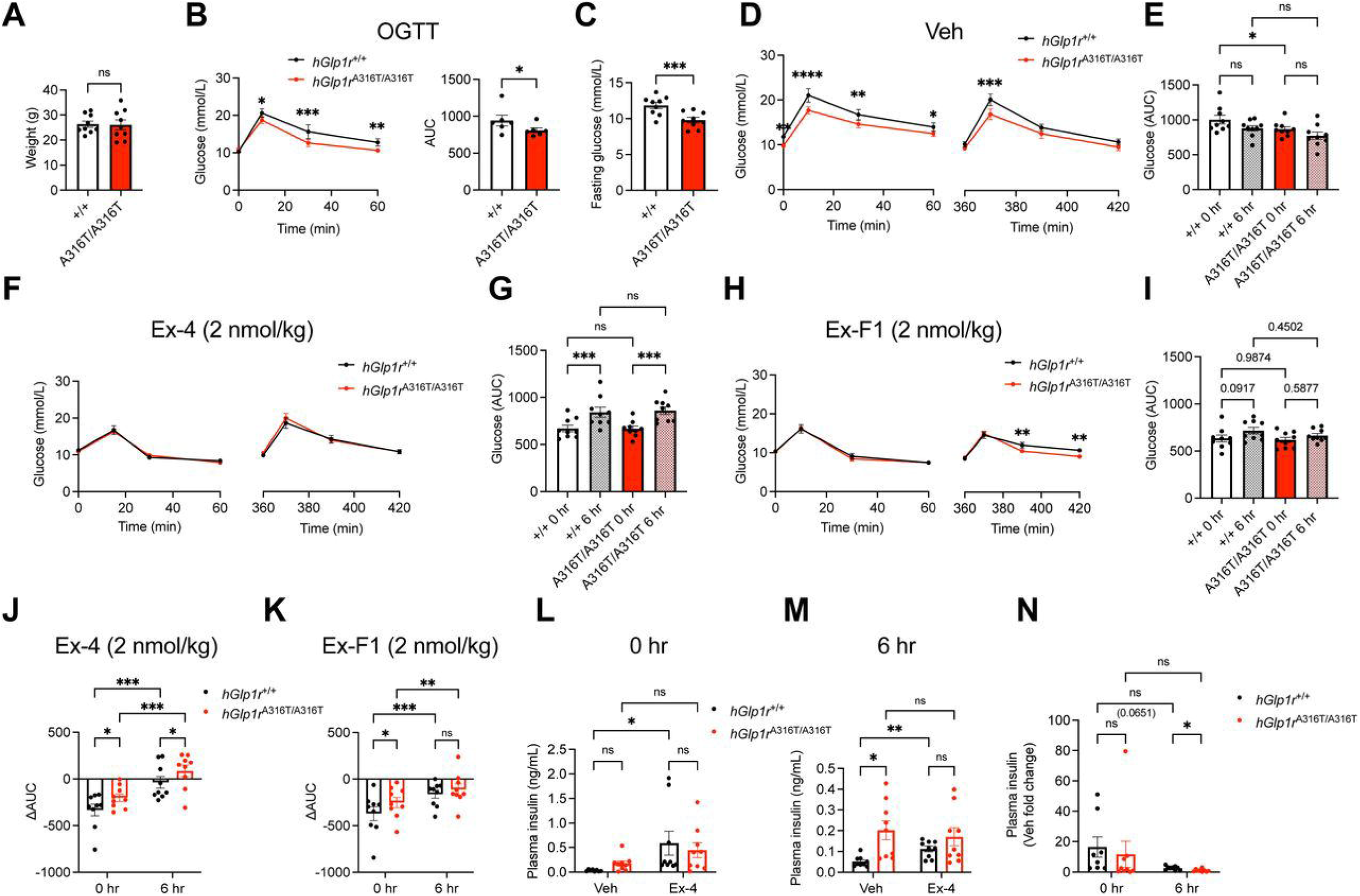
*In vivo* glucose responses of adult *hGlp1r*^A316T/A316T^ *versus hGlp1r*^+/+^ mice on a chow diet. (**A**) Age-matched body weight of *hGlp1r*^+/+^ *versus* ^A316T/A316T^ mixed sex adult littermate mice (*n* = 9 per genotype) maintained on a normal chow diet. (**B**) OGTT responses and corresponding AUCs following 2 g/kg body weight glucose administration via oral gavage after 5 hours of fasting in a mixed sex cohort of lean, adult *hGlp1*^+/+^ *versus* ^A316T/A316T^ mice on a chow diet (*n* = 6 per genotype). (**C**) Glucose levels after 2 hours of fasting from mice in (A). (**D** and **E**) IPGTTs (2 g/kg body weight glucose) following acute (0 hours), or 6 hours post-intraperitoneal administration of saline vehicle (Veh) in chow-fed *hGlp1r*^+/+^ *versus* ^A316T/A316T^ mixed sex adult mice. Glucose curves (D) and corresponding AUCs (E) shown; *n* = 9 mice per genotype. (**F** to **I**), As in (D and E) following administration of 2 nmol/kg exendin-4 (Ex-4) (F and G) or exendin-F1 (Ex-F1) (H and I). (**J** and **K**) Vehicle-corrected ΔAUC responses from (G) and (I), respectively. (**L** and **M**) Plasma insulin levels acutely (L) and 6 hours (M) post-administration of Veh or Ex-4 in chow-fed *hGlp1r*^+/+^ *versus* ^A316T/A316T^ mixed sex adult mice. (N) Plasma insulin vehicle fold changes from (L and M). Data are mean ± SEM; *p<0.05; **p<0.01; ***p< 0.001; ****p<0.0001; ns, not significant by paired t-tests, one- or two-way ANOVA with Sidak’s post-hoc tests.

Plasma samples collected 10 minutes into the IPGTTs showed higher insulin levels in *hGlp1r*^A316T/A316T^ *versus hGlp1r*^+/+^ littermates after vehicle administration, an effect that became significant for the 6-hour time-point (Fig. 1, L to M). As expected, plasma insulin levels increased in response to exendin-4 for the *hGlp1r*^+/+^ mice, but this effect was attenuated in *hGlp1r*^A316T/A316T^. When expressed as vehicle fold changes, exendin-4-dependent plasma insulin rises were reduced for *hGlp1r*^A316T/A316T^ *versus hGlp1r*^+/+^ littermates, a difference that became significant at the 6-hour time-point (Fig. 1N).

We next subjected our mouse cohort from above to 12-14 weeks of HFHS diet feeding to investigate variant-associated phenotypes under diet-induced metabolic stress. During the HFHS feeding period, *hGlp1r*^A316T/A316T^ mice displayed slower weight gain compared to *hGlp1r*^+/+^ littermates (Fig. 2A and B) but still reached the same final weight at the end of the HFHS feeding period, allowing us to assess glucoregulatory effects without confounding differences in body weight. Consistent with our results in lean conditions, oral glucose tolerance was significantly improved in *hGlp1r*^A316T/A316T^ *versus hGlp1r*^+/+^ mice (Fig. 2C), but no changes were now observed in fasting glycemia between genotypes (Fig. 2D). HFHS-fed *hGlp1r*^A316T/A316T^ mice exhibited significantly enhanced intraperitoneal (ip) glucose tolerance under vehicle conditions compared to *hGlp1r*^+/+^ mice, both acutely and at the 6-hour time-point (Fig. 2E and F). Glucose responses to exendin-4 and exendin-F1 resembled those of *hGlp1r*^+/+^ mice, but with slightly more pronounced desensitisation for *hGlp1r*^A316T/A316T^ mice (Fig. 2, G to J). Direct comparison between AUCs for the different conditions and genotypes per time-point is shown in fig. S3A and B. When calculated as vehicle ΔAUC, the glucose-lowering effect of both GLP-1RAs was significantly reduced in HFHS-fed *hGlp1r*^A316T/A316T^ *versus hGlp1r*^+/+^ littermates, with differences apparent both acutely and 6-hours post-administration of either agonist (Fig. 2K and L). Effects were also apparent when calculated as vehicle fold changes (fig. S3C), and no significant differences in prolonged minus acute effect per agonist and genotype were detected, indicating that the loss of GLP-1RA responses in *hGlp1r*^A316T/A316T^ mice is already present acutely (fig. S3D and E). Basal plasma insulin levels were once more increased in HFHS-fed *hGlp1r*^A316T/A316T^ *versus hGlp1r*^+/+^ mice, but now significantly also at the acute time-point (fig. S3F).

**Fig. 2.**
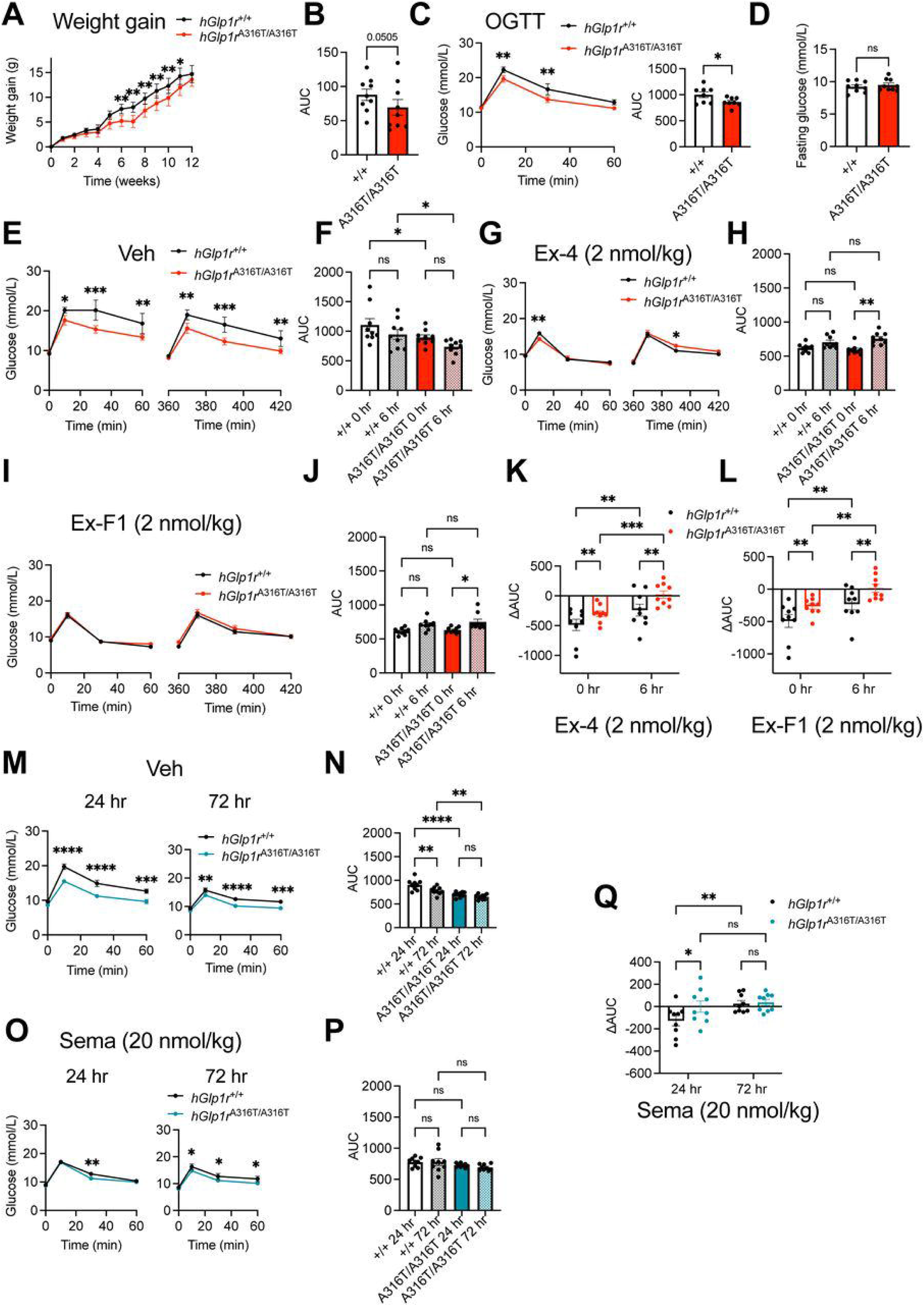
*In vivo* glucose responses of adult *hGlp1r*^A316T/A316T^ *versus hGlp1r*^+/+^ mice on a HFHS diet. (**A**) Body weight gain for *hGlp1r*^+/+^ *versus* ^A316T/A316T^ mixed sex adult littermate mice (*n* = 9 per genotype) maintained on a HFHS diet for the indicated times. (**B**) AUCs from (A). (**C**) OGTT responses and corresponding AUCs following 2 g/kg body weight glucose administration via oral gavage after 5 hours of fasting in a mixed sex cohort of adult *hGlp1r*^+/+^ *versus* ^A316T/A316T^ mice on a HFHS diet (*n* = 8 per genotype). (**D**) Glucose levels after 2 hours of fasting in mice from (A). (**E** and **F**) IPGTTs (2 g/kg body weight glucose) following acute (0 hours), or 6 hours post-intraperitoneal administration of saline vehicle (Veh) in HFHS-fed *hGlp1r*^+/+^ *versus* ^A316T/A316T^ mixed sex adult mice. Glucose curves (E) and corresponding AUCs (F) shown; *n* = 9 mice per genotype. (**G** to **J**) As in (E and F) following administration of 2 nmol/kg exendin-4 (Ex-4) (G and H) or exendin-F1 (Ex-F1) (I and J). (**K** and **L**) Vehicle-corrected ΔAUC responses from (H) and (J), respectively. (**M** to **P**) Glucose curves and corresponding AUCs following vehicle (Veh) (M and N), or 20 nmol/kg semaglutide (Sema) (O and P) treatment of HFHS-fed *hGlp1r*^+/+^ *versus* ^A316T/A316T^ mice; IPGTTs performed 24- and 72-hours post-agonist administration. (**Q**) Vehicle-corrected ΔAUC responses from (P). Data are mean ± SEM; *p<0.05; **p<0.01; ***p< 0.001; ****p<0.0001; ns, not significant by paired t-tests, one- or two-way ANOVA with Sidak’s post-hoc tests.

To assess the effects of the A316T variant in response to a GLP-1RA with prolonged pharmacokinetics, we conducted additional experiments in HFHS-fed animals treated with the long-acting GLP-1RA semaglutide (*33*). We observed a similar profile of enhanced glucose tolerance under vehicle conditions at 24- and 72-hours in HFHS-fed *hGlp1r*^A316T/A316T^ *versus hGlp1r* ^+/+^ littermates (Fig. 2M and N). Glucose responses to semaglutide were very similar to those of *hGlp1r*^+/+^ littermates, with slightly lower glucose levels across the time course for *hGlp1r*^A316T/A316T^ but no significant changes in AUC (Fig. 2O and P). Direct comparison between AUCs per time-point is shown in fig. S3, G to H. Results calculated as ΔAUC showed that semaglutide induces lower responses in *hGlp1r*^A316T/A316T^ *versus hGlp1r*^+/+^ littermates 24 hours post-agonist treatment, with these differences no longer apparent at the 72-hour time-point, when both genotypes display fully desensitised responses (Fig. 2Q). These effects were similarly apparent when calculated as vehicle fold changes (fig. S3I). Consequently, we observed a nearly significant reduction in prolonged minus acute effects for semaglutide in *hGlp1r*^A316T/A316T^ compared to *hGlp1r*^+/+^ mice (Fig. S3J), indicating that the loss of response to semaglutide in *hGlp1r*^A316T/A316T^ mice occurs acutely, potentially due to accelerated receptor desensitisation. Importantly, despite heterozygous *hGlp1r*^A316T/+^ mice showing more attenuated differences *versus hGlp1r*^+/+^ littermates for weight gain, oral glucose tolerance, fasting glucose, and acute *versus* 6-hour IPGTT responses (fig. S4, A to L) under a HFHS diet, *hGlp1r*^A316T/+^ mice exhibited significantly increased desensitisation 6-hours post-exendin-4 administration (fig. S4G and H), as well as improved glucose tolerance 24-hours post-vehicle administration (fig. S4M and N), which led to significantly worse ΔAUC responses to semaglutide *versus hGlp1r*^+/+^ littermates (fig. S4, O to Q), an effect that, as for exendin-4 administration in chow-fed mice, led to paradoxically higher glucose levels compared to those of vehicle-treated *hGlp1r*^A316T/+^ mice (fig. S4Q).

### Alterations of islet cytoarchitecture and α/β-cell identity in human Glp1r^A316T/A316T^ mice

Previous studies suggest that, due to its proliferative and anti-apoptotic effects, prolonged GLP-1R activity might lead to alterations in critical aspects of the islet cytoarchitecture such as size, cellular composition, and α/β-cell identity (*34–37*). Changes in islet cytoarchitecture have also been linked to β-cell dysfunction and T2D progression (*38–41*). With the A316T variant conferring a GoF phenotype in both oral glucose tolerance and ip vehicle responses, indicative of increased basal GLP-1R activity, we hypothesised that it might also lead to changes in islet cytoarchitecture and α- and/or β-cell identity, a possibility that has not been assessed in previous *in vitro* analyses of this variant. To test this, we conducted histological examinations of pancreatic tissue and islets isolated from chow- and HFHS-fed *hGlp1r*^A316T/A316T^ *versus hGlp1r*^+/+^ mice. Chow-fed *hGlp1r*^A316T/A316T^ islets showed a non-significant tendency for increased numbers of both insulin- and glucagon-positive cells, without changes in percentages (Fig. 3A and B). HFHS-fed *hGlp1r*^A316T/A316T^ islets, however, displayed a remarkable increase in the percentage of glucagon-positive cells, and a concomitant decrease in insulin-positive cells compared to *hGlp1r*^+/+^ islets (Fig. 3C and D). Results were recapitulated in whole pancreas sections: while chow-fed mice did not show significant differences in β- or α-cell mass, α-to-β cell ratio, or islet diameter between genotypes (Fig. 3, E to H), immunolabelling of pancreas sections from HFHS-fed *hGlp1r*^A316T/A316T^ mice revealed increased β-cell mass, α-to-β-cell ratio, and islet diameter *versus hGlp1r*^+/+^ (Fig. 3, I to L), appearing in response to the HFHS diet. Interestingly, β-cells from HFHS-fed *hGlp1r*^A316T/A316T^ islets also showed a distinct loss of localisation to the islet periphery (the two outermost islet cell layers) and were instead intermingled with β-cells throughout the islet core (Fig. 3M). Closer examination of intra-islet endocrine cell localisation revealed a significant proportion of non-peripheral cells expressing both insulin and glucagon (Fig. 3I, arrowheads), potentially indicating β- to- βcell trans-differentiation in response to HFHS diet in *hGlp1r*^A316T/A316T^ islets. Analysis of expression of β-cell “enriched” and “disallowed” genes revealed upregulation of *Mafa*, a gene associated with β-cell identity, in islets from both chow- and HFHS-fed *hGlp1r*^A316T/A316T^ mice, with no changes in β-cell disallowed gene expression (*Slc16a1*, *Ldha*), and only a tendency for increased expression of β-cell-specific *Gcg* (fig. S5A and B). Interestingly, expression of the β-cell disallowed gene *Acot7* (*42*) was significantly upregulated in chow-fed *hGlp1r*^A316T/A316T^ islets.

**Fig. 3.**
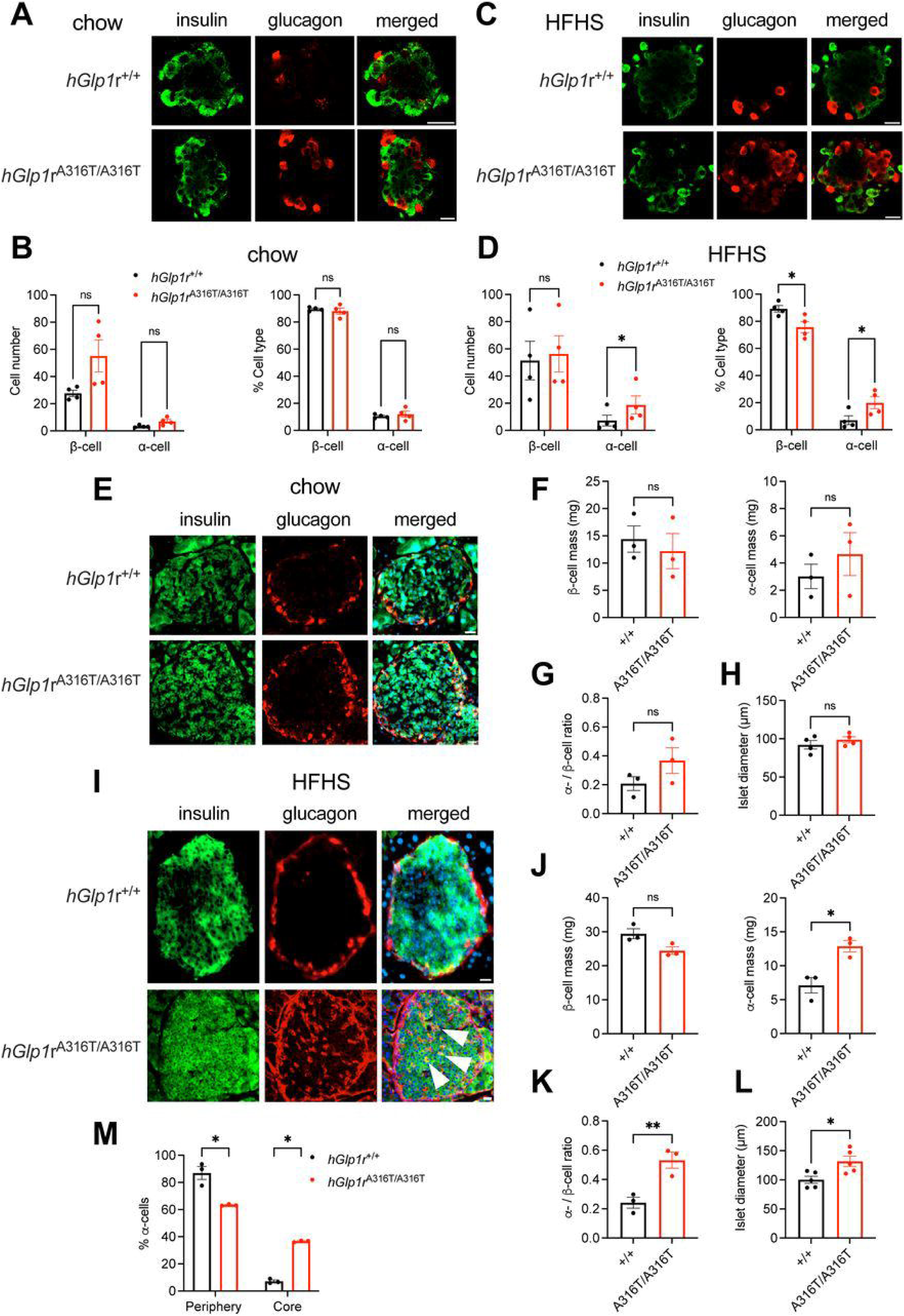
Morphological analyses of *hGlp1r*^A316T/A316T^ *versus hGlp1r*^+/+^ mouse islets. (**A** and **B**) Immunofluorescence analysis of insulin (green), and glucagon (red) localisation in isolated islets from chow-fed *hGlp1r*^+/+^ *versus* ^A316T/A316T^ mice; size bars, 20 µm. Representative images (A) and α-*versus* β-cell number and percentage quantifications (B); *n* = 4. (**C** and **D**) As in (A and B) for islets from HFHS-fed *hGlp1r*^+/+^ *versus* ^A316T/A316T^ mice; *n* = 4. (**E** to **H**) Immunofluorescence analysis of insulin (green), and glucagon (red) localisation in pancreas sections from chow-fed *hGlp1r*^+/+^ *versus* ^A316T/A316T^ mice; nuclei (DAPI), blue; size bars, 20 µm. Representative images (E), quantification of α- and β-cell number (F), α-over β-cell ratio (G) and islet diameter (H); *n* = 3-4. (**I** to **L**) as for (**E** to **H**) in pancreas sections from HFHS-fed *hGlp1r*^+/+^ *versus* ^A316T/A316T^ mice. Representative images (I), quantification of α-and β-cell number (J), α-over β-cell ratio (K) and islet diameter (L); *n* = 3-5; size bars, 20 µm; arrowheads indicate cells positive for both insulin and glucagon. (**M**) Percentage of islet peripheral/mantle *versus* core α-cells in pancreas sections from HFHS-fed *hGlp1r*^+/+^ *versus* ^A316T/A316T^ mice; *n* = 3. Data are mean ± SEM; *p<0.05; **p<0.01; ns, not significant by paired t-tests or two-way ANOVA with Sidak’s post-hoc tests.

### Changes in cell surface expression, agonist-induced cAMP and insulin secretion in human Glp1r^A316T/A316T^ mouse islets

Accounting for potential variations in cell surface expression of GLP-1R variants is crucial, as differences in this parameter can influence baseline receptor activity and subsequent GLP-1RA responses (*29*). Of note, the humanised *Glp1r* mouse model employed in this study displays reduced levels of GLP-1R surface expression compared to non-humanised mice (fig. S5C and D), potentially due to *hGlp1r* being expressed as a mini-gene inserted in the murine *Glp1r* locus (*31*). We therefore have restricted our comparisons to humanised *Glp1r* mice with the same genetic background to avoid any confounding effects. Islets extracted from both chow- and HFHS-fed *hGlp1r*^A316T/A316T^ mice showed reduced surface GLP-1R levels *versus hGlp1r*^+/+^, assessed by labelling with a saturating concentration of the fluorescent GLP-1R antagonist exendin-9-TMR. Results were corroborated *in vivo* by subcutaneous injection of exendin-9-Cy5 [LUXendin645 (*2*)] prior to transcardial fixation (Fig. 4A and B).

**Fig. 4.**
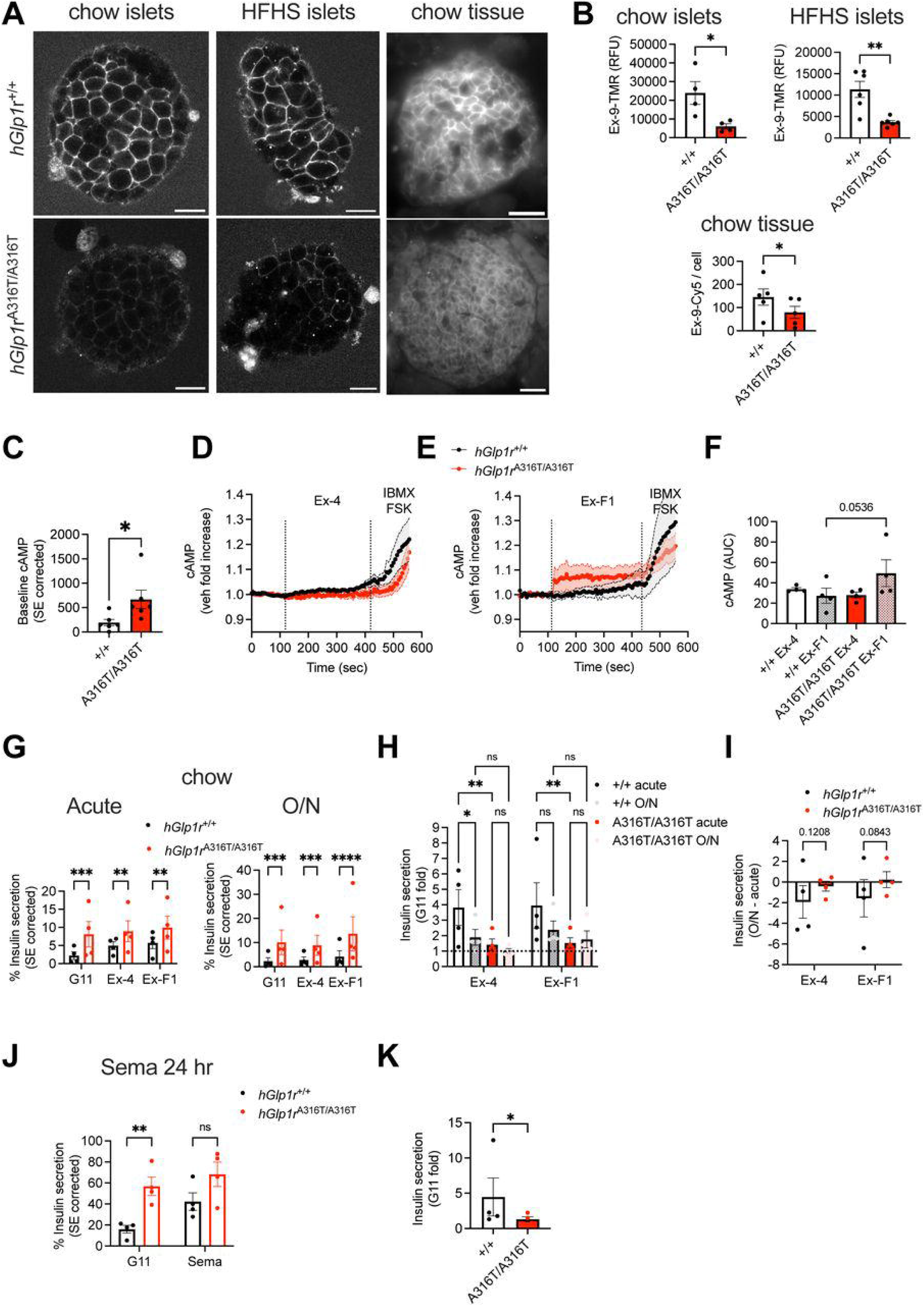
GLP-1R surface expression and downstream signalling in *hGlp1r*^A316T/A316T^ *versus hGlp1r*^+/+^ mouse islets. (**A** and **B**) GLP-1R surface expression, assessed *ex vivo* with exendin-9-TMR (Ex-9-TMR) in purified chow and HFHS islets, or *in vivo* following subcutaneous injection of exendin-9-Cy5 (Ex-9-Cy5, LUXendin645) and fluorescence imaging of pancreatic tissue sections of *hGlp1r*^+/+^ *versus* ^A316T/A316T^ mice. Representative images (A) and quantifications of surface GLP-1R levels (B); size bars, 20 µm; *n* = 4-6. (**C**) Basal cAMP responses measured with the Green Up cADDis cAMP biosensor in chow-fed *hGlp1r*^+/+^ *versus* ^A316T/A316T^ mouse islets; *n* = 6. (**D** and **E**) cAMP responses to 100 nM exendin-4 (Ex-4) (D) or exendin-F1 (Ex-F1) (E) in islets from (C); responses normalised to baseline; IBMX and forskolin (FSK) added at the acquisition end for maximal responses; *n* = 4. (**F**) AUCs calculated from agonist-stimulated periods in (**D** and **E**); *n* = 4 per genotype. (**G**) Percentage of acute and overnight (O/N) surface expression (SE)-corrected insulin secretion in chow-fed *hGlp1r*^+/+^ *versus* ^A316T/A316T^ mouse islets stimulated with 11 mM glucose (G11) alone or supplemented with 100 nM exendin-4 (Ex-4) or exendin-F1 (Ex-F1); *n* = 4. (**H**) Insulin secretion fold changes to G11 calculated from data in (G). (**I**) Overnight over acute insulin secretion responses calculated from data in (H). (**J**) Percentage of SE-corrected insulin secretion in chow-fed *hGlp1r*^+/+^ *versus* ^A316T/A316T^ mouse islets stimulated for 24 hours with 11 mM glucose (G11) alone or supplemented with 100 nM semaglutide; *n* = 4. (**K**) Insulin secretion fold changes to G11 calculated from data in (J). Data are mean ± SEM; *p<0.05; **p<0.01; ***p< 0.001; ****p<0.0001; ns, not significant by paired t-tests, one- or two-way ANOVA with Sidak’s post-hoc tests.

We next analysed acute cAMP responses in chow-fed *hGlp1r*^A316T/A316^ *versus hGlp1r*^+/+^ islets transduced with the baculoviral cAMP biosensor cADDis (*43*). Once corrected for surface expression, basal cAMP was significantly increased in *hGlp1r*^A316T/A316^ relative to *hGlp1r*^+/+^ islets (Fig. 4C). While agonist-induced responses were not significantly different between genotypes, we observed a trend for reduced cAMP in response to exendin-4, whereas, conversely, there was a near-significant increase in cAMP in response to exendin-F1 in *hGlp1r*^A316T/A316^ islets (Fig. 4, D to F). *Ex vivo* assays of insulin secretion from the same islets showed increased outputs in response to both acute (30 minutes) and sustained (16 hours, cumulative) exposure to 11 mM glucose in *hGlp1r*^A316T/A316^ relative to *hGlp1r*^+/+^ (Fig. 4G). Agonist-mediated increases in glucose-stimulated insulin secretion, measured as fold over 11 mM glucose responses, were however blunted in *hGlp1r*^A316T/A316^ *versus hGlp1r*^+/+^ islets, with complete loss of the potentiating effect of exendin-4 (∼1-fold) but, interestingly, no significantly detrimental effect for exendin-F1 following overnight agonist exposure (Fig. 4H). As *in vivo*, no differences were detected in prolonged minus acute responses for either agonist *versus hGlp1r*^+/+^ islets, although there was a tendency for the bulk of the A316T detrimental effect to occur acutely (Fig. 4I). Parallel experiments in islets from HFHS-fed mice showed similar tendencies which, however, did not reach statistical significance (fig. S5, E to G). Secretion responses to 24-hours stimulation with semaglutide again showed a significant decrease in agonist-induced potentiation of glucose-mediated insulin secretion in islets from HFHS-fed *hGlp1r*^A316T/A316^ *versus hGlp1r*^+/+^ mice (Fig. 4I and J).

Further experiments were performed using islets from *hGlp1r*^-/-^ mice, generated *in house* from the *hGlp1r*^+/+^ mouse line, subsequently transduced with adenoviral vectors expressing SNAP/FLAG-tagged WT or A316T hGLP-1R (pAV-SNAP/FLAG-*hGlp1r*^WT^ or ^A316T^). Using a membrane impermeable fluorescent SNAP tag probe, we again measured reduced surface expression of SNAP/FLAG-*hGlp1r*^A316T^ compared to its WT counterpart (fig. S6, A and B). Intracellular Ca^2+^ mobilisation, another downstream signalling readout associated with insulin granule exocytosis, was examined using the fluorescent calcium indicator Cal-520 AM. A clear increase in basal Ca^2+^ levels was apparent in SNAP/FLAG-*hGlp1r*^A316T^-*versus* SNAP/FLAG-*hGlp1r*^WT^-expressing islets (fig. S6C). Agonist-induced calcium responses were not significantly different between WT and A316T for GLP-1, exendin-4 or semaglutide, but we observed significantly increased Ca^2+^ mobilisation in response to exendin-F1 in *hGlp1r*^-/-^ pAV-SNAP/FLAG-*hGlp1r*^A316T^ islets (fig. S6, D to H), in line with our previous cAMP results with this biased GLP-1RA.

### Impact of the A316T variant on primary human islets and a human β-cell line

Given the strong evidence from previous GWAS and *in vitro* HEK293 data (*44*), and our *in vivo* observations of the effect of the GLP-1R A316T variant on glucose homeostasis, we next investigated its impact on human β-cell models, including in human donor islets and in the well-characterised human β-cell line EndoC-βH3 (*45*) genetically modified by CRISPR/Cas9 to generate a *Glp1r*^-/-^-enriched multiclonal cell population and then transduced with adenoviruses expressing SNAP/FLAG-*hGlp1r*^WT^ or ^A316T^ as above. EndoC-βH3 *Glp1r*^-/-^ SNAP/FLAG-*hGlp1r*^A316T^ cells again showed reduced GLP-1R cell surface expression (Fig. 5A and B), and similar insulin secretion profiles to those observed in *hGlp1r*^A316T/A316T^ mouse islets, with enhanced basal insulin secretion but significantly reduced exendin-4-stimulated fold increases (Fig. 5C and D). Experiments were translated to primary human islets transduced with SNAP/FLAG-*hGlp1r*^WT^ or^A316T^ adenoviruses, a system that mimics a heterozygous phenotype for the A316T variant expression. Results again showed reduced surface expression of SNAP/FLAG-*hGlp1r*^A316T^ *versus* ^WT^ (Fig. 5E and F), as well as a near-significant loss of exendin-4-induced potentiation of insulin secretion (Fig. 5G and H). Interestingly, both human A316T models retained some degree of exendin-4 responsiveness compared to glucose alone responses, which was for the most part absent in islets from *hGlp1r*^A316T/A316^ mice.

**Fig. 5.**
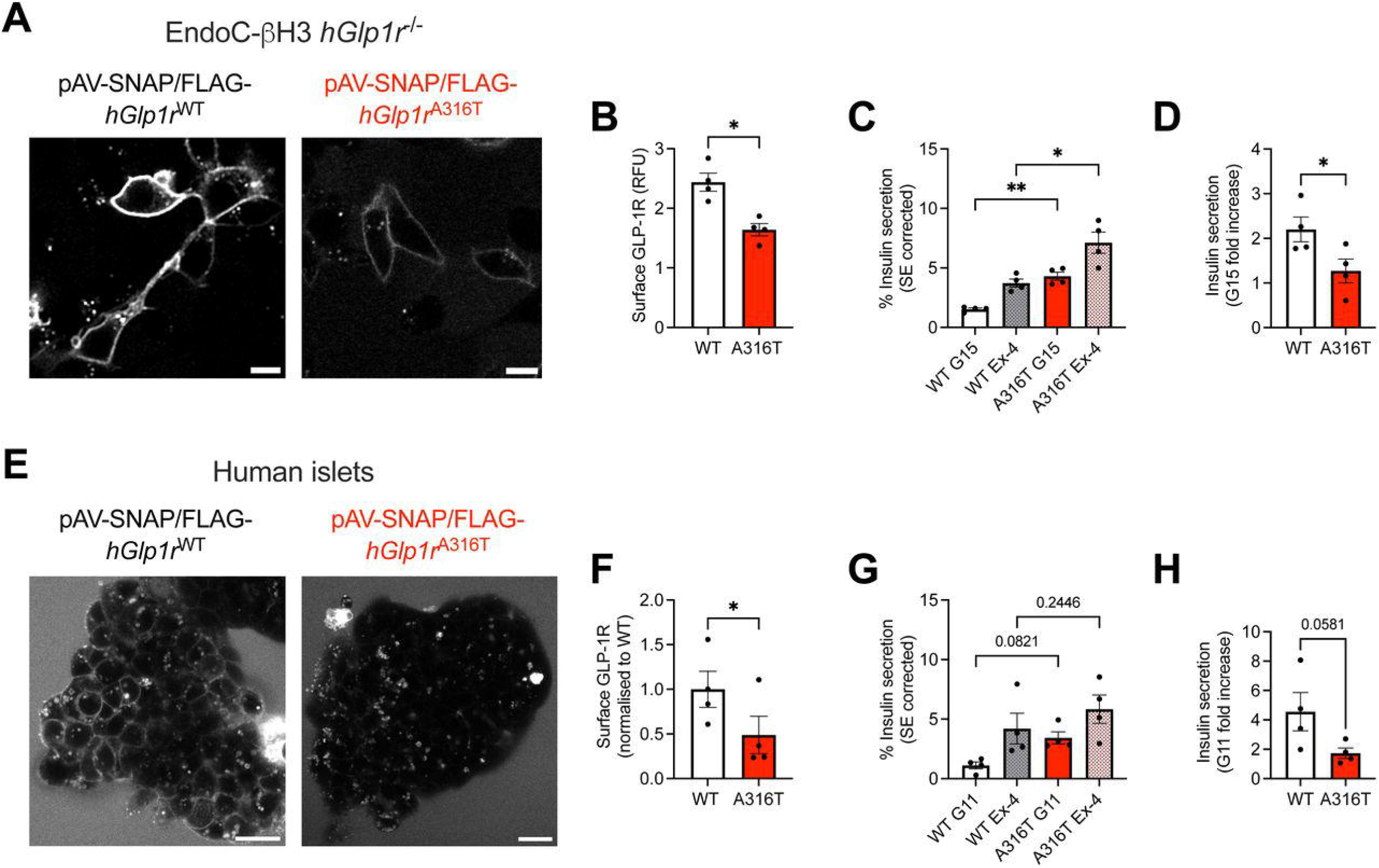
SNAP/FLAG-*hGlp1r* ^WT^ *versus* ^A316T^ surface expression levels and insulin secretion in human β-cell models. (**A** and **B**) Surface expression in EndoC-βH3 *hGlp1r*^-/-^ cells transduced with pAV-SNAP/FLAG-*hGlp1r*^WT^ or ^A316T^ labelled with SNAP-Surface Alexa Fluor 647. Representative images (A) and quantification of surface GLP-1R (B); size bars, 10 μm; *n* = 4. (**C**) Percentage of surface expression (SE)-corrected insulin secretion in EndoC-βH3 *hGlp1r*^-/-^ cells transduced with pAV-SNAP/FLAG-*hGlp1r*^WT^ or ^A316T^ in response to 15 mM glucose (G15) alone or supplemented with 100 nM exendin-4 (Ex-4); *n* = 4. (**D**) Insulin secretion fold changes to G15 calculated from data in (C). (**E** and **F**) Surface expression in primary human islets transduced with pAV-SNAP/FLAG-*hGlp1r*^WT^ or ^A316T^ labelled with SNAP-Surface Alexa Fluor 647. Representative images (E) and quantification of surface GLP-1R (F); size bars, 20 μm; *n* = 4. (**G**) Percentage of SE-corrected insulin secretion in primary human islets transduced with pAV-SNAP/FLAG-*hGlp1r*^WT^ or ^A316T^ in response to 11 mM glucose (G11) alone or supplemented with 100 nM exendin-4 (Ex-4); *n* = 4. (**H**) Insulin secretion fold changes to G11 calculated from data in (G). Data are mean ± SEM; *p<0.05; **p<0.01; ***p< 0.001; ****p<0.0001; ns, not significant by paired t-tests or one-way ANOVA with Sidak’s post-hoc tests.

### Modulation of GLP-1R trafficking by the A316T variant

Having characterised the functional effects of the A316T variant *in vivo* and *ex vivo*, we next explored the molecular mechanisms underlying the observed changes in GLP-1R cell surface expression and signalling. To do so, we employed an *in vitro* INS-1 832/3 rat β-cell model where endogenous GLP-1R expression is deleted by CRISPR/Cas9 (*46*), to generate stable multiclonal cell lines expressing SNAP/FLAG-tagged WT or A316T hGLP-1R. Similarly to our previous findings, SNAP/FLAG-*hGlp1r*^A316T^ cells showed reduced receptor surface expression *versus* its WT counterpart (Fig. 6A). Moreover, assessment of SNAP/FLAG-*hGlp1r*^wt^ *versus* ^A316T^ expression by Western blotting, or with the cell-permeable SNAP-tag probe BG-OG (*47*), revealed significantly reduced total levels of the A316T variant compared to the WT receptor (Fig. 6B and C), suggesting increased basal degradation or reduced protein synthesis. To clarify the mechanism underlying this phenotype, we used bafilomycin A1, an established vacuolar-type H^+^-ATPase inhibitor that prevents lysosomal degradation (*48*). Bafilomycin A1 exposure resulted in complete recovery of surface expression levels in INS-1 832/3 *Glp1r*^-/-^ SNAP/FLAG-*hGlp1r*^A316T^ cells, now comparable to that of WT controls (Fig. 6D). The same effect was observed using *hGlp1r*^-/-^ islets transduced with SNAP/FLAG-*hGlp1r*^WT^ *versus* ^A316T^ adenoviruses (Fig. 6E), or in islets purified from *hGlp1r*^+/+^ *versus* ^A316T/A316T^ mice (Fig. 6F), indicating increased basal turnover resulting in enhanced lysosomal degradation as the cause for the decrease in surface and total levels of the GLP-1R A316T variant. Next, we assessed GLP-1R trafficking behaviours in response to GLP-1RA stimulation in INS-1 832/3 *Glp1r*^-/-^ SNAP/FLAG-*hGlp1r*^WT^ and ^A316T^ cells by high-content microscopy. A316T GLP-1R internalisation was similar to WT in response to exendin-4 and exendin-F1 but was significantly increased with the endogenous agonist GLP-1, reflecting potentially faster kinetics at earlier time-points (Fig. 6G, fig. S7A and B). A316T receptor recycling to the plasma membrane was again not different to WT with exendin-4 or exendin-F1 but was significantly reduced for GLP-1 (Fig. 6H, fig. S7C and D). Finally, receptor degradation in response to agonist stimulation was not significantly different between WT and A316T for any of the agonists tested (Fig. 6I, fig. S7E and F). Overall, the A316T variant receptor appears to show a specific trafficking pattern in response to GLP-1, but very similar trafficking behaviours to the WT receptor with exendin-4-derived agonists, perhaps reflecting differences in its specific response to the endogenous agonist which might account for the improved oral glucose tolerance (which depends on endogenous GLP-1) but reduced pharmacological GLP-1RA responses observed in these mice.

**Fig. 6.**
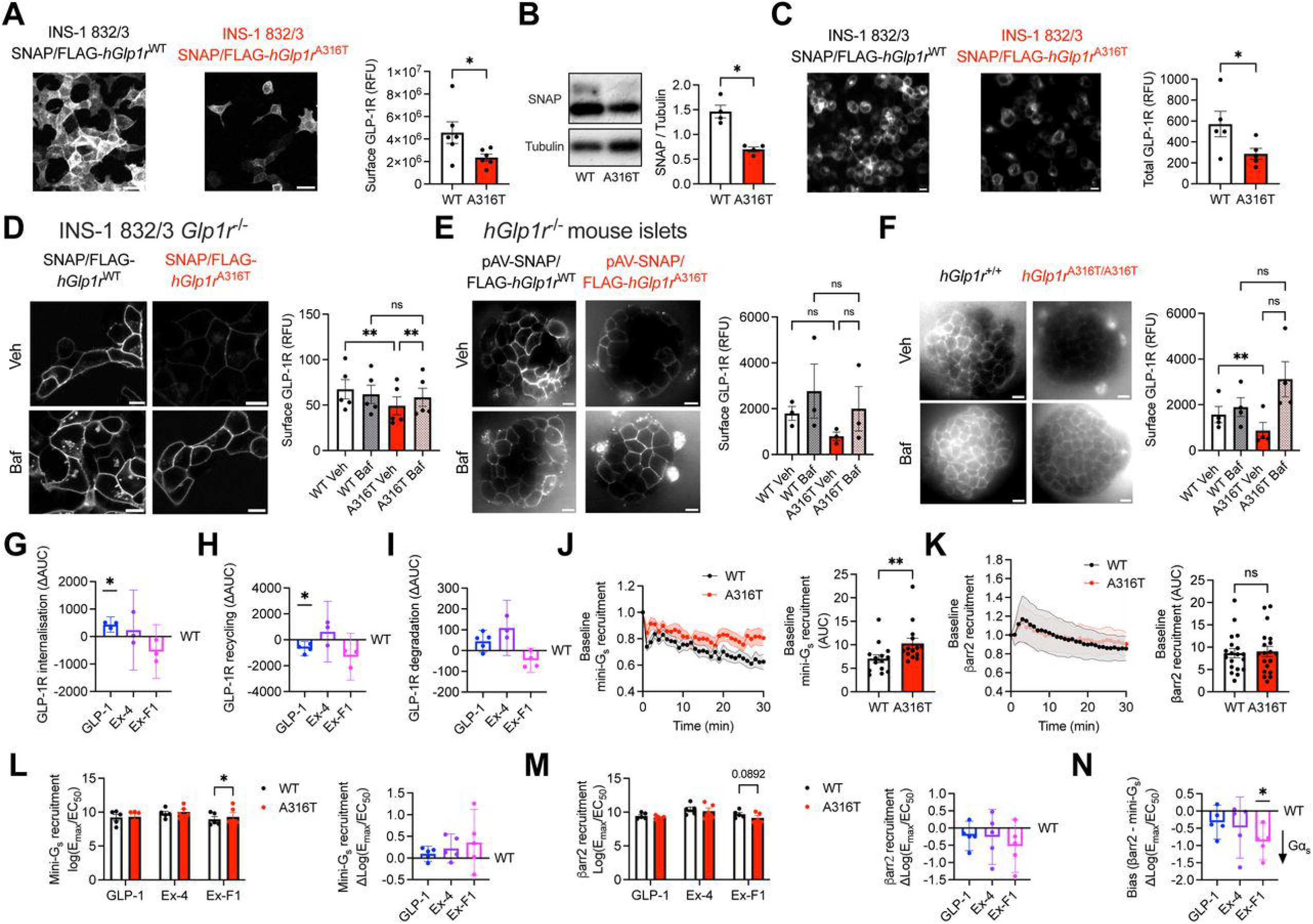
Changes in basal turnover and functional characterisation of INS-1 832/3 *hGlp1r*^-/-^SNAP/FLAG-*hGlp1r*^WT^ *versus* ^A316T^ sublines. (**A**) Surface GLP-1R expression in rat insulinoma INS-1 832/3 *hGlp1r*^-/-^ SNAP/FLAG-*hGlp1r*^WT^ *versus* ^A316T^ cells labelled with SNAP-Surface Alexa Fluor 647. Representative images and quantification shown; size bars, 10 μm; *n* = 6. (**B**) Total GLP-1R expression in INS-1 832/3 *hGlp1r*^-/-^ SNAP/FLAG-*hGlp1r*^WT^ *versus* ^A316T^ cells, analysed by anti-SNAP tag Western blotting, with tubulin used as loading control; *n* = 4. (**C**) Total GLP-1R expression in INS-1 832/3 *hGlp1r*^-/-^ SNAP/FLAG-*hGlp1r*^WT^ *versus* ^A316T^ cells labelled with membrane-permeable SNAP-tag probe BG-OG. Representative images and quantification shown; size bars, 10 μm; *n* = 5. (**D**) Representative images and quantification of surface GLP-1R expression in INS-1 832/3 *hGlp1r*^-/-^ SNAP/FLAG-*hGlp1r*^WT^ *versus* ^A316T^ cells treated for 2 hours with vehicle (Veh) or 400 μM bafilomycin A1 (Baf) prior to labelling with SNAP-Surface Alexa Fluor 647; size bars, 10 μm; *n* = 5. (**E**) As in (D) for *hGlp1r*^-/-^ mouse islets transduced with pAV-SNAP/FLAG-*hGlp1* ^WT^ or ^A316T^; size bars, 10 μm; *n* = 3. (**F**) As in (D) for *hGlp1r*^+/+^ *versus* ^A316T/A316T^ mouse islets labelled with exendin-9-Cy5 (LUXendin645); size bars, 10 μm; *n* = 4. (**G**) GLP-1R internalisation ΔAUC (SNAP/FLAG-*hGlp1r*^A316T^ minus ^WT^, calculated from data in fig. S7, A and B) in INS-1 832/3 *hGlp1r*^-/-^ SNAP/FLAG-*hGlp1r*^WT^ *versus* ^A316T^ cells stimulated with the indicated agonists; *n* = 3. (**H**) GLP-1R plasma membrane recycling ΔAUC (SNAP/FLAG-*hGlp1r*^A316T^ minus ^WT^, calculated from data in in fig. S7, C and D) in INS-1 832/3 *hGlp1r*^-/-^ SNAP/FLAG-*hGlp1r*^WT^ *versus* ^A316T^ cells stimulated with the indicated agonists; *n* = 3-4. (**I**) GLP-1R degradation ΔAUC (SNAP/FLAG-*hGlp1r*^A316T^ minus ^WT^, calculated from data in fig. S7, E and F) in INS-1 832/3 *hGlp1r*^-/-^SNAP/FLAG-*hGlp1r*^WT^ *versus* ^A316T^ cells stimulated with the indicated agonists; *n* = 3-5. (**J**) Baseline LgBiT-mini-G_s_ recruitment to *hGlp1r*^WT^ or ^A316T^-SmBiT in INS-1 832/3 *hGlp1r*^-/-^ cells; responses over time and AUCs shown; *n* = 14. (**K**) Baseline LgBiT-β-arrestin 2 (βarr2) recruitment to *hGlp1r*^WT^ or ^A316T^-SmBiT in INS-1 832/3 *hGlp1r*^-/-^ cells; responses over time and AUCs shown; *n* = 19. (**L**) LgBiT-mini-G_s_ recruitment to *hGlp1r*^WT^ or ^A316T^-SmBiT in INS-1 832/3 *hGlp1r*^-/-^ cells in response to the indicated agonists; log(E_max_/EC_50_) calculated from data in fig. S7, G and H; SNAP/FLAG-*hGlp1r*^A316T^ minus ^WT^ Δlog(E_max_/EC_50_) also shown; *n* = 5. (**M**) LgBiT-β-arrestin 2 (βarr2) recruitment to *hGlp1r*^WT^ or ^A316T^-SmBiT in INS-1 832/3 *hGlp1r*^-/-^ cells in response to the indicated agonists; log(E_max_/EC_50_) calculated from data in fig. S7, I and J; SNAP/FLAG-*hGlp1r*^A316T^ minus ^WT^ Δlog(E_max_/EC_50_) also shown; *n* = 5. (**N**) LgBiT-β-arrestin 2 minus LgBiT-mini-G_s_ (βarr2 - mini-G_s_) recruitment to *hGlp1r*^A316T^ minus ^WT^-SmBiT, shown as Δlog(E_max_/EC_50_), in INS-1 832/3 *hGlp1r*^-/-^ cells as a measure of signal bias. Data are mean ± SEM except for *hGlp1r*^A316T^ minus ^WT^ responses [ΔAUC or Δlog(E_max_/EC_50_)] which are mean ± 95% confidence interval (CI); *p<0.05; **p<0.01; ns, not significant by paired t-tests, one-way ANOVA with Sidak’s or two-way ANOVA with Tukey’s post-hoc tests.

### A316T-associated changes on GLP-1R coupling to Gα_s_ and β-arrestin 2, and downstream signalling effectors

We next explored changes in the GLP-1R A316T capacity for Gα_s_ and β-arrestin 2 coupling using NanoBiT complementation assays in INS-1 832/3 *Glp1r*^-/-^ SNAP/FLAG-*hGlp1r*^WT^ *versus* ^A316T^ cells. We first analysed mini-G_s_ and β-arrestin 2 recruitment to WT *versus* A316T receptors under vehicle conditions to evaluate changes in baseline coupling activity. The A316T GLP-1R demonstrated significantly increased basal mini-G_s_, with no change in basal β-arrestin 2 recruitment compared to WT receptors (Fig. 6J and K). No significant changes *versus* WT receptors were found for A316T mini-G_s_ recruitment in response to GLP-1 or exendin-4, but we measured increased potency for A316T mini-G_s_ recruitment in response to exendin-F1 (Fig. 6L, fig. S7G and H). Agonist-induced recruitment of β-arrestin 2 was not significantly different for A316T *versus* WT besides a near-significant reduction with exendin-F1 (Fig. 6M, fig. S7I and J). When both effects were combined to calculate β-arrestin 2 over Gα_s_ signalling bias, we found a significant shift towards Gα_s_ recruitment for A316T *versus* WT with exendin-F1 (Fig. 6N). As changes in GLP-1R cAMP outputs are often the result of combined trafficking and Gα_s_ / β-arrestin 2 coupling effects, we next analysed cAMP responses to the different GLP-1RAs in INS-1 832/3 *Glp1r*^-/-^ SNAP/FLAG-*hGlp1r*^WT^ *versus* ^A316T^ cells. While we could not detect significant differences, there was a tendency for increased cAMP in response to GLP-1 and exendin-F1, but reduced responses to exendin-4, and no change in response to semaglutide for A316T *versus* WT receptors in this *in vitro* model (fig. S7, K to P).

We next used a previously described bystander NanoBiT assay, based on the recruitment of active Gα_s_-binding nanobody 37 (Nb37) to plasma membrane *versus* endosomal locations (*49*) to assess the effect of the A316T variant on subcellular receptor activity. Results showed a non-significant tendency for increased basal endosomal over plasma membrane activity of A316T compared to WT receptors (Fig. 7A). Additionally, while agonist-induced endosomal A316T activity showed a tendency to be increased *versus* WT with GLP-1, this was significantly reduced in response to exendin-4, without changes in plasma membrane activity *versus* WT with either agonist (Fig. 7B and C). We also tested potential changes in receptor conformational dynamics, indicative of receptor-agonist binding and activation, using a TR-FRET assay that measures the proximity between the receptor extracellular domain (ECD) and the plasma membrane, as a surrogate for receptor ECD opening. While no changes were detected under basal conditions (Fig. 7D), significantly increased TR-FRET was measured for A316T *versus* WT in response to GLP-1 stimulation (Fig. 7E). We next determined the impact of the A316T mutation on GLP-1R recruitment to plasma membrane lipid nanodomains, or ‘rafts’, a preferential localisation for active GLP-1Rs prior to being internalised into endosomes (*50*). In line with our previous findings of a higher level of constitutive receptor activity, the A316T variant presented with increased basal levels at lipid rafts (Fig. 7F). While there were no changes in raft segregation in response to GLP-1 for A316T *versus* WT, raft localisation was significantly reduced for the A316T variant in response to exendin-4 (Fig. 7G). Given the close association between receptor activation, lipid raft segregation, plasma membrane diffusion, and clustering (*50, 51*), we next investigated changes in these parameters by Raster image correlation spectroscopy (RICS) analysis of INS-1 832/3 *Glp1r*^-/-^ SNAP/FLAG-*hGlp1r*^WT^ *versus* ^A316T^ cells. Results showed significantly reduced plasma membrane diffusion coefficients for A316T *versus* WT GLP-1R under vehicle conditions, an effect normally observed only following agonist stimulation in WT receptors (*52*), as also seen here (Fig. 7, H to J). Finally, we analysed changes in GLP-1R ubiquitination, a key post-translational modification (PTM) linked to changes in GLP-1R signalling and post-endocytic trafficking (*53*). As previously seen (*53*), GLP-1R was basally ubiquitinated under vehicle conditions for both WT and A316T receptors. While exendin-4 stimulation did not affect the level of ubiquitination of the WT receptor, it resulted in significantly increased ubiquitination of the A316T receptor (Fig. 7K), a feature typically associated with receptor targeting to lysosomes for degradation and signal termination (*54*).

**Fig. 7.**
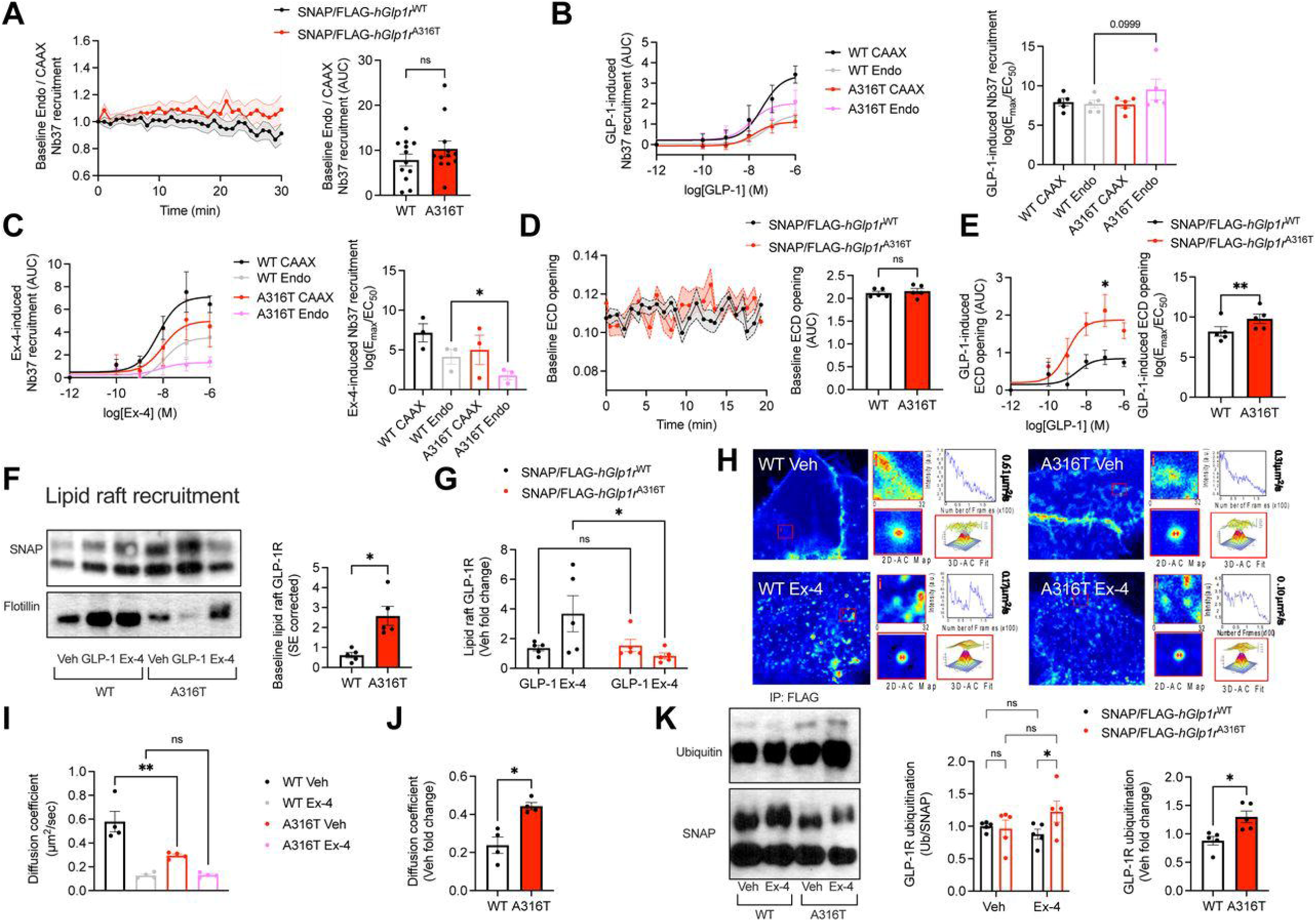
Molecular characterisation of GLP-1R behaviours in INS-1 832/3 *hGlp1r*^-/-^ SNAP/FLAG-*hGlp1r*^WT^ *versus* ^A316T^ sublines. (**A**) Baseline endosomal over plasma membrane GLP-1R activity, measured by SmBiT-Nb37 complementation with Endofin-LgBiT over LgBiT-CAAX under vehicle conditions in INS-1 832/3 *hGlp1r*^-/-^ SNAP/FLAG-*hGlp1r*^WT^ *versus* ^A316T^ cells; AUCs also shown; *n* = 12. (**B**) AUC dose response curves and log(E_max_/EC_50_) of plasma membrane and endosomal GLP-1R activity in response to GLP-1 stimulation, measured by SmBiT-Nb37 complementation with LgBiT-CAAX and Endofin-LgBiT in INS-1 832/3 *hGlp1r*^-/-^ SNAP/FLAG-*hGlp1r*^WT^ *versus* ^A316T^ cells; *n* = 4. (**C**) AUC dose response curves and log(E_max_/EC_50_) of plasma membrane and endosomal GLP-1R activity in response to exendin-4 (Ex-4) stimulation, measured by SmBiT-Nb37 complementation with LgBiT-CAAX and Endofin-LgBiT in INS-1 832/3 *hGlp1r*^-/-^ SNAP/FLAG-*hGlp1r*^WT^ *versus* ^A316T^ cells; *n* = 3. (**D**) Baseline GLP-1R ECD opening, measured by SNAP-Lumi4-Tb - NR12A TR-FRET over time under vehicle conditions in INS-1 832/3 *hGlp1r*^-/-^ SNAP/FLAG-*hGlp1r*^WT^ *versus* ^A316T^ cells; AUCs also shown; *n* = 5. (**E**) AUC dose response curves and log(E_max_/EC_50_) of GLP-1R ECD opening in response to GLP-1 stimulation, measured by SNAP-Lumi4-Tb - NR12A TR-FRET increases over time in INS-1 832/3 *hGlp1r*^-/-^ SNAP/FLAG-*hGlp1r*^WT^ *versus* ^A316T^ cells; *n* = 5. (**F**) Representative blots showing SNAP/FLAG-*hGlp1r* recruitment to DRM fractions (or lipid rafts) in INS-1 832/3 *hGlp1r*^-/-^ SNAP/FLAG-*hGlp1r*^WT^ *versus* ^A316T^ cells following stimulation for 2 minutes with the indicated agonist, with Flotillin used as DRM loading control; quantification of surface expression (SE)-corrected baseline SNAP/FLAG-*hGlp1r*^WT^ *versus* ^A316T^ DRM levels under vehicle conditions also shown; *n* = 5. (**G**) SNAP/FLAG-*hGlp1r*^WT^ *versus* ^A316T^ DRM recruitment fold changes to vehicle (Veh), quantified from (F); *n* = 5. (**H**) Representative images from RICS analysis of GLP-1R WT and A316T plasma membrane lateral diffusion in INS-1 832/3 *hGlp1r*^-/-^ SNAP/FLAG-*hGlp1r*^WT^ *versus* ^A316T^ cells labelled with SNAP-Surface Alexa Fluor 647 and subsequently stimulated with vehicle (Veh) or 100 nM Ex-4. (**I** and **J**) GLP-1R WT *versus* A316T average RICS diffusion coefficients (I) and vehicle (Veh) fold changes (J); *n* = 4. (**K**) Representative blots showing SNAP/FLAG-*hGlp1r* ubiquitination following anti-FLAG immunoprecipitation in INS-1 832/3 *hGlp1r*^-/-^ SNAP/FLAG-*hGlp1r*^WT^ *versus* ^A316T^ cells under vehicle (Veh) or 100 nM Ex-4 stimulation for 10 minutes; quantification of GLP-1R ubiquitination and vehicle (Veh) fold changes also shown; *n* = 5.

Next, to assess the global signalling effect of the GLP-1R A316T variant at a whole cell level, we performed a mass spectrometry-based interactome analysis of vehicle *versus* agonist-(GLP-1 or exendin-4) stimulated A316T GLP-1R using INS-1 832/3 *Glp1r*^-/-^ SNAP/FLAG-*hGlp1r*^A316T^ in parallel with SNAP/FLAG-*hGlp1r*^wt^ cells [WT data previously shown in (*55*)]. Results normalised to isolated receptor amounts per immunoprecipitate revealed increased GLP-1R-interactor binding across the whole interactome for A316T compared to WT under vehicle conditions, a measure indicative of the level of basal activation inherent to each receptor type (fig. S8). Stimulation of A316T with either GLP-1 or exendin-4 led to further increases in GLP-1R-protein binding for most interactors, but we identified some factors exhibiting reduced binding to active *versus* inactive A316T receptors, reversing the normal pattern of increased binding to these factors following GLP-1RA stimulation of WT receptors. These included, notably, the ER membrane contact site (MCS) tether VAP-B and the A-kinase anchoring protein (AKAP) SPHKAP, two GLP-1R binding partners for which we have recently identified a critical role in the control of mitochondrial function by the GLP-1R (*55*).

### Structural determination of the GLP-1R A316T variant

To elucidate structural changes associated with the A316T variant receptor, a cryo-EM structure of the GLP-1-bound GLP-1R A316T in complex with dominant negative (DN)-Gα_s_ was determined at 3.3 Å resolution (Fig. 8A and B). In general, the complex was well resolved and amino acid side chains modelled for almost the entire structure. The α-helical domain of DN-Gα_s_, the flexible ECD, the extracellular loop 3 (ECL3) and the intracellular loop 3 (ICL3) of GLP-1R A316T, as well as the final two C-terminal residues of GLP-1, were less well resolved in the cryo-EM map, in line with the intrinsic flexibility of these regions. The overall positioning of the A316T transmembrane domains (TMDs) was highly similar to that of the WT receptor (PDB 6X18, Fig. 8C), and the binding arrangement of GLP-1 at the backbone level essentially identical to that previously reported for WT GLP-1R by high-resolution cryo-EM (*56*) (Fig. 8D). In the WT GLP-1R structure, the C-terminus of GLP-1 interacts with the ECD and ECL1/2 of GLP-1R, forming extensive contacts with transmembrane (TM) helices 1, 2, 3 and 5, enabling the peptide N-terminal residues to engage deep into the orthosteric site with a conserved class B1 GPCR central polar network, including a salt bridge with R190^2.60^. A network of structural waters connects the central parts of TM3, 5 and 6 below the peptide binding site, forming hydrogen bonds (H-bonds) with the receptor backbone, including with the highly conserved G361^6.50^ at the P^6.47^xxG^6.50^ motif (*56*), crucial for receptor activation and transducer coupling. Additionally, the backbone carbonyl of Y241^3.44^, part of the conserved central polar network, directly interfaces through a H-bond with a structural water located between TM3 and TM5, within H-bond distance of the backbone carbonyl of A316^5.46^, contributing to increase the interhelical packing and stiffness. In the T316 variant, however, this backbone carbonyl sits ∼1.5 Å away from the structural water and has a shallower angle compared to A316^5.46^, making it poorly favourable for H-bonding (Fig. 8E). Intriguingly, when modelled as the ‘upwards’ rotamer, the hydroxyl group of T316^5.46^ can H-bond with the oxygen of this structural water, likely precluding the known interactions discussed above. The interface between the GLP-1R and Gα_s_, highly conserved with other class B1 GPCRs (*57*), is preserved in the A316T GLP-1R, with interactions between the variant receptor and the α5 helix of Gα_s_ very similar to those found in WT GLP-1R agonist-bound structures (*56*) (Fig. 8F and G). Higher resolution structures of agonist-bound GLP-1R in complex with Gα_s_ have uncovered important structural waters in this region, including one acting as a key bridge between E247^3.50^, in the highly conserved HETX motif, and Y391^Gαs^. At this resolution, these waters were not present in the density map of GLP-1R A316T, with E247^3.50^ poorly defined compared to neighbouring residues of similar size, which is not unusual for an acidic residue in single-particle cryo-EM at 3.3 Å resolution. N338^ICL3^ was also poorly resolved, indicating that this residue is comparatively mobile (Fig. 8H, left graph). The position of R176^2.46^ is rotated towards the α5 helix of Gα_s_ by 2 Å in GLP-1R A316T compared to WT and, independently, H387 is rotated slightly from its position in the WT GLP-1R structure (Fig. 8H, right graph). Overall, we observed subtle differences that may contribute to the altered Gα_s_ recruitment sometimes observed with this mutant (*24, 27*). In summary, the static model built into the consensus density map of the GLP-1:GLP-1R(A316T):DN-Gα_s_:Nb35 complex, while largely similar to the WT receptor, suggests changes to key water-mediated interactions between TM5 and its neighbouring helices that could alter the dynamics of the TMD. Additionally, subtle changes to interactions between the intracellular face of the receptor and the α5 helix of DN-Gα_s_ may correlate with the pharmacological profile of the A316T variant.

**Fig. 8.**
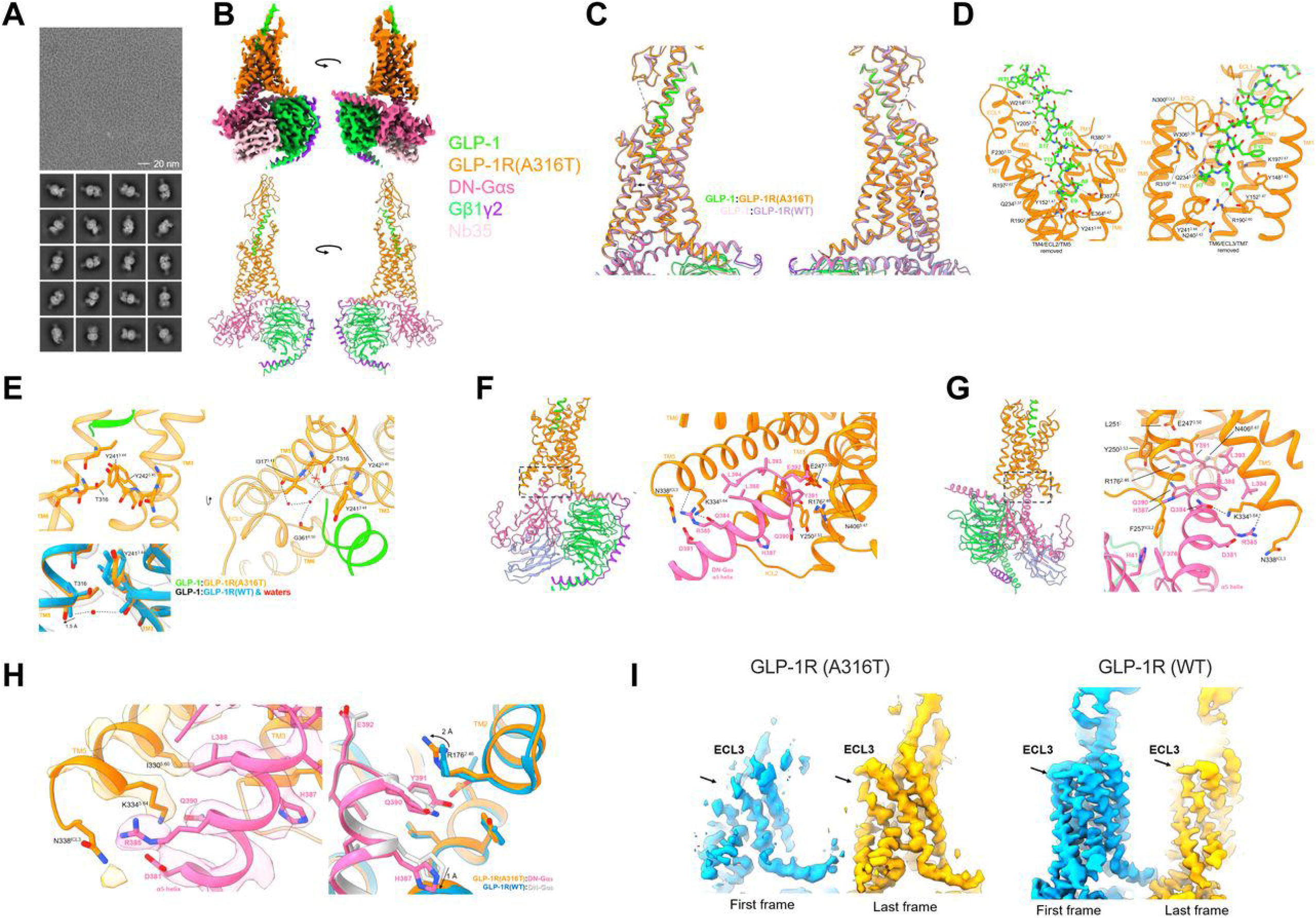
Cryo-EM structure of GLP-1:GLP-1R A316T:DN-Gα_s_:Nb35. (**A**) Representative micrograph from 200 kV cryo-EM imaging of the complex (top), with representative 2D class averages of particles used in the consensus refinements (bottom). (**B**) Non-uniform refined density map used as the primary source for modelling (top), with atomic model built into the density map, including backbone trace of ECD (bottom). (**C**) Transmembrane domains, rendered as cartoons, of GLP-1R A316T (orange) and WT (lavender) overlaid (left), and 140° rotation (right); black arrows indicate the position of residue 316. (**D**) Binding arrangement of residues 7-31 of GLP-1 (green) with GLP-1R A316T (orange), facing inwards from TM5 (left), and from TM6 (right); dashed lines indicate H-bonds predicted by ChimeraX 1.5. (**E**) top left, Positioning of T316^5.46^ and Y241^3.44^ below the binding site for GLP-1 (green); (**E**) bottom left, comparison between local arrangement of residues T/A316^5.46^ and Y241^3.44^ in the A316T variant (orange) and the WT GLP-1R (azure blue; PDB 6X18) (bottom left); a structural water (red) from the Å WT structure has H-bond interactions (dashed lines) with the backbone of these residues; black arrow indicates movement of T316 carbonyl away from A316^5.46^ position; the 2 Å zoned density map (3.3 Å resolution) of the GLP-1R A316T structure is represented as a transparent grey surface; (**E**) right, Intracellular-facing view of the receptor core, showing the positioning of the TM3 - TM5 - TM6 structural water network (red) from PDB 6X18. A red cross indicates a potential H-bond lost for T316. TM1, 2, and 7 have been removed from each panel for clarity. (**F**) Key interactions between the GLP-1R A316T intracellular cavity (orange) and the α5 helix of DN-Gα_s_ (pink) with ‘forward’ view indicated by left cartoon. H-bonds, determined by ChimeraX 1.5, are dashed lines. (**G**) Alternative view of GLP-1R A316T – DN-Gα_s_ α5 helix interactions, as indicated by left cartoon. (**H**) left, Cryo-EM density map used to model interactions, depicted as coloured transparent silhouettes. There is no density for N338 ICL3, while other nearby side chains are well-resolved; (**H**) right, Comparison between GLP-1R A316T - α5 helix (orange and pink) and GLP-1R WT - α5 helix (blue and white) interactions. (**I**) left, Close-up of first and last frames from principal component 3 after 3DVA of GLP-1R A316T, with distant density clipped for clarity; (**I**) right, Principal component 3 from 3DVA for GLP-1R WT; raw data from (*111*); ECL3 is indicated by bold arrows.

To explore these observations further, 3D variability analysis (3DVA) was performed in cryoSPARC 3.2, a powerful tool for examining conformational heterogeneity in cryo-EM datasets (*58*). These data were analysed by a principal component analysis (PCA) model searching for three principal components with a tight mask around the receptor and G proteins to exclude variability from the detergent micelle. Previous analyses of GLP-1R cryo-EM structures with peptide ligands have demonstrated three major modes of motion: a coordinated twisting movement of the receptor TMD, a large oscillation of the peptide C-terminus and ECD, and a twisting/rocking of the receptor and G protein (*56*). A striking increase in variability for ECL3 in GLP-1R A316T compared to the WT receptor was observed in all three principal components and was preserved at low contour. This motion was particularly evident in principal component 3 and was accompanied by more extreme movements in TM6 and the N-terminal half of the peptide (Fig. 8I). Compared to the 3DVA of the GLP-1R WT, GLP-1R (A316T) TM6 - ECL3 - TM7 underwent a larger motion down and away from the receptor core (Fig. 8I).

### Molecular dynamics (MD) simulations of WT versus A316T GLP-1R: Impact of the A316T SNP on receptor dynamics, structural water molecules, and H-bond interactions

To retrieve atomistic insights about the effect of the A316T substitution on the structure and dynamics of the GLP-1R, A316T and WT GLP-1Rs were simulated in three different configurations: i) the isolated TMD in the absence of any agonist; ii) in complex with the endogenous agonist GLP-1 [i.e., the new A316T cryo-EM structure presented here compared to PDB 6X18 (fig. S9A)]; and iii) in complex with exendin-F1 and Gα_s_ (i.e., the WT cryo-EM structure presented in PDB 9C0K and a modelled A316T structure based on the former). In the TMD, the side chain of T316^5.46^ (A316T) formed a H-bond with Y242^3.45^, an interaction which was not present in the WT GLP-1R, where Y242^3.45^ interacted with the P312^5.42^ backbone instead (Fig. 9A, fig. S9B). This A316T-specific Y242^3.45^ - T316^5.46^ H-bond (table S1) was very persistent in the isolated TMD (85%), while it was less persistent but still present for more than one-third of the simulations in A316T GLP-1R in complex with either GLP-1 or with exendin-F1 and Gα_s_ (41.8% and 37.7%, respectively). This suggests that the engagement of Y242^3.45^ with the hydrophilic side chain of T316^5.46^ instead of the P312^5.42^ backbone occurs in a ligand-dependent manner. The reason for the Y242^3.45^ - T316^5.46^ H-bond strength in the isolated TMD of A316T lies in the rotameric state of T316^5.46^ adopting a configuration prone to accept the H-bond from Y242^3.45^ and donating the hydrogen to the P312^5.42^ backbone (State 1 in Fig. 9A, table S1, movie S1). In A316T GLP-1R in complex with exendin-F1 and Gα_s_, the persistence of the Y242^3.45^ - T316^5.46^ H-bond was similar to that in A316T GLP-1R in complex with GLP-1, but the Y242^3.45^ - P312^5.42^ and T316^5.46^ - P312^5.42^ interactions were similar to those found in the isolated A316T GLP-1R TMD, suggesting different local GLP-1R A316T dynamics with exendin-F1 or with the presence of bound Gα_s_. For the A316T GLP-1R in complex with GLP-1, the T316^5.46^ side chain flipped between rotameric State 1 and another conformation able to form a H-bond with the backbone of I313^5.43^ and a water bridge with the Y241^3.44^ backbone (State 2 in Fig. 9B, tables S1 and S2). This H-bond reshuffling is consistent with our previous MD simulations of oxyntomodulin and Gα_s_-complexed WT *versus* A316T GLP-1R (*24*) where the Y242^3.45^ - P312^5.42^ H-bond in the WT (88.9% occupancy) was replaced by the Y242^3.45^ - T316^5.46^ H-bond in A316T (62.3% occupancy) due to a consistent rotameric State 1 throughout the simulations, corroborating an allosteric effect of the agonist in the T316^5.46^ dynamics and, in turn, Y242^3.45^ - T316^5.46^ H-bond. Two clear rotameric states for T316^5.46^ were also observed in our consensus cryo-EM maps, able to accommodate both an “upward” and a “downward” side-chain orientation (Fig. 8), supporting the existence of both rotameric states of A316T occupied with similar frequencies in the presence of GLP-1.

**Fig. 9.**
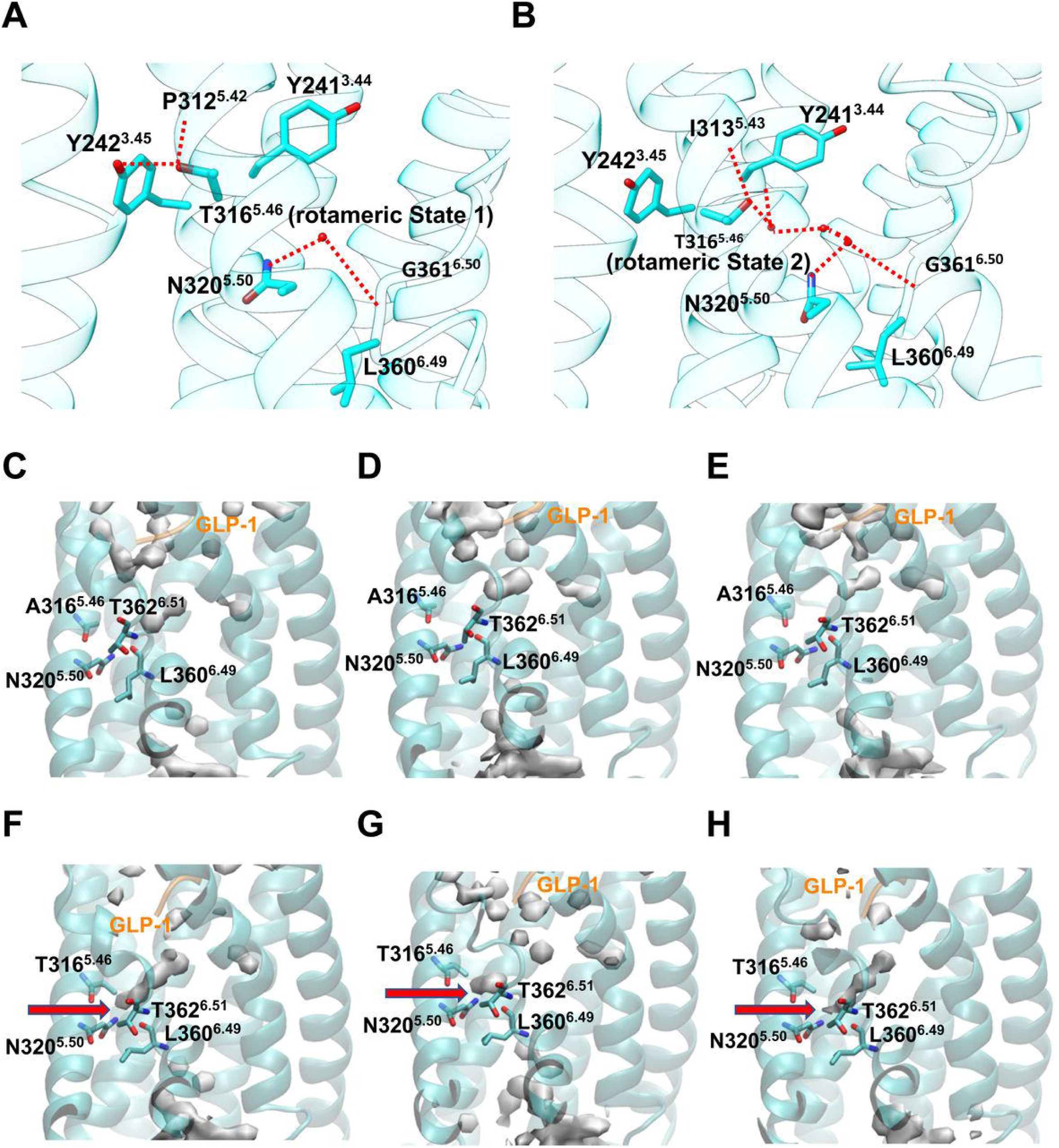
Divergent H-bond and solvation patterns in WT *versus* A316T GLP-1R. (**A**) T316^5.46^ rotameric State 1 forms a stable H-bond with Y242^3.45^. (**B**) T316^5.46^ rotameric State 2 is involved in water-mediated interactions with the polar core of the receptor. N320^5.50^ forms stable water bridges with TM6 kink (L360^6.49^ and G261^6.50^) in A316T. (**C** to **E**) volumetric map (isovalue 0.3) of water molecules within the TMD of WT GLP-1R (cyan ribbon) in complex only with the endogenous agonist GLP-1 (orange ribbon) from 3 independent MD simulation replicas. (**F** to **H**) Volumetric map (isovalue 0.3) of water molecules within the TMD of A316T (cyan ribbon) in complex only with the endogenous agonist GLP-1 (orange ribbon) from 3 independent MD simulation replicas. The red arrows indicate the position of stable water molecules interacting with N320^5.50^ and the TM6 kink.

T316^5.46^ (A316T) in rotameric State 2 weakly interacted with the conserved N320^5.50^ through a water bridge (Fig. 9B, table S2, movie S1). This was also the case for GLP-1R (WT) bound to GLP-1, but here rotameric State 2 favoured a higher persistence of H-bond interactions with a water network lining T316^5.46^ and N320^5,50^. A detailed analysis of the interaction involving N320^5.50^ (table S3, movie S1) reveals a stable direct H-bond with the backbone of L360^6.49^ at the TM6 kink level in the three WT GLP-1R simulations [occupancy 68.5% (isolated TMD), 79.6% (with GLP-1), and 65.4% (with exendin-F1 and Gα_s_)]. This N320^5.50^ - L360^6.49^ H-bond is weaker in A316T when in complex with the endogenous agonist GLP-1 (35.6%) and for the isolated TMD (30.6%), but only slightly weaker in the case of A316T in complex with exendin-F1 and Gα_s_ (54.8%) (movie S1), in line with different local dynamics imprinted by exendin-F1. The diminished H-bond strength is counterbalanced by an increase in water-mediated interactions between N320^5.50^ and L360^6.49^, and between N320^5.50^ and G361^6.50^, in A316T in complex with GLP-1 (12.2% in WT *versus* 26.8% in A316T) and the isolated TMD (19.5% in WT *versus* 49.3% in A316T). Such water-mediated interactions involving N320^5.50^ in A316T were due to a deeper water molecule permeation in the polar core of the receptor compared to WT GLP-1R (Fig. 9, C to H), stabilised by a concerted effect of the polar side chain of T316^5.46^ and the presence of the agonist. Indeed, a significant increase of up to 15% in the water molecule residence time was observed in A316T in complex with GLP-1 or with exendin-F1 and Gα_s_ (table S4) but not in the isolated TMD, where the water was exchanged with bulk solvent instead of being stabilised into structural water molecules, as occurring in the agonist-complexed A316T configurations.

To retrieve further insights into the allosteric effect of the persistent Y242^3.45^ - T316^5.46^ H-bond formed by T316^5.46^ in rotameric State 1, we further analysed the MD simulations involving the isolated TMD of GLP-1R. We reasoned that the GLP-1R structure other than the TMD (i.e. ECD and ECL1), or the binders GLP-1 and Gα_s_, can either directly (GLP-1, Gα_s_) or allosterically (ECD, ECL1) influence the possible conformations that the TMD adopts because of the A316T mutation, *de facto* shielding any subtle allosteric effects. The Y242^3.45^ - T316^5.46^ H-bond in A316T moved TM3 and TM5 away from each other, as shown by the increased backbone distance between the closely downstream residues L245^3.48^ - V319^5.49^ (fig. S9B and C). This is consistent with our cryo-EM maps of A316T in complex with GLP-1 and Gα_s_, where the bulkier T316^5.46^ side chain sterically displaces the T316^5.46^ backbone carbonyl ∼1.5 Å away from the interhelical space compared to the WT A316^5.46^ (Fig. 8E). Structural contractions or expansions at the extracellular or intracellular side of a GPCR can allosterically influence the receptor dynamics at the opposite side (*59*). Consistent with this GPCR structural effect, in A316T GLP-1R the TM3-TM5 intracellular distance, measured at the L251^3.54^ - V327^5.57^ level, was ∼1 Å shorter compared to that of the WT GLP-1R (fig. S9B and C), favoured by a more persistent H-bond between Y252^3.55^ and R326^5.56^ (35.7% occupancy in A316T *versus* 11.4% in WT; fig. S9B). In concert with the TM3 - TM5 intracellular contraction, the TM6 outwards opening, a structural hallmark of GPCR activation to accommodate the Gα_s_ protein, was about 2.5 Å wider in A316T than in WT (27.65 ± 2.02 Å *versus* 25.12 ± 1.80 Å, measured between the Cα atoms of C347^6.36^ and E408^7.63^, fig. S9D). As a reference, in the static cryo-EM structure of A316T, and in PDB 6X18, the same distance is 22.94 Å and 22.08 Å, respectively. The wider TM6 opening in the isolated TMD of A316T was however not observed when A316T was in complex with GLP-1, suggesting that the agonist can modulate the TM3 - TM5 distance by influencing the rotameric state of T316 and in turn the formation of the Y242^3.45^ - T316^5.46^ H-bond.

## Discussion

GPCRs are well known to harbour SNPs, leading to phenotypic variability amongst individuals and potentially impacting pharmacological responses and disease traits (*60*). Despite the importance of GLP-1R as a major T2D and obesity drug target, and the known variability of responses to GLP-1RAs amongst individuals taking this drug class (*61*), there is surprisingly little research on the functional consequences of *Glp1r* genetic variants associated with glycaemic traits and human metabolic diseases, particularly *in vivo*. Although most *Glp1r* variants are low-frequency mutations, understanding their phenotypes and associated pathophysiological changes in humans is valuable as a basis for the development of personalised therapeutic strategies, especially when considering the large number of individuals affected by diseases than can be treated with pharmacological GLP-1RAs, as well as to model conformational effects to be mimicked or avoided when developing novel GLP-1RAs. Our study has attempted to bridge this gap by employing a CRISPR/Cas9-based approach to generate a humanised knock-in mouse line bearing the A316T mutation in the human *Glp1r* locus in conjunction with *in silico*, *ex vivo*, and *in vitro* approaches to understand the association of the rs10305492 G>A variant with T2D protection. Although this missense variant has been previously studied *in vitro*, analyses using HEK293T, CHO or INS-1 cell lines (*24, 27, 29, 62*) have not provided a comprehensive mechanistic explanation for its functional effects, and no data is available in primary tissues or *in vivo*. Phenotypes such as potential changes in islet cytoarchitecture, as unveiled here, or specific variant effects under metabolic stress or T2D conditions cannot be modelled *in vitro*, and GWAS studies might not reflect genetic variant effects in individuals with prevalent disease as this is often tested in individuals of “normal” physiology (*63*).

Here, our *in vivo* A316T model exhibits improved oral glucose tolerance under both chow and HFHS diet conditions, suggesting enhanced sensitivity to endogenous GLP-1, a phenotype that has not previously been identified in GWAS studies and that correlates with a specific GLP-1R A316T trafficking profile in response to this agonist *in vitro*. A316T-expressing mice also present with significantly lower fasting glucose levels in the chow-fed state, this time in agreement with GWAS reported effects (*22–24*). Interestingly, the variant exhibits slower weight gain in diet-induced obese mice, suggesting a protective effect of A316T for the development of obesity, detectable here despite the reduced GLP-1R expression of our humanised *Glp1r* mouse model, which is likely to mask the overall effect of GLP-1RAs in appetite control. We therefore expect that the effect of the A316T variant in reducing weight gain in organisms expressing normal levels of GLP-1R might be even greater, and suggest that this phenotype warrants additional investigations outside of the scope of the present study.

Despite the positive effect of the A316T variant on glucose homeostasis and endogenous GLP-1 responses, it also presented with blunted responses to several pharmacological GLP-1RAs. Not all agonists were equally affected, however, with exendin-4 and semaglutide responses being particularly hampered, to the point of causing an apparent increase in blood glucose levels *versus* vehicle exposure at prolonged time-points, suggesting a potentially detrimental impact on glucoregulation under those conditions, while exendin-F1, a Gα_s_-biased analogue of exendin-4, retained some beneficial effect 6-hours post-agonist administration despite an initial acute loss in chow-fed homozygous A316T *versus* WT mice. The *in vivo* reduction in blood glucose levels under vehicle administration correlate with increases in plasma insulin in both lean and obese A316T mice under these unstimulated conditions, likely due to the increased constitutive activity of GLP-1R A316T, which mimics agonist-induced effects in the pancreas, and either precludes further increases in insulin secretion by exendin-4 or triggers excessive desensitisation following exendin-4 exposure, resulting in the loss of downstream signalling responses to this GLP-1RA. We have also established the clinical relevance of our observations, as heterozygous *hGLP1r*^A316T/+^ mice, the most likely presentation of this variant due to its low frequency in human populations, presented with blunted responses to semaglutide in HFHS-fed conditions.

In addition to the *in vivo* study, we have also performed *ex vivo* analyses of the effects of the A316T variant on islet gene expression, morphology, and α-*versus β*-cell mass. While effects were relatively mild in islets from chow-fed mice, we observed increased islet diameters and a striking accumulation of α-cells in *hGLP1r*^A316T/A316T^ islets following HFHS feeding. Under this metabolic stress condition, α-cells were unusually distributed in these islets across the islet core instead of being restricted only to the islet mantle. Furthermore, we observed the existence of bi-hormonal cells expressing both insulin and glucagon, the presence of which has been sometimes associated with human islet exposure to GLP1RA treatment (*64, 65*). High-fat diet and metabolic stress can also induce β-cell-like features in a subset of α-cells, as well as increase α-cell-derived GLP-1 secretion (*66, 67*); however, this is, to our knowledge, the first description of increased levels of α-cells, suggestive of a β-to-α cell conversion, under a HFHS diet with chronically overactive GLP-1Rs. While the reasons behind our observation remain unclear, it is possible that a compensatory mechanism might take place in HFHS diet-exposed mice expressing A316T GLP-1R resulting in an abnormal accumulation of both α- and bi-hormonal cells; such effect might potentially modify intra-islet paracrine signalling, for example by increased local stimulation of β-cell GLP-1Rs by glucagon under low glucose conditions, contributing to the observed low fasting glucose phenotype and potentially triggering episodes of hypoglycaemia in carriers of the A316T variant. Despite these changes in islet morphology and endocrine cell type, we only observed increased expression of the β-cell enriched gene *MafA*, with only a tendency for increased expression of both *Ins1* and *Gcg* in HFHS diet-fed *hGLP1r*^A316T/A316T^ mouse islets. Our experiment however does not allow the determination of cell type-specific transcriptional profiles and might potentially have been underpowered. Further determination of transcriptional signatures from cells in these islets would be required to clarify the endocrine cell identify changes observed in the islets of HFHS-fed A316T mice.

To further understand the mechanisms underlying the changes in glucoregulation observed with the GLP-1R A316T variant, we have assessed the levels of receptor cell surface expression, cAMP production and insulin secretion in *hGLP1r*^+/+^ *versus*^A316T/A316T^ islets. In agreement with previous *in vitro* observations (*29, 68*), we detected a significant reduction in cell surface expression of the A316T variant, both *ex vivo* and *in vivo*, by injecting the LUXendin645 probe into living mice. This reduced surface expression of the A316T receptor has been corroborated throughout this study in all systems tested, including human EndoC-βH3 *hGlp1r* ^-/-^ cells, human islets and islets from *hGlp1r* ^-/-^ mice transduced with adenoviral vectors expressing pAV-SNAP/FLAG-*hGlp1r*^WT^ or ^A316T^, as well as in INS-1 832/3 *hGlp1r*^-/-^ cells stably expressing the WT or A316T human GLP-1R variant, despite receptor over-expression in some of these models. We have also established here that this effect is due to an increased rate of receptor turnover, involving increased rates of endocytosis and lysosomal targeting of the GLP-1R A316T variant under basal conditions, in keeping with a higher degree of constitutive activity conferred by the A316T mutation, rather than involving changes in transcription or biosynthetic delivery to the plasma membrane. Further indications of the increased basal activity of GLP-1R A316T obtained in this study include increased levels of basal cAMP, basal intracellular calcium, and mini-Gs recruitment under vehicle conditions, as well as increased constitutive receptor recruitment to lipid rafts, and reduced basal levels of receptor plasma membrane diffusion, as well as an overall increase in binding to receptor protein interactors under unstimulated conditions, as shown in the MS interactome analysis.

In this study, we have also measured enhanced levels of glucose-stimulated insulin secretion for the A316T variant in the various *ex vivo* and *in vitro* human and mouse β-cell models tested, which, in agreement with our *in vivo* results, did not translate to further increases in insulin secretion in response to GLP-1RAs, blunting the effect of these compounds over vehicle alone responses. Prolonged exendin-4 responses were particularly affected, a result that correlates with the *in vivo* loss of glucoregulatory benefit for this agonist 6 hours post-agonist administration. This effect further correlates with other indicators of poor signalling responses or increased desensitisation in response to exendin-4 for the A316T variant, including significantly reduced receptor segregation to lipid rafts and reduced endosomal activity, increased receptor ubiquitination, and a tendency for reduced exendin-4-induced cAMP generation *versus* WT GLP-1Rs.

Conversely, cAMP generation or intracellular Ca^2+^ mobilisation in response to the Gα_s_-biased analogue exendin-F1 were improved in A316T compared to WT GLP-1R-expressing islets, which correlates with a milder impairment in glucoregulation, particularly at prolonged time points, with this GLP-1RA. We hypothesise that this effect might be linked to the favourable signalling profile of this Gα_s_-biased GLP-1RA, which furthermore presents with an extra bias towards Gα_s_ v*ersus* β-arrestin 2 at the A316T compared to the WT receptor, as shown in the present study, further reducing its propensity to trigger increased desensitisation of the variant receptor. This characteristic would enable exendin-F1 to partly counteract the accelerated desensitisation of the A316T variant receptor, a feature that instead would be exacerbated by a fast-desensitising agonist such as exendin-4 (*32*). Exendin-F1 might also enable the A316T receptor to adopt a more favourable conformation for signal transduction, as highlighted here by the different local dynamics imprinted by this agonist in our MD simulations.

Another interesting observation from our study is the consistently improved oral glucose tolerance present in mice expressing A316T *versus* WT receptors, observed here in mice fed either chow or HFHS diets, the latter despite no significant reductions in fasting glucose levels for the A316T genotype. While these results might be indirect, they could be the result of enhanced A316T responses to the endogenous incretin GLP-1 compared to pharmacological GLP-1RAs. Some indications in favour of the latter include a higher degree of ECD opening in response to this agonist for the A316T compared to the WT receptor, and a specific trafficking profile of GLP-1-stimulated A316T not present in response to exendin-4-based GLP-1RAs, involving increased internalisation and reduced recycling for the variant *versus* the WT receptor when stimulated with the physiological agonist.

Further insights are provided here by the cryo-EM structure of the GLP-1R A316T variant in complex with Gα_s_ and GLP-1, showing a consensus atomic model similar to that of the WT GLP-1R but with subtle changes to interactions between TM3, 5 and 6. Of note, we were only able to determine the cryo-EM structure of the A316T receptor with a relatively modest (3.3 Å) resolution, a result that we attribute to potentially higher flexibility or dynamics of the A316T variant compared to the WT receptor. It has previously been proposed, using the X-ray crystal structure of exendin-4-bound GLP-1R as a model, that the A316T variant might disrupt h-bonding between N320^5.50^ and E364^6.53^ by the formation of a direct interaction between E364^6.53^ and T316 (*22*). However, the cryo-EM structure of the GLP-1-bound A316T receptor obtained in this study does not exhibit these structural changes, instead suggesting, as supported by our MD simulations, the potential disruption of central polar network residues via several subtle alterations in the interhelical water network bridging TM5 and 6. Additionally, consistent with the mechanism suggested by the MD simulations, the GLP-1-bound GLP-1R A316T cryo-EM structure offers support for the disruption of interactions between Y242^3.45^ and P312^5.42^ in favour of the Y242^3.45^ - T316^5.46^ H-bond, selective for a specific T316^5.46^ rotameric state. Interestingly, 3DVA of the GLP-1R A316T structure also suggest greater mobility in the extracellular loops of the variant receptor, particularly in ECL3, compared to the WT receptor bound to the same agonist (*69, 70*). If this region is truly more dynamic in the A316T variant, it is conceivable that GLP-1 peptide cycling (disengagement and re-engagement) with the receptor could be faster, leading to more signalling events per unit time than for the WT GLP-1R. These observations are subject to certain limitations related to the abovementioned modest resolution of the A316T variant structure, which allows for confident placement of the receptor TMD side chains but is below the resolution required to place structural waters implicated in the proposed mechanism of constitutive activity for this variant. To overcome this limitation, we have employed MD simulations which are a useful tool to study water molecules within biological systems with good reliability (*71*). Here, they aided in predicting a water network involved in the communication between TM5 and TM6 of A316T, possibly contributing to the unique pharmacology of A316T. Furthermore, the simulations also suggested an allosteric mechanism triggered by Y242^3.45^ - T316^5.46^ H-bond, transmitted to the intracellular side of the receptor by altering the mobility of TM6 in the apo-state A316T. However, as there are no other published structures of known constitutively active GLP-1R mutants, it is difficult to conclude if the differences in water network interactions and extracellular loop dynamics unveiled in the present structure are sufficient to fully account for the constitutively increased activity of A316T. Our findings additionally suggest that the main source of this enhanced activity is the receptor itself rather than being triggered by ligand binding, despite the different modes of engagement of the variant receptor with specific GLP-1RAs such as exendin-F1. Apo-state structures for both the WT and the A316T receptors would need to be obtained to fully elucidate structural changes in the absence of bound agonist.

Despite these limitations, when taken together, our MD simulations aligned well with the cryo-EM data and are consistent with a concerted mechanism behind the enhanced constitutive A316T activity which is triggered by the bulkier, more polar T316 side chain. The structural model informed by the cryo-EM and MD simulations data for the effect of the Ala to Thr substitution in position 316 (fig. S9E) suggests changes to interactions between TM3 and TM5 triggered at position 316 that are propagated to the intracellular side of GLP-1R. In particular, a stabilising effect of water molecules on the TM6 kink, in concert with the tighter helical packing between TM3 and TM5, aids to the intracellular opening movement of TM6, pivotal for Gα_s_ recruitment and activation. In physiological conditions, this allosteric effect would move the inactive-active A316T equilibrium towards the active state, favouring the constitutive activation of the receptor and the increased basal Gα_s_ coupling observed in our functional experiments.

To conclude, we present here a comprehensive assessment of the effects of the A316T missense mutation in human and murine pancreatic β-cells, primary islets and *in vivo* using a genetically engineered humanised knock-in mouse model, including assessment of both agonist-dependent and - independent responses. We demonstrate *in vivo* increased basal but reduced GLP-1RA-dependent glucoregulatory effects, with parallel changes present for islet insulin secretion, correlating with changes in *ex vivo* and *in vitro* signalling responses. Our study analyses in detail the agonist-dependent and -independent trafficking and signalling profiles of this variant in physiologically relevant β-cell systems, confirming some results from previous *in vitro* and GWAS studies, including a marked GoF phenotype for the A316T missense mutation leading to increased basal desensitisation and turnover, and resulting in reduced cell surface expression of the mutant receptor, which can be rescued by lysosomal inhibition. We also observe varying responses to stimulation with different GLP-1RAs, with Gα_s_-biased agonists such as exendin-F1 associated with a more favourable signalling profile compared to more those of balanced agonists such as exendin-4 and semaglutide (see table S5 for a summary of the main results of the study). These data have important implications for the design of future GLP-1R agonists and for precision medicine approaches, whereby structural residue-residue interactions could be emulated to achieve allosteric effects similar to those present in the A316T variant, or individuals could be prescribed selected GLP-1RAs based on their specific *Glp1r* genetic profile. Our study also serves as a model for future studies to be carried out with other gene variants of the *Glp1r* and other related receptors which are the target of antidiabetic and anti-obesity treatments.

## Materials and Methods

### Peptides, plasmids, and reagents

GLP-1(7–36)NH_2_ was purchased from Bachem; semaglutide was obtained from Imperial College London Healthcare NHS Trust pharmacy; exendin-4 and exendin-F1 were custom synthesized by Wuxi AppTec/Insight Biotechnology and were at least 95% pure. Tetramethylrhodamine (TMR)-labelled exendin-9 and exendin-4 (exendin-9-TMR and exendin-4-TMR) have been described and validated before (*72, 73*). Exendin-9-Cy5 (LUXendin645) has been described before (*74*) and is a kind gift from Prof David Hodson (Oxford Centre for Diabetes, Endocrinology and Metabolism, UK). SNAP-Surface fluorescent probes were purchased from New England Biolabs. Surface cleavable BG-S-S-649 (*75–77*) was a gift of Dr Ivan Corrêa Jr, New England Biolabs. LgBiT-mini-G_s_ (*78*) was a gift from Prof Nevin Lambert (Augusta University, USA). LgBiT-β-arrestin 2 (*53, 79*) was purchased from Promega. pSNAP/FLAG-tagged human *Glp1r* (*hGlp1r*)^A316T^ and *hGlp1r*^A316T^-SmBiT constructs were generated *in house* by site-directed mutagenesis from pSNAP/FLAG-*hGlp1r*^wt^ (Revvity) and *hGlp1r*^WT^-SmBiT (*53*) with the following primers: Forward 5’-CCGGCTGCCCATTCTCTTTACCATTGGGG-3’, and Reverse 5’-ACCCCAATGGTAAAGAGAATGGGCAGCCG-3’ for pSNAP/FLAG-*hGlp1r*^A316T^; Forward 5’-CCGGCTGCCCATTCTCTTTACCATTGGGG-3’, and Reverse 5’-ACCCCAATGGTAAAGAGAATGGGCAGCCG-3’ for *hGlp1r*^A316T^-SmBiT, using the QuikChange Site-Directed Mutagenesis Kit (Agilent) following the manufacturer’s instructions.

### Animal studies

All animal procedures were approved by the British Home Office under the UK animals (Scientific Procedures) Act 1986 (Project License number PP7151519 to Dr Aida Martinez-Sanchez, Imperial College London, UK) and from the local ethical committee (Animal Welfare and Ethics Review Board) at the Central Biological Services unit of Imperial College London. Animals were housed in groups of up to four adult mice in individually ventilated cages under controlled conditions at 21-23°C with 12h:12h light to dark cycles (lights on at 07:00). Ad libitum access to water and feed was provided unless otherwise stated. Lean mice were fed a standard chow diet [RM1 (E); Special Diet Services]. For high-fat high-sucrose (HFHS) diet studies, animals were put on a 58 kcal % fat and sucrose diet (D12331, Research Diets, Inc.) ad libitum for 12-14 weeks prior to experiments.

### Generation of hGlp1r^A316T/A316T^ and hGlp1r^-/-^ mice

Transgenic mice were generated on a C57BL/6 *hGlp1r^+/+^* mouse background from Taconic Artemis GmbH that has been described before (*31*). To generate mice with the *hGlp1r* A316T mutation, one-cell stage *hGlp1r*^+/+^ embryos were electroporated with 100 µL Opti-MEM (Thermo Fisher Scientific) containing ribonucleoprotein (RNP) complexes of *S. pyogenes* Cas9 (1.2 µM Alt-R® *S.p.* Cas9 nuclease 3NLS) with 6 µM single guide RNA (sgRNA) and donor single-stranded repair oligo DNA nucleotides (ssODNs, 300 ng/µL) using the NEPA21 electroporator (NEPA GENE Co. Ltd.) DNA nucleotides, *S.p.* Cas9 nuclease and sgRNA were purchased from Integrated DNA Technologies. The sgRNA used to introduce the Ala316 to Thr mutation was 5’-GCCCATTCTCTTTGCCATTG-3’, designed using RGEN tool (http://www.rgenome.net/cas-designer/), targeting a sequence in exon 9 of *hGlp1r* flanking the rs10305492 single nucleotide polymorphism (SNP). The donor ssODNs in this study are: Forward 5’-CATGAACTACTGGCTCATTATCCGGCTGCCCATTCTCTTTACCATTGGGGTGAACTTCCTCATCTTTGTTCGGGTCATCTGCAT-3’, and Reverse 3’-ATGCAGATGACCCGAACAAAGATGAGGAAGTTCACCCCAATGGTAAAGAGAATGGGCAGCCGGATAATGAGCCAGTAGTTCATG-5’, designed using the Alt-R CRISPR HDR design tool (https://eu.idtdna.com/pages/tools/alt-r-crispr-hdr-design-tool/) with the same target sequence as for the sgRNA (fig. S1A). Following electroporation, two-cell stage embryos were placed into oviducts of pseudo-pregnant *hGlp1r*^+/+^ foster mothers to generate the F0 founder mice, which were subsequently screened for the presence of the A316T substitution as described below (fig. S1b). Mice with frameshift disruptions of *hGlp1r* were also selected from F0 to establish a *hGlp1r*^-/-^ mouse colony.

### Genomic DNA extraction and genotyping

To genotype the transgenic mice, ear samples were collected from weaned animals and genomic DNA extracted in alkaline lysis buffer (25 mM NaOH, 0.2 mM EDTA, pH 8.0) for 1 hour at 95°C and subsequently neutralised by addition of 13 mM Tris-HCl (pH 7.4) (*53*). The purified genomic DNA was used as a template for PCR with appropriate primers prior to Sanger sequencing to identify the A316T mutation. The genotyping primers are described in table S6.

### Oral glucose tolerance tests (OGTTs) and intraperitoneal glucose tolerance tests (IPGTTs)

For OGTTs, chow or HFHS fed *hGlp1r*^A316T/A316T^ mice and littermate controls were fasted for 5 hours, and blood glucose levels serially assessed at 0, 10, 30, and 60 minutes via tail venipuncture using a handheld glucometer (Accu-Chek), after administration of a glucose challenge (2 g/kg of body weight) into the gut by oral gavage. To examine acute and prolonged glycaemic responses to GLP-1R agonists (GLP-1RAs), animals were fasted for 2 hours, starting at 8 a.m. on the morning of the experiments, prior to IPGTTs. GLP-1RAs were then co-administered at the indicated concentrations with 20% (w/v) D-glucose via IP injection at weight-adjusted volumes. Baseline glucose readings were taken via tail venipuncture, and subsequent readings taken at various time-points after glucose ± GLP-1RA administration as for the OGTTs (0-hour “acute” IPGTTs). For GLP-1RAs with shorter half-lives (exendin-4 and exendin-F1), IPGTTs were repeated 6 hours after the start of the initial IPGTT, with continued fasting. For the extended half-life GLP-1RA semaglutide, IPGTTs were performed 24 and 72 hours after GLP-1RA administration, with access to food restored at the end of each IPGTT. The *in vivo* IPGTT studies were performed with a crossover design, with each mouse receiving vehicle, and each GLP-1RA treatment with washout periods of at least one week between the different GLP-1RA treatments.

### Measurement of in vivo plasma insulin levels

To measure insulin release during IPGTTs, 10 µL blood samples were collected via tail venipuncture into potassium EDTA cuvettes (Microvette CB 300, 16.444.100, Starstedt) 10 minutes after glucose challenge, and stored on ice before centrifugation at 8,000 x g for 10 minutes at 4°C to collect plasma supernatants. Plasma was collected in fresh Eppendorf tubes and kept at −80°C until further analysis using the Mouse Plasma Insulin Homogeneous Time Resolved Fluorescence (HTRF) kit (#62IN3PEF; Revvity) following the manufacturer’s instructions.

### Immunostaining of pancreatic sections

Whole pancreata from *hGlp1r*^A316T/A316T^ mice and *hGlp1r*^+/+^ control littermates were dissected, placed in ice cold phosphate buffer saline (PBS), cleared from surrounding fat and non-pancreatic tissues, weighed, and fixed for 24 hours in 4% paraformaldehyde (PFA) at 4°C and then cryoprotected in 30% sucrose, followed by embedding in optical cutting temperature (OCT) and stored at −80°C. Frozen 10 µm sections were prepared using a cryostat and immunostained with guinea pig anti-insulin polyclonal antibody (1:100; Dako IR002, Agilent Technologies; Alexa Fluor 488 secondary antibody, Thermo Fisher Scientific) for β-cell, and mouse anti-glucagon antibody (1:500; G2654, Sigma-Aldrich; Alexa Fluor 568 secondary antibody, Thermo Fisher Scientific) for α-cell detection. Samples were mounted with Prolong Diamond Antifade Mountant with DAPI (P36962, Thermo Fisher Scientific) for nuclei visualisation. Stained slides were imaged at 4x to quantify total pancreatic area, and at 40x to image pancreatic islet insulin and glucagon positive areas using a wide-field Zeiss Axio Observer inverted microscope from Imperial College Facility for Imaging by Light Microscopy (FILM). Total pancreatic, insulin and glucagon positive areas were measured per slide using the Fiji software. β- and α-cell mass was calculated by normalising to total pancreatic section area and pancreas weight as described before (*80, 81*). Islet size and cytoarchitecture analyses were performed on the same images used for β- and α-cell mass assessments. All non-islet regions were erased using the ImageJ outline and clear outside functions. Cells were assigned to be in the islet periphery/mantle if they were within the 2 outermost cell layers of the islet, or in the islet core if they fell within any layers deeper than these.

### In vivo hGLP-1R labelling in the pancreas

The *hGlp1r*^A316T/A316T^ mice and *hGlp1r*^+/+^ control littermates were injected subcutaneously with 100 pmol/g exendin-9-Cy5 two hours before terminal anaesthesia (1.5% sodium pentobarbital) and transcardial perfusion fixation with PBS at pH 7.2, supplemented with 4% fresh PFA (*2*). Pancreata were dissected and frozen sections prepared and mounted as above. Images of sections were captured using a Nikon Eclipse Ti spinning disk confocal microscope and 60x oil immersion objective with an ORCA-Flash 4.0 camera (Hamamatsu) and Metamorph software (Molecular Devices), with λ_ex_ □= 660 nm and fluorescent intensity in islet areas normalised by number of islet cell nuclei calculated in Fiji.

### Isolation and maintenance of islets

The appropriate animals were humanely euthanised prior to pancreas inflation via injection of RPMI 1640 medium (S1745602, Nordmark Biochemicals) supplemented with 1 mg/mL collagenase from Clostridium histolyticum (S1745602, Nordmark Biochemicals) into the common bile duct. The pancreas was then dissected and incubated in a water bath at 37°C for 10 minutes. Digestion was quenched with cold mouse islet culture medium: RPMI 1640 supplemented with 10% (v/v) foetal bovine serum (FBS) (F7524, Sigma-Aldrich) and 1% (v/v) penicillin/streptomycin (P/S) (15070-063, Invitrogen). Islets were subsequently washed, purified on a Histopaque gradient (Histopaque-1119 and −1083, Sigma-Aldrich), and allowed to recover overnight at 37°C in 5% CO_2_ in mouse islet culture medium prior to any experiments. Human donor islets from deceased heart-beating donors of both sexes were provided by the European Diabetes Study Centre (CEED), Strasbourg, France. Human pancreatic tissue was collected with consent from next of kin where required, following the French bioethics legislation and INSERM guidelines (French bylaw, published on December 29, 1998), and national and ethical approvals in place. Human islets were cultured in RPMI 1640 supplemented with L-glutamine, 10% (v/v) FBS, 1% (v/v) P/S, 0.25 β g/mL amphotericin B, and 5.5 mM glucose at 37°C in 5% CO_2_. Human donor characteristics are listed in table S7.

### Cell line generation and culture

INS-1 832/3 cells stably expressing pSNAP/FLAG-*hGlp1r*^WT^ or ^A316T^ were derived *in house* from the parental INS-1 832/3 *Glp1r*^-/-^ cell line, where endogenous rat GLP-1R has been eliminated by CRISPR/Cas9 (*46*), a gift from Dr Jacqui Naylor, AstraZeneca, by transfecting the corresponding plasmid followed by selection in 1 mg/mL G418 and fluorescence-activated cell sorting (FACS) of the SNAP-positive cell population, and maintained in RPMI-1640 with 11 mM glucose, 10 mM HEPES, 2 mM glutamine, 1 mM sodium pyruvate, 50 βM β-mercaptoethanol, 10% FBS, 1% P/S and 1 mg/mL G418. Human EndoC-βH3 cells (Human Cell Design) were cultured on 2 □*µ*g/mL fibronectin and 1% extracellular matrix (E1270, Sigma-Aldrich) coated plates in Advance DMEM/F-12 (Thermo Fisher Scientific), 6.7 ng/ml sodium selenite, 10 mM nicotinamide, 5.5 □*µ*g/mL human transferrin, 2mM L-glutamine, 2% BSA Fraction V (Roche), 1% P/S solution and 50 □µM β-mercaptoethanol. All cells were incubated in a humidified air incubator containing 5% CO_2_ at 37°C.

#### Generation and characterisation of EndoC-βH3 hGlp1r^-/-^ cells

##### a) CRISPR/Cas9 dgRNA design

To generate EndoC-βH3 *hGlp1r*^-/-^ cells, a lentiviral CRISPR/Cas9 approach was used as previously described (*82*). Cas-Designer (http://www.rgenome.net/cas-designer/) was used to design two 17 nucleotide-long (truncated) guide RNAs (gRNAs), 5’-GCTGCTCGGGATGGTGGGCA-3’ and 5’-GTTGCAGAACAAGTCTGTGG-3’ (reverse complement), flanking the target DNA sequences. Designed gRNAs were cloned into a pSpCas9(BB)-2A-Blast backbone (plasmid #118055, Addgene), which allows for blasticidin selection of cells after transfection, following a previously described dual gRNA cloning strategy (*82*) as follows: oligonucleotides corresponding to deletion-yielding pairs of gRNAs were assembled in a reaction mix containing 1x Q5 reaction buffer, 200 µM dNTPs, 0.5 µM primer, 0.02 U/µL Q5 high-fidelity DNA polymerase and 0.25 ng/µL of pScaffold-H1 vector (plasmid #118152, Addgene), with Ta = 58°C, 15 second extension, for 30 amplification cycles. PCR products were digested and ligated to pSpCas9(BB)-2A-Blast in a one-step digestion ligation reaction with 1x Tango buffer, 1 µL FastDigest BbsI (Thermo Fisher Scientific), 1µL DTT 0.1M, 1µL ATP 10 mM, 0.5 µL T7 ligase, 100 ng vector and 1 µL of PCR product (diluted 1:20). Digestion-ligation reactions underwent 6 cycles of 5 minutes at 37°C and 5 minutes at 23°C, and a final 5minute incubation at 37°C, after which 1 µL of ligation product was transformed into NEB Stable competent *E. coli* (New England Biolabs). Successful double gRNA (dgRNA) insertion was confirmed by Sanger sequencing.

##### b) Lentiviral production and transduction of EndoC-βH3 cells

To produce CRISPR/Cas9 lentiviral particles, HEK293T cells were co-transfected with dgRNA vector from above as well as packaging (psPAX2) and envelope (pMD2.G) vectors using a calcium phosphate protocol (*83*). Viral supernatants were harvested 48 and 72 hours after transfection, cleared using a 0.45-µm Millex-HV filter, and concentrated by adding PEG 8000 (Sigma) and NaCl until final concentrations of 10% and 0.15 M, respectively. The mixture was then incubated for 16 to 20 hours on a shaker at 4°C, followed by centrifugation for 30 minutes at 10,000 g at 4°C. Virus pellets were resuspended in 100 µL PBS and stored at −80°C. 48 hours before transduction, EndoC-βH3 cells were seeded at 1 million cells per well in 6-well plates, and 50 β L of virus used to infect each well. After 72 hours, cells were selected for vector integration using blasticidin (20 µg/mL) for 10 days, with selection medium changed every ∼3 days (fig. S1C). After selection, cells were labelled with fluorescent exendin-9-TMR for 10 minutes and reverse FACS sorted to recover the population enriched in non-labelled *hGlp1r*^-/-^ cells.

##### c) EndoC-βH3 hGlp1r^-/-^ validation by HTRF cAMP assay

EndoC-βH3 WT and *hGlp1r*^-/-^ cells were pre-treated for 3 weeks with 1 µM of 4-hydroxytamoxifen to induce beta cell differentiation (*45*). On the day of the experiment, cells were incubated in 0.5 mM glucose comtaining EndoC-βH3 medium ± 100 nM exendin-4 and the phosphodiesterase inhibitor isobutyl methylxanthine (IBMX). Cells were then lysed in in Krebs-Ringer bicarbonate-HEPES (KRBH) (140 mM NaCl, 3.6 mM KCl, 1.5 mM CaCl_2_, 0.5 mM MgSO_4_, 0.5 mM NaH_2_PO_4_, 2 mM NaHCO_3_, 10 mM HEPES, saturated with 95% O_2_/5% CO_2_; pH 7.4) buffer + 1% Triton X-100 and lysates used to determine cAMP production using a cAMP Dynamic 2 homogeneous time-resolved fluorescence (HTRF)-based assay kit (Revvity) in a PHERAstar reader (BMG Labtech), following the manufacturer’s instructions. Results show decreased cAMP responses in *EndoC-βH3 hGlp1r^-/-^ versus* WT cells (fig. S1D), suggesting enrichment in *hGlp1r^-/-^*cells within the CRISPR/Cas9-transduced cell population.

### Insulin secretion assays

Purified mouse islets used for insulin secretion assays were treated in 24-well non-adherent plates. Eight islets were used per replicate with three technical replicates per condition. For acute studies, islets were preincubated for 1 hour in KRBH buffer containing 0.1% (w/v) bovine serum albumin (BSA) (10775835001, Roche) and 3 mM glucose before incubation with 11 mM glucose ± 100 nM GLP-1RAs in KRBH buffer in a shaking 37°C water bath (80 rpm) for 1 hour. For overnight studies, preincubation was carried out in RPMI 1640 medium containing FBS, P/S, and 3 mM glucose, followed by treatment with medium supplemented with 11 mM glucose ± GLP-1RAs for 16 hours. Supernatants containing secreted insulin were collected, centrifuged at 1000 rpm for 1 minute, and transferred to fresh tubes. For total insulin content, islets were lysed using acidic ethanol (75% ethanol, 1.5 mM HCl). Lysates were sonicated 3x 10 seconds each in a water bath and centrifuged at 10,000 g for 10 minutes, and supernatants collected. For human islets, the same experiment was performed with islets previously transduced with pAV-SNAP/FLAG-*hGlp1r*^WT^ or ^A316T^ adenoviruses (generated by VectorBuilder) at a multiplicity of infection (MOI) of 1, 24 hours prior to the start of experiments. Insulin secretion assays with the human EndoC-βH3 cell line were performed following 3 weeks of treatment with 4-hydroxytamoxifen as above, followed by transduction with pAV-SNAP/FLAG-*hGlp1r*^WT^ or ^A316T^ adenoviruses, with insulin secretion assessed as previously described (*45*), with the following modifications: after preincubation with 0.5 mM glucose KRBH buffer, cells were stimulated with 15 mM glucose ± exendin-4 for 60 minutes. Total and secreted insulin samples from islets and cells were stored at −20°C until quantification using an Insulin Ultra-Sensitive HTRF kit (62IN2PEG, Revvity) in a PHERAstar plate reader (BMG Labtech), according to the manufacturer’s instructions.

### Real-time cAMP imaging

To measure cAMP, *hGlp1r*^A316T/A316T^ *versus* ^+/+^ control mouse islets, or INS-1 832/3 SNAP/FLAG-*hGlp1r*^WT^ *versus* ^A316T^ cells cultured on glass-bottom MatTek dishes, were transduced with cADDis (Green Gs cADDis cAMP Kit, Montana Molecular), a baculovirus genetically encoding a fluorescent cAMP *in situ* biosensor, following the manufacturer’s instructions. Transduced islets were subsequently encased into Matrigel on glass-bottom MatTek dishes 24 hours after transduction, and both islets and cells were imaged at λ_ex_ □= 488 nm in a Nikon Eclipse Ti spinning disk confocal microscope and 20x objective with an ORCA-Flash 4.0 camera (Hamamatsu) for time-lapse recordings with image acquisitions every 6 seconds in KRBH buffer containing 0.1% (w/v) BSA and 6 mM glucose for 2 minutes to record the baseline, with 100 nM GLP-1RA subsequently added by manual pipetting, and islets imaged for a further 5 minutes before addition of 100 β M IBMX + 10 βM forskolin for the final 2 minutes of the acquisition. Raw green fluorescence intensity traces were extracted from whole islet regions of interest (ROIs) using Fiji, and mean intensities calculated for each ROI and time point. Responses were plotted relative to average fluorescence intensity per islet during the baseline period before GLP-1RA addition.

### Islet immunofluorescence

For immunostaining, *hGlp1r*^A316T/A316T^ or ^+/+^ control mouse islets were fixed in 4% PFA and stored in PBS. After 15 minutes of permeabilisation with 0.5% (v/v) Triton X-100 in PBS, the islets were washed once with PBS and incubated in blocking buffer (PBS + 1% BSA) for 1 hour. Primary antibodies against insulin (1:50; Dako IR002, Agilent Technologies) and glucagon (1:500; G2654, Sigma-Aldrich) were added overnight in blocking buffer, followed by 3x washes in PBS and 1 hour incubation at room temperature with secondary anti-guinea pig Alexa Fluor 488 and anti-mouse Alexa Fluor 647 (both at 1:500; Thermo Fisher Scientific). Islets were then loaded onto glass-bottom MatTek dishes in PBS and three images acquired at different z positions by confocal microscopy in a Zeiss LSM-780 inverted confocal laser-scanning microscope and a 63×/1.4 numerical aperture oil immersion objective from Imperial College FILM Facility. Images were analysed using Fiji.

### Measurement of mini-G_s_ and β-arrestin 2 recruitment by NanoBiT complementation

For mini-G_s_ recruitment, INS-1 832/3 *Glp1r*^-/-^ cells were seeded in 6-cm dishes and transiently co-transfected with 1.7 µg each of *hGlp1r*^WT^ or ^A316T^-SmBiT and LgBiT-mini-G_s_ constructs for 24 hours. Cells were then detached, resuspended in Hank’s buffered salt solution (HBSS) containing NanoGlo Live Cell Reagent (1:20; Promega) with furimazine, and seeded onto white 96-well half-area plates. Baseline luminescence was immediately recorded over 8 minutes at 37°C in a Flexstation 3 plate reader (Molecular Devices), and for a further 30 minutes after GLP-1RA addition at serial doses up to 100 nM. Readings were taken every 30 seconds and normalised to average well baseline, and average vehicle-induced signal subtracted to establish the agonist-induced effect. To measure β-arrestin 2 recruitment, cells were processed as above with slight modifications: INS-1 832/3 *Glp1r*^-/-^ cells were co-transfected with 1.77 *µ*g each of *hGlp1r*^WT^ or ^A316T^-SmBiT and LgBiT-β-arrestin 2 plasmids for 24 hours and GLP-1RAs added at serial doses up to 1 *µ*M. Areas under the curve (AUCs) were calculated for each agonist concentration and fitted to log(agonist) *versus* response four-parameter curves using GraphPad Prism 10.1.2, with errors propagated for mini-G_s_ over β-arrestin 2 recruitment bias calculations.

### High content microscopy GLP-1R trafficking assays

#### a) Internalisation assay

INS-1 832/3 SNAP/FLAG-*hGlp1r*^WT^ or ^A316T^ cells were seeded onto poly-D-lysine-coated black 96-well plates for 24 hours before the assay. Cells were labelled for 30 minutes with 1 μM SNAP surface cleavable BG-SS-649 probe at 37°C and washed once with HBSS before treatment with GLP-1RA or vehicle for 60, 30, 15, or 5 minutes at 37°C in complete medium. Cells were then washed with cold HBSS, and subsequent steps carried out at 4°C. The cell impermeable reducing agent Mesna [100 mM in alkaline TNE buffer (20 mM TrisHCl, 1 mM EDTA, 150 mM NaCl, pH 8.6)] or alkaline TNE buffer without Mesna was applied for 5 minutes to cleave BG-SS-649 probe left at the cell surface and then washed with HBSS. Plates were imaged immediately without fixation using a modular microscope platform from Cairn Research incorporating a Nikon Ti2E, LED light source (CoolLED), fitted with a 20x phase contrast objective, assisted by custom-written high content analysis software implemented in Micro-Manager. A minimum of 9 images per well were acquired for both epifluorescence and transmitted phase contrast. Internalised GLP-1R, indicated by cell-associated fluorescence, was analysed after flat-field correction of fluorescent images using the BaSiC plugin (*84*), segmentation of cell-containing regions from phase contrast images using Phantast (*85*), and fluorescence intensity quantification. The percentage of internalised receptor was calculated as follows:

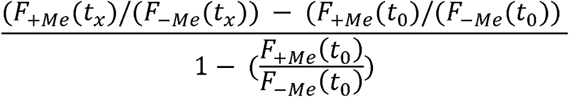

Where *F_+Me_* and *F_-Me_* are mean fluorescence intensities ± MesNa at times *t_x_* (5, 15, 30, or 60 minutes) or *t_0_* (0 minutes). Percentages of internalisation were plotted to calculate the AUC using GraphPad Prism 10.1.2.

#### b) Recycling assay

INS-1 832/3 SNAP/FLAG-*hGlp1r*^WT^ or ^A316T^ cells were seeded as above for 24 hours, washed with HBSS, pre-treated with vehicle or GLP-1RA for 1 hour at 37°C in complete medium, washed again and treated with 100 nM exendin-4-TMR in reverse time order for 3 hours and 1 hour at 37°C in complete medium. Plates were then promptly imaged after washing as above. In this assay, total exendin-4-TMR uptake is indicative of residual surface receptor at the end of the GLP-1RA treatment and cumulative reappearance of the surface receptor during the recycling period.

#### c) Degradation assay

INS-1 832/3 SNAP/FLAG-*hGlp1r*^WT^ or ^A316T^ cells were seeded as above. Cells were then washed with HBSS before addition of 50 μg/mL cycloheximide in serum-free medium to halt protein translation. After one hour, a time course was performed with the GLP-1RA added in reverse time order for 8, 4, 2, 1, 0.5, and 0 hours. For the final 30 minutes of the experiment, the cell medium was replaced with 1 μM of total SNAP-labelling fluorescent probe BG-OG in complete medium, to label total residual GLP-1R. Cells were then washed 3x with HBSS prior to plate imaging as above using a green fluorescence filter set. Total GLP-1R was quantified as the average cellular fluorescence labelling per well (as defined above), with further subtraction of non-specific fluorescence (i.e. cells without BG-OG labelling). Results were expressed as a percentage relative to vehicle treated BG-OG labelled cells and AUCs recorded as above.

### Lysosomal inhibition assay

INS-1 832/3 SNAP/FLAG-*hGlp1r*^WT^ or ^A316T^ cells on coverslips, mouse *hGlp1r*^-/-^ islets transduced with pAV-SNAP/FLAG-*hGlp1r*^WT^ or ^A316T^ adenoviruses, or *hGlp1r*^A316T/A316T^ and ^+/+^ control islets were incubated for 2 hours ± 100 nmol/L bafilomycin-A1 (Sigma-Aldrich), an established vacuolar H^+^-ATPase and lysosomal trafficking inhibitor (*86*), in complete medium. Cells and transduced *hGlp1r* ^-/-^islets were then labelled with 1 µM SNAP-Surface 549, and *hGlp1r*^A316T/A316T^ *versus* ^+/+^ islets treated with exendin-9-TMR in complete medium at 37 °C for 10 minutes. Cells were washed and fixed in 4% PFA, mounted in Prolong Diamond antifade reagent with DAPI, while islets were loaded onto glass-bottom MatTek dishes in PBS and images acquired by confocal microscopy with a Zeiss LSM-780 inverted confocal laser-scanning microscope and a 63x/1.4 numerical aperture oil immersion objective from the Imperial College FILM Facility. Images were analysed using Fiji.

### Surface GLP-1R expression analysis

INS-1 832/3 SNAP/FLAG-*hGlp1r*^WT^ or ^A316T^ cells, or EndoC-βH3 cells transduced with pAV-SNAP/FLAG-*hGlp1r*^WT^ or ^A316T^ adenoviruses were labelled with 1 µM SNAP-Surface 549 at 37°C in 5% CO_2_, washed twice in PBS and fixed with 4% PFA at 4°C for 20 minutes, mounted onto slides in Diamond Prolong anti-fade Mountant with DAPI (Thermo Fisher Scientific), and images acquired by confocal microscopy with the Zeiss LSM-780 inverted confocal laser-scanning microscope from above using a 20x objective. Human and *hGlp1r*^-/-^ mouse islets transduced with pAV-SNAP/FLAG-*hGlp1r*^WT^ or ^A316T^ adenoviruses were labelled with 1 µM SNAP-Surface 649 for 10 minutes, washed with PBS and imaged as above with a 63x/1.4 numerical aperture oil immersion objective. *hGlp1r*^A316T/A316T^ *versus* ^+/+^ transgenic mouse islets were labelled with a saturating concentration (1 µM) of exendin-9-TMR in complete medium at 37°C for 10 minutes and imaged as above. Images were analysed using Fiji.

### RNA extraction and qPCR

Total RNA was extracted from islets (50–100) isolated from *hGlp1r*^A316T/A316T^ *versus* ^+/+^ mice, using TRIzol reagent (Invitrogen) and briefly vortexed for homogenisation. Chloroform was added to achieve phase separation, and the upper aqueous phase collected. RNA was recovered overnight by precipitation with isopropanol (Thermo Fisher Scientific) and 100-500 ng reverse transcribed using the MultiScribe Reverse Transcriptase (Thermo Fisher Scientific) according to the manufacturer’s instructions. Quantitative PCR (qPCR) using SYBR Green Technology was carried out on an Applied Biosystems 7500 real-time PCR system. Data were analysed using the 2−ΔΔCt method (*87*). A list of qPCR primer sequences is provided in table S8.

### Protein extraction and Western blotting

INS-1 832/3 SNAP/FLAG-*hGlp1r* ^WT^ or ^A316T^ cells were treated with vehicle or exendin-4 for 5 minutes, washed once in cold PBS and lysed 1x TNE lysis buffer (20 mM Tris, 150 mM NaCl, 1 mM EDTA, 1% NP40, protease and phosphatase inhibitors) for 10 minutes at 4°C. Samples were then sonicated in a water bath sonicator 3x 10 seconds and centrifuged at 13,000 rpm for 5 minutes. Supernatants were collected and resolved by SDS-polyacrylamide gel electrophoresis (10% acrylamide gels) in 2x urea loading buffer (200mM Tris HCl, 5% w/v SDS, 8 M urea,100 mM DTT, 0.02% w/v bromophenol blue, pH 6.8). A Bio-Rad wet transfer system was used for immunoblotting onto PVDF membranes (Immobilon-P, 0.45-βm pore size, IPVH00010, Merck) before incubation with appropriate primary and secondary antibodies in 5% skimmed milk. SNAP-tagged GLP-1R was detected with an anti-SNAP tag rabbit polyclonal antibody (1:500; P9310S, New England Biolabs) followed by goat anti-rabbit IgG HRP (1:2,000; Abcam). Post-stripping, Tubulin was labelled with anti-β-tubulin mouse monoclonal antibody (1:5,000; T5168, Sigma) followed by sheep anti-mouse secondary antibody HRP (1:5,000; Abcam). Blots were developed using Clarity™ Western ECL Substrate System (BioRad) in a Xograph Compact X5 processor and specific band densities quantified in Fiji.

### Cryo-EM determination of GLP-1:GLP-1R (A316T):Gα_s_ complex structure

#### a) Construct design

A human GLP-1R construct with residue Ala316 substituted for Thr (c.946G>A) was generated from the WT sequence by Q5 enzyme site-directed mutagenesis (New England Biolabs) and cloned into the pFastBac vector. The native signal peptide was replaced with HA to enhance expression, and an N-terminal FLAG epitope and a C-terminal 6xHis epitope added as affinity tags with 3C protease cleavage sites. These modifications do not significantly alter the function of WT GLP-1R (*88*).

#### b) Purification of GLP-1:GLP-1R (A316T):DN-Gα_s_:Nb35

A-FLAG-GLP-1R (A316T)-His, DN-Gα_S_, and G_β1β2_-His P2 baculoviruses were co-infected in a 500 mL culture of *T. ni* Hi5 insect cells in suspension (density 3-4×10^6^ cells/mL), at a ratio of 2:1:1. After 48 hours, the cell preparations were harvested by centrifugation at 8,000 g for 10 minutes and the supernatant discarded. The pelleted cell mass was stored at −80°C until use. For purification, cell preparations were rapidly thawed and resuspended in 50 mL of buffer. Receptor-G protein complex was formed in the presence of 10 βM GLP-1 for 30 minutes at room temperature with gentle stirring. After this period, nanobody 35 (Nb35, 1 mg) and apyrase (5 µL/L of starting culture, equivalent to 2.5 U) were added to the resuspension and incubated at room temperature for a further 30 minutes. Purified Nb35 was a kind gift from Minakshi Baruah.

#### c) Vitrification and data collection of purified samples

2 µL of sample was applied to glow-discharged R1.1/1.2 gold foil grids, manufactured by Quantifoil at the Melbourne Centre for Nanofabrication (MCN) immediately followed by plunge-freezing in liquid ethane using a Vitrobot Mark IV (Thermo Scientific) set at 4°C, 100% humidity. At least 3 grids were prepared for each sample, and blot force and blot time varied between 16-18 and 6-8 seconds, respectively. Vitrification conditions for grids used in data collection were as follows: blot force 16 and blot time 7 seconds. cryo-EM data were collected on a Glacios cryo-TEM (Thermo Scientific) operating at 200 kV accelerating voltage in cryo-TEM nanoprobe mode with a 100 βm objective aperture inserted. Data were collected using aberration-free image shift in fast acquisition mode in the EPU software (Thermo Scientific).

#### d) Data processing for GLP-1:GLP-1R (A316T):DN-Gα_s_:Nb35

3,170 micrographs were assigned to 69 optics groups based on their beam-image shift coordinates, motion corrected in RELION 3.1.2 and CTF parameters estimated using CTFFIND 4.1 within RELION (*89*). For quality control, micrographs with an estimated maximum resolution >20 Å were excluded, leaving 2,940 micrographs in the dataset. 1,667,111 particles were picked using a pre-trained general model in crYOLO 1.7.6 (*90*) and extracted with 4.5-fold binning. These particles were imported into cryoSPARC 3.2 (*91*) and subjected to rounds of 2D classification to homogenise the particle stack, with 583,410 particles remaining. These particles were re-extracted at the full box size (288 px2) and subjected to Bayesian polishing in RELION 3.1.2. The polished particles were re-imported into cryoSPARC, filtered for polishing artefacts by 2D classification, and a final set of 330,000 was subjected to non-uniform refinement to obtain a 3.3 Å global resolution consensus density map. Atomic model refinement used an existing model of WT GLP-1R bound to GLP-1 and DN-Gα_s_ proteins (PDB: 6X18) where Ala316 of GLP-1R was mutated to Thr. The mutated model was fitted into the experimental density map and subjected to MDFF refinement using Isolde (*92*) within ChimeraX 1.5 (*93*). Iterative rounds of real-space refinement in PHENIX 1.20 (*94*), followed by MDFF in Isolde, were used to obtain the final model for analysis. Data collection, processing and refinement statistics for the cryo-EM of GLP-1:GLP-1R (A316T):DN-Gα_s_:Nb35 complex are summarised in table S9.

### Computational methods and MD simulations

#### a) MD simulations of the A316T and WT GLP-1R structures

The cryo-EM GLP-1:GLP-1R (A316T):Gα_s_ complex structure and the corresponding GLP-1:GLP-1R(WT):Gα_s_ (PDB 6X18 (*56*)) were subjected to the same preparation for MD simulations. The missing stalk region (residues 130-135) and ICL3 (residues 339-343) of 6X18 were added by superimposition from the GLP-1:GLP-1R:Gα_s_ structure simulated in (*9*) and Gα_s_ was removed. The resulting systems (i.e., WT and A316T) were superimposed with the PDB 6X18 OPM reference (*95*) in order to opportunely orient the receptor, and parameterized with the CHARMM36 force field (*96*), employing in-house Python HTMD (*97*) and Tool Command Language (TCL) scripts. Hydrogen atoms were added at a simulated pH of 7.0 using the pdb2pqr (*98*) and propka (*99*) software, and the protonation of titratable side chains was checked by visual inspection to correctly orient the receptor before inserting (*100*) in a rectangular 100 Å x 101 Å 1-palmitoyl-2-oleyl-sn-glycerol-3-phosphocholine (POPC) bilayer (previously built by using the VMD Membrane Builder plugin 1.1 at http://www.ks.uiuc.edu/Research/vmd/plugins/membrane/), removing the lipid molecules overlapping the receptor TMD bundle. TIP3P water molecules (*101*) were added to the simulation box (100 Å x 101 Å x 136 Å) employing the VMD Solvate plugin 1.5 at http://www.ks.uiuc.edu/Research/vmd/plugins/solvate/. Overall charge neutrality was finally reached by adding Na^+^/Cl^-^ counter ions (final ionic concentration of 0.150 M), using the VMD Autoionize plugin 1.3 at http://www.ks.uiuc.edu/Research/vmd/plugins/autoionize/. For WT and A316T GLP-1Rs, equilibration and production replicas were computed using ACEMD (*102*). For the equilibration, isothermal-isobaric conditions (Berendsen barostat (*103*) with a target pressure of 1 atm; Langevin thermostat (*104*) with a target temperature of 300 K and damping of 1 picosecond^-1^) were employed to equilibrate the systems through a multi-stage procedure (integration time step of 2 femtoseconds). First, clashes between lipid atoms were reduced through 1,500 conjugate-gradient minimisation steps. A positional constraint of 1 kcal/mol Å^-2^ lipid phosphorus atoms, protein Ca carbon and all the other protein-heavy atoms were applied and gradually released over 4, 100 and 80 nanoseconds respectively, before further 50 nanoseconds without any restraints for a total of 150 nanoseconds. Productive trajectories (2.5 Å microseconds for each replica, 7.5 microseconds for each system) were computed with an integration time step of 4 femtoseconds in the canonical ensemble (NVT) at 300 K, using a thermostat damping of 0.1 ps^-1^ and the M-SHAKE algorithm (*105*) to constrain the bond lengths involving hydrogen atoms. The cut-off distance for electrostatic interactions was set at 9 Å, with a switching function applied beyond 7.5 Å. Long-range Coulomb interactions were handled using the particle mesh Ewald summation method (PME) (*106*) by setting the mesh spacing to 1.0 Å.

#### b) GLP-1R WT and A316T TMD simulations

Using the GLP-1:GLP-1R:Gα_s_ cryo-EM complex [PDB 6X18 (*56*)] as the starting structure, we removed Gα_s_, GLP-1, ECD, and extracellular loop 1 (ECL1) to minimise the possible interference with the GLP-1R TMD dynamics (fig. S9A). Position 316 of the resulting TMD WT was mutated from Ala to Thr to model the A316T TMD and after membrane insertion and solvation, the two systems (i.e. TMD WT and TMD A316T) underwent 12 milliseconds of MD each; ICL3 residues 339I - 343 were modelled using Modeller 9.18 (*107*) through Chimera (*93*) (v1.14) user interface, considering the best DOPE score. A316T was modelled using Chimera. The resulting GLP-1R TMD systems (i.e., WT and A316T) were prepared for MD simulations as reported above, after superimposition with the PDB 6X18 OPM reference (*95*), with the CHARMM36 force field (*96*) by inserting (*100*) them in a rectangular 96 Å x 96 Å POPC bilayer. For both WT and A316T, three independent equilibrations and production replicas were computed using ACEMD (*102*) and the settings reported above Productive trajectories (4 microseconds for each replica, 12 microseconds for each system) were computed with an integration time step of 4 femtoseconds in the canonical ensemble (NVT) at 300 K.

#### c) MD simulations of exendin-F1 (Ex-F1) in complex with A316T and WT GLP-1R

The Ex-F1:GLP-1R (A316T):Gα_s_ was modelled from the Ex-F1:GLP-1R(WT):Gα_s_ complex structure (PDB 9C0K) by mutating *in silico* A316^5.46^ to T316^5.46^. Ex-F1:GLP-1R(WT):Gα_s_ and Ex-F1:GLP-1R (A316T):Gα_s_ were prepared for MD simulations by superimposing the structures on GLP-1R coordinates retrieved from the OPM database in order to opportunely orient the receptor before being inserted in a rectangular 125 Å x 116 Å POPC bilayer, removing the lipid molecules overlapping the receptor TMs bundle. TIP3P water molecules (*101*) were added to the simulation box (125 Å x 116 Å x 190 Å) and overall charge neutrality was finally reached by adding Na^+^/Cl^-^ counter ions (final ionic concentration of 0.150 M). Equilibration and MD productive simulations computed using ACEMD in isothermal-isobaric conditions (target pressure 1 atm; target temperature 310 K) were employed to equilibrate the systems through 125 nanoseconds of simulation (integration time step of 2 femtoseconds). First, clashes between lipid atoms were reduced through 2,500 conjugate-gradient minimization steps, and then a positional constraint of 1 kcal mol^-1^ Å^-2^ on protein and lipid phosphorus atoms was applied. Lipids restraints were gradually released over a time window of 6 nanoseconds, protein atoms other than alpha carbon atoms were gradually released from the restraints over 100 nanoseconds, while alpha carbon atoms were gradually released over 120 nanoseconds. No restraint was applied in the last 5 nanoseconds of equilibration. Productive trajectories (four 500 nanoseconds-long replicas for each GLP-1R complex) were computed with an integration time step of 4 femtoseconds in the canonical ensemble (NVT) at 310 K as reported above.

#### d) MD analysis

Atom distances were computed at the C µLevel using MDTraj (*108*). H-bonds were quantified using the GetContacts analysis tool (https://getcontacts.github.io/) as the percentage of frames over all the frames obtained by merging the different replicas in which they were present. Water molecule occupancy was computed employing the VMD (*109*) Volmap plugin (https://www.ks.uiuc.edu/Research/vmd/plugins/volmapgui/) with 1 Å resolution.

#### e) Numbering system

Throughout the manuscript, the Wootten residues numbering system for class B GPCRs (*110*) was adopted.

### Statistical analyses

All data analyses and graph generation were performed with GraphPad Prism 10.1.2. The *n* numbers refer to the number of independent biological replicates, representing the number of mice per genotype for *in vivo* experiments, or the number of biologically independent experiments performed from islets isolated from separate mice or from a separate batch of cells, for *ex vivo* or *in vitro* experiments, respectively. Technical replicates within the same assay were averaged to determine the mean value for each biological replicate, prior to statistical tests as indicated in the corresponding figure legends. Statistical significance was assessed with the indicated test, with matched analyses performed for matched design experiments. Statistical significance was inferred when p<0.05. Unless indicated, values represented are mean ± SEM.

## Data availability

Cryo-EM maps and atomic models for GLP-1:GLP-1R (A316T):Gα_s_ are deposited in the PDB and EMDB databases with the following codes: PDB ID 9E2A, EMD-47447. Proteomics data is deposited in the ProteomeXchange (PRIDE) repository with accession number PXD056782 (*55*).

## Code availability

RICS data was analysed with SIM-FCS4 freeware, developed by Prof Enrico Gratton, University of California, Irvine. Software used for the MD simulations is available as follows: Chimera 1.14 (https://www.cgl.ucsf.edu/chimera/download.html); HTMD (https://github.com/Acellera/htmd?tab=readme-ov-file); VMD 1.9.3 (https://www.ks.uiuc.edu/Development/Download/download.cgi?PackageName=VMD); ACEMD3 (https://software.acellera.com/acemd/index.html); MDTraj 1.9.4 (https://mdtraj.org/1.9.4/index.html); GetContacts (https://getcontacts.github.io/).

## Acknowledgements

The authors thank Steve Rothery from the Imperial College Facility for Imaging by Light Microscopy (FILM) for technical support on microscopy experiments and microscopy data analysis; Prof D. Hodson, University of Oxford, for providing LUXendin645; and Elodie Ndjetehe and Zoe Webster from the Transgenics and Embryonic Stem Cell Facility, MRC Laboratory of Medical Sciences, for their help with CRISPR/Cas9 embryo electroporation and generation of knock-in mice. We additionally thank Drs Bryn Owen and Aida Martinez-Sanchez for providing animal project licenses.

## Funding

This work was supported by Diabetes UK PhD Studentship grant 19/0006094 to A.T. and a Society for Endocrinology Research Grant to A.T. and L.E.E. The A.T. group is additionally funded by an MRC Project Grant (MR/X021467/1, with B.J. as co-I), and a Wellcome Trust Discovery Award (301619/Z/23/Z, with B.J. and J.B.S. as co-Is). J.B.S. also acknowledges funding from BBSRC (BB/V019791/1) and MRC (MR/W024985/1). B.J. is further supported by MRC Clinician Scientist Fellowship MR/Y00132X/1 and a Project Grant from Diabetes UK. The Section of Endocrinology at Imperial College London is funded by grants from the MRC, NIHR and is supported by the NIHR Biomedical Research Centre Funding Scheme and the NIHR/Imperial Clinical Research Facility. The views expressed are those of the author(s) and not necessarily those of the NHS, the NIHR or the Department of Health. G.A.R. was supported by a Wellcome Trust Investigator Award (WT212625/Z/18/Z), MRC Programme Grant (MR/R022259/1), NIH-NIDDK Project Grant (R01DK135268), CIHR-JDRF Team Grant (CIHR-IRSC TDP-186358 and JDRF 4-SRA-2023-1182-S-N), CRCHUM start-up funds, and an Innovation Canada John R. Evans Leader Award (CFI 42649). D.J.W. acknowledges support from the MRC (MC-A654-5QB40). S.J.M. was supported by a Wellcome Trust ISSF Fellowship (204834/Z/16/Z, award no. RSRO_67869), Society for Endocrinology Early Career and Small Equipment Grants and a Diabetes Research & Wellness Foundation Pump Priming Grant. D.W. and P.M.S. are Leadership Fellows of the National Health and Medical Research Council of Australia (NHMRC) (Grant IDs 2026300 and 2025694, respectively). Their work was also supported by an NHMRC Ideas grant to D.W. (ID:1184726).

## Supplementary Materials

Supplementary Methods, Figs. S1 to S9, Tables S1 to S9, Movie S1.

## Notes

### Competing Interest Statement

K.W.S. is an employee of Eli Lilly and Company and may own company stock. A.T. and B.J. have received Lilly Research Award Program (LRAP) support from Eli Lilly and Company.

### Summary of Updates

Changes in the manuscript text Changes in the Figures and Supplementary Information

