## Supplementary Information for "*In vivo* functional profiling and structural characterisation of the human *Glp1r* A316T variant"

### Supplementary Materials

### Supplementary Methods

#### *Nb37 bystander NanoBiT assays*

INS-1 832/3 *Glp1r*$^{-/-}$ cells were seeded in 6-well plates and co-transfected with the following plasmids: 200 ng pSNAP/FLAG-*hGlp1r*$^{WT}$ or $^{A316T}$, 500 ng G$\alpha_s$ (human, short isoform), 500 ng G$\beta$1 (human), 500 ng G$\gamma$2 (human), 100 ng RIC8B (human, isoform 2), and either 100 ng LgBiT-CAAX (KRAS CAAX motif N-terminally fused with LgBiT) or 500 ng Endofin-LgBiT (Endofin FYVE domain C-terminally fused with LgBiT), plus 100 ng (plasma membrane assay) or 500 ng (endosomal assay) SmBiT-Nb37 [Nb37 cDNA synthesised by GenScript C-terminally fused to SmBiT with a 15 amino acid flexible linker (*112*)] and adjusted for total DNA with pcDNA3.1 for the LgBiT-CAAX transfection. 24 hours after transfection, cells were exposed to NanoGlo Live Cell Reagent (Promega) with furimazine (1:20 dilution) and bioluminescence read in a Flexstation 3 plate reader for 8 minutes before and 30 minutes after stimulation by serial doses of up to 1 µM GLP-1. AUCs were calculated for each agonist concentration and fitted to log(agonist) *versus* response four-parameter curves using GraphPad Prism 10.1.2, with $logEC_{50}$ (potency) and $E_{max}$ (efficacy) calculated for each replicate dose response curve. Baseline endosomal over plasma membrane bioluminescence ratios were also calculated.

#### *GLP-1R conformational ECD opening time resolved-fluorescence resonance energy transfer (TR-FRET) assay*

INS-1 832/3 SNAP/FLAG-*hGlp1r*$^{WT}$ or $^{A316T}$ cells were labelled with SNAP-Lumi4-Tb (Revvity; 40 nM, 60 minutes at 37°C in complete medium). After washing to remove unbound probe, cells were resuspended in HBSS ± 100 nM plasma membrane dye NR12A (a gift from Prof Andrey Klymchenko, University of Strasbourg, France) for 5 minutes before washing again. Labelled cells were transferred to 96-well half-area white plates and baseline TR-FRET signal from Lumi-4Tb (donor) and NR12A (acceptor) recorded for 5 minutes at 37°C using a Flexstation 3 plate reader with the following settings: $\lambda_{ex}$ = 335 nm, $\lambda_{em}$ = 490 and 590 nm, delay 50 microseconds, integration time 300 microseconds. GLP-1 was then added at serial doses, and signal serially monitored for 30 minutes. The TR-FRET ratio, i.e. the ratio of fluorescence intensities at 590 and 490 nm, was considered indicative of the proximity of the GLP-1R ECD to the plasma membrane. Baseline-normalised AUCs were used to generate concentration–response curves fitted using GraphPad Prism 10.1.2, with $logEC_{50}$ and $E_{max}$ calculated for each replicate dose response curve.

***Real-time cytosolic calcium assay***

*hGlp1r*$^{-/-}$ mouse islets were transduced with pAV-SNAP/FLAG-*hGlp1r*$^{WT}$ or $^{A316T}$ adenoviruses for 24 hours, labelled with the $Ca^{2+}$ dye Cal-520 AM (4.5 µM; Stratech) diluted in KBRH buffer containing 6 mM glucose for 1 hour and encased in Matrigel onto 35-cm glass bottom MatTek dishes. Islets were then imaged every 6 seconds at 488 nm using a Nikon Eclipse Ti microscope with an ORCA-Flash 4.0 camera (Hamamatsu) and Metamorph software (Molecular Devices) at 37°C on a heated stage before and after addition of the indicated GLP-1RA followed by 11 mM glucose and then 20 mM KCl at the indicated time-points. Raw fluorescence intensity traces from islet ROIs were extracted using Fiji. Responses were plotted relative to the average fluorescence intensity during the 6 mM glucose baseline period, before agonist addition, and AUCs separately calculated for the GLP-1RA + 6 mM and 11 mM glucose stimulation periods.

***Isolation of detergent-resistant membrane (DRM) fractions***

To purify lipid rafts, DMR fractions were isolated from INS-1 832/3 SNAP/FLAG-*hGlp1r*$^{WT}$ or $^{A316T}$ cells by ultracentrifugation based on their insolubility in non-ionic detergents at 4°C and their low density because of their high lipid-to-protein ratio. Cells seeded onto 6-cm dishes were stimulated ± 100 nM exendin-4 or GLP-1 for 2 minutes, washed with ice-cold PBS and osmotically lysed in cold low salt 20 mM Tris-HCl, pH 7.0 buffer supplemented with cOmplete EDTA-free protease inhibitor cocktail (Roche) and phosphatase inhibitor cocktail 2 (Sigma-Aldrich) for 10-15 minutes on a rocking platform at 4°C. The cell suspension was collected and homogenised by passing through a 21-gauge needle, followed by ultracentrifugation at 63,000 g for 1 hour at 4°C with a Sorvall Discovery M120 ultracentrifuge. Pellets were resuspended in 200 µL ice-cold PBS supplemented with cOmplete EDTA-free protease inhibitor cocktail (Roche) and phosphatase inhibitor cocktail 2 and 1% Triton X-100, followed by rotating incubation for 30 minutes at 4°C. Detergent-treated membrane fractions were re-centrifuged at 63,000 g for 1 hour at 4°C. The supernatants, containing the 'disordered' detergent-soluble membrane (DSM) fractions, were retained for analysis, while the 'ordered' DRM fraction pellets were resuspended in 1% SDS supplemented with protease inhibitor and phosphatase inhibitor cocktails as above, sonicated with the Ultrasonic cell disruptor XL-2000 probe sonicator (Misonix, Inc) and centrifuged at 14,000 g for 5 minutes at 4 °C before Western blot analysis with anti-SNAP rabbit polyclonal antibody (1:500; P9310S, New England Biolabs) and goat anti-rabbit HRP secondary (1:2,000; ab6721, Abcam), stripping and reblotting with anti-flotillin-1 mouse monoclonal antibody (1:200; sc-74566, Santa Cruz Biotechnology) and sheep anti-mouse HRP secondary (1:5,000; ab6808, Abcam) as a DRM loading control. GLP-1R enrichment into DRMs was calculated from SNAP over flotillin band intensity levels under the different conditions tested using Fiji.

***Raster Image Correlation Spectroscopy (RICS) image capture and analysis***

INS-1 832/3 SNAP/FLAG-*hGlp1r*$^{WT}$ or $^{A316T}$ cells were seeded onto 35-cm glass bottom MatTek dishes and labelled with SNAP-Surface Alexa Fluor 647 for 10 minutes at 37°C in full media, washed and imaged in RMPI-1640 media without phenol red using a 100x oil objective on a Leica Stellaris 8 STED FALCON microscope from the Imperial College FILM Facility, using confocal settings. Cell were imaged and analysed as previously described (*113*), at the basal plasma membrane, both under vehicle conditions and following stimulation with 100 nM exendin-4 with a format size of 256 x 256 pixels and 80 nm pixel size for 200 consecutive frames. RICS analysis was carried out using the SimFCS 4 Software (Global

Software, G-SOFT Inc) to determine the diffusion coefficient of WT versus A316T GLP-1R in vehicle and exendin-4-stimulated conditions.

***GLP-1R ubiquitination assays***

INS-1 832/3 SNAP/FLAG-*hGlp1r*$^{WT}$ or $^{A316T}$ cells plated onto 6-cm dishes were stimulated ± 100 nM exendin-4 for 10 minutes, washed with PBS and osmotically lysed in lysis buffer (50 mM Tris-HCl, pH 7.4, 150 mM NaCl, 1% Triton X-100, 1 mM EDTA) with 1% protease inhibitor (Roche) and 1% phosphatase inhibitor cocktail (Sigma-Aldrich) at 4°C. Cell lysates were centrifuged at 12,000 g and 4°C for 10 minutes prior to the addition of 40 µL anti-FLAG M2 beads (A2220, Sigma-Aldrich) and overnight rotating incubation at 4°C to increase FLAG-tag binding efficiency. Samples were then centrifuged at 5,000 g for 30 seconds, bead pellets washed 3x in 500 µL wash buffer (50 mM Tris-HCl, 150 mM NaCl, pH 7.4), and proteins eluted with 2x urea buffer at 37°C for 10 minutes before Western blot analysis with anti-ubiquitin antibody (1:1,000; sc-8017, Santa Cruz Biotechnology) and sheep anti-mouse HRP secondary (1:5,000; ab6808, Abcam) followed by stripping and reblotting with anti-SNAP antibody as above to assess immunoprecipitated receptor per sample. GLP-1R ubiquitination was calculated from ubiquitin over SNAP band intensity levels under the different conditions tested using Fiji.

***LC-MS/MS GLP-1R interactome analysis***

INS-1 832/3 SNAP/FLAG-*hGlp1r*$^{WT}$ or $^{A316T}$ cells were plated onto 6-cm dishes and stimulated with vehicle or 100 nM GLP-1RA for 5 minutes as previously described (*114*). Following stimulation, GLP-1R was immunoprecipitated with anti-FLAG M2 beads as above and bead pellets washed 5x in wash buffer (50 mM Tris-HCl, 150 mM NaCl, pH 7.4), before storage at -80°C until further analysis. Beads were resuspended in 150 µL urea 1 M, 20 mM HEPES, 1 mM DTT, pH 8.0 followed by addition of 1.5 µg trypsin gold (Promega) and overnight digestion at 37°C. Supernatants were subsequently acidified with 0.5% trifluoroacetic acid (TFA), followed by desalting using reversed-phase spin tips (Glygen Corp) and dried using vacuum centrifugation. Dried peptides were resuspended in 0.1% TFA and Liquid

Chromatography Tandem Mass Spectrometry (LC-MS/MS) performed as previously described (*115*) using an Ultimate 3000 nano-HPLC coupled to a Q-Exactive mass spectrometer (Thermo Scientific) via an EASY-Spray source. Peptides were loaded onto a trap column (Acclaim PepMap 100 C18, 100 µm x 2 cm) at 8 µL/min in 2% acetonitrile, 0.1% TFA. Peptides were eluted on-line to an analytical column (Acclaim Pepmap RSLC C18, 75 µm x 75 cm). A 90-minute stepped gradient separation was used with 4-25% acetonitrile 80%, formic acid 0.1% for 60 minutes, followed by 25-45% for 30 minutes. Eluted peptides were analysed by Q-Exactive operating in positive polarity and data-dependent acquisition mode. Ions for fragmentation were determined from an initial MS1 survey scan at 70,000 resolution (at m/z 200), followed by higher-energy collisional dissociation of the top 12 most abundant ions at a resolution of 17,500. MS1 and MS2 scan AGC targets were set to 3e6 and 5e4 for a maximum injection time of 50 and 75 milliseconds, respectively. A survey scan m/z range of 350 – 1800 m/z was used, with a normalised collision energy set to 27%, minimum AGC of 1e3, charge state exclusion enabled for unassigned and +1 ions and a dynamic exclusion of 45 seconds. Data was processed using the MaxQuant software platform (v1.6.10.43) (*116*), with database searches carried out by the in-built Andromeda search engine against the Swissprot Rattus Norvegicus database (downloaded – 21$^{st}$ May 2022, entries: 8,132) concatenated with the human GLP-1R protein sequence. A reverse decoy database approach was used at a 1% FDR for peptide spectrum matches and protein identifications. Search parameters included: maximum missed cleavages set to 2, variable modifications of methionine oxidation, protein N-terminal acetylation, asparagine deamidation and cyclisation of glutamine to pyroglutamate. Label-free quantification was enabled with a label-free quantitation (LFQ) minimum ratio count of 1. Proteins were filtered for those containing at least one unique peptide and identified in all biological replicates. For normalisation, proteins were ranked by fold change increase in abundance compared with control (vehicle-treated) immunoprecipitates. The resulting data was pre-processed by elimination of proteins that are non-specifically bound and residual background proteins, such as common contaminants and reverse database proteins. To further facilitate the selection of GLP-1R interactors, the following criteria were established; i) identified at least twice from four runs; ii) not nuclear proteins; and (iii) previously identified in pancreatic β-cells. Untargeted LFQ intensity values, normalised to hGLP-1R levels in each immunoprecipitated sample, were analysed by LFQ Analyst (*117*) to identify differences in levels of individual protein - GLP-1R interactions between vehicle and GLP-1RA treated conditions. Paired t-tests adjusted for multiple comparisons using Benjamin-Hochberg false discovery rate correction and Perseus type imputations were used for statistical analysis.

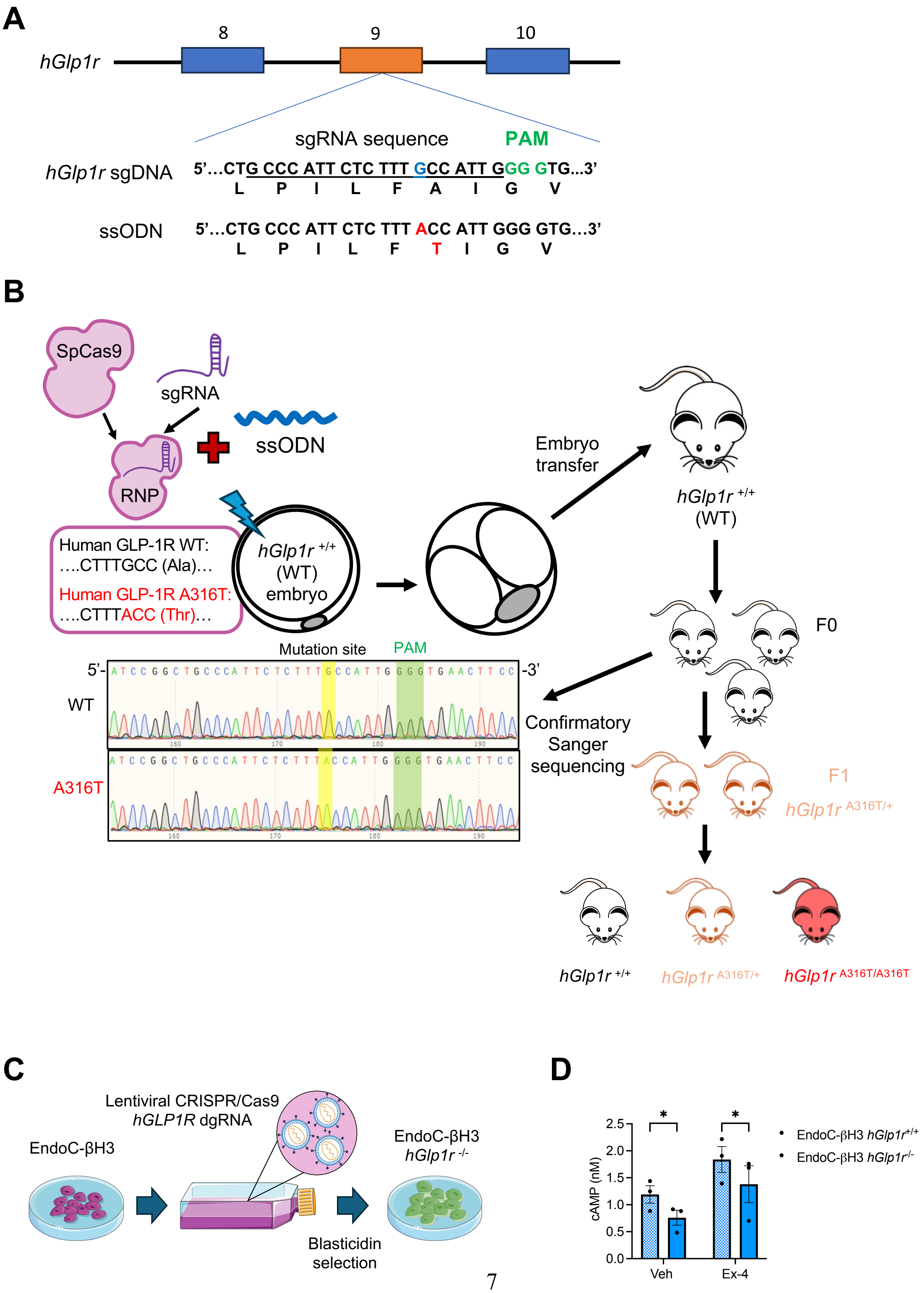
A
hGlp1r
8
9
10
sgRNA sequence
PAM
hGlp1r sgDNA
5'…CTG CCC ATT CTC TTT GCC ATT GGG GTG…3'
L P I L F A I G V
ssODN
5'…CTG CCC ATT CTC TTT ACC ATT GGG GTG…3'
L P I L F T I G V
B
SpCas9
sgRNA
ssODN
RNP
Human GLP-1R WT:
….CTTTGCC (Ala)…
Human GLP-1R A316T:
….CTTTACC (Thr)…
hGlp1r +/+ (WT) embryo
Embryo transfer
hGlp1r +/+ (WT)
F0
Mutation site
PAM
5'-
-3'
WT
A316T
Confirmatory Sanger sequencing
F1
hGlp1r A316T/+
hGlp1r +/+
hGlp1r A316T/+
hGlp1r A316T/A316T
C
Lentiviral CRISPR/Cas9
hGLP1R dgRNA
EndoC-βH3
EndoC-βH3
hGlp1r -/-
Blasticidin selection
D
cAMP (nM)
Veh
Ex-4
EndoC-βH3 hGlp1r+/+
EndoC-βH3 hGlp1r-/-

**Fig. S1. Model generation.** (**A**) Targeting strategy for *hGlp1r*$^{A316T/A316T}$ knock-in mouse generation; CRISPR single guide RNA (sgRNA) targeting the Cas9 nuclease to the indicated PAM region of exon 9 in the human *Glp1r* gene and corresponding ssODN homology directed repair template sequences are indicated. (**B**) Schematic diagram of the method to generate *hGlp1r*$^{A316T/A316T}$ knock-in mice: electroporation of Cas9, sgRNA and ssODN templates into *hGlp1r*$^{+/+}$ pronucleus, followed by IVF transfer of two cell embryo into *hGlp1r*$^{+/+}$ female mice resulting in F0 generation and screening via genomic PCR and Sanger sequencing, with sequences from *hGlp1r*$^{+/+}$ and $^{A316T/A316T}$ mice included. (**C**) Schematic diagram of the method to generate EndoC-βH3 *hGlp1r*$^{-/-}$ cells: transduction with engineered lentiviral CRISPR/Cas9 double guide RNA (dgRNA) vector targeting *hGlp1r* followed by blasticidin selection. (**D**) Static cAMP levels from *EndoC-βH3 hGlp1r*$^{+/+}$ *versus* $^{-/-}$ cells under vehicle (Veh) and 30 minutes exendin-4 (Ex-4) stimulation.

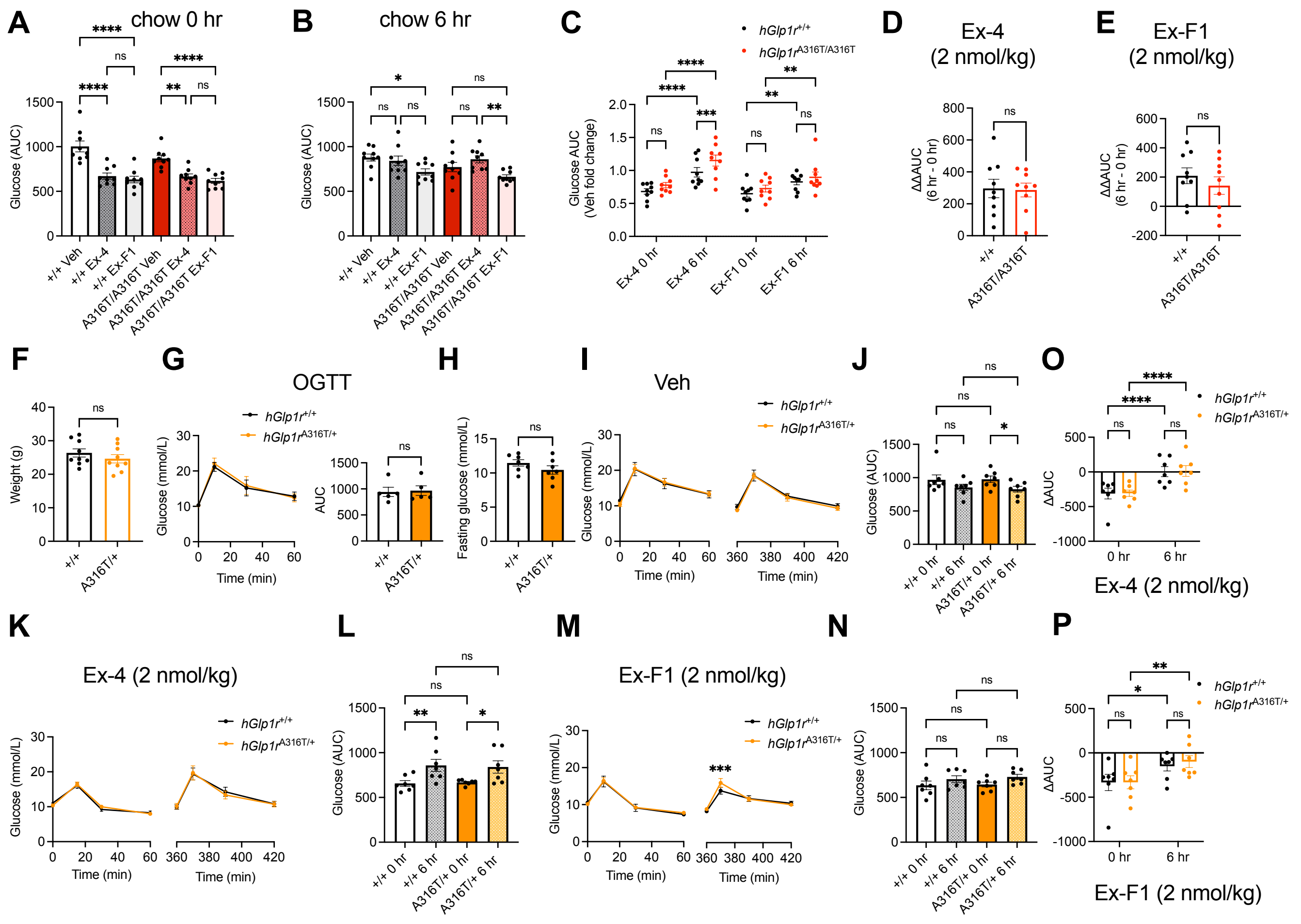


**Fig. S2. *In vivo* glucose responses of adult *hGlp1r*$^{A316T/A316T}$ and *hGlp1r*$^{A316T/+}$ *versus hGlp1r*$^{+/+}$ mice on a chow diet – extra data.** (**A**) AUCs from acute IPGTTs taken from Fig. 1E, G and I. (**B**) As in (A) for IPGTTs 6 hours post-vehicle or agonist administration. (**C**) Vehicle fold AUC responses calculated from Fig. 1E, G and I. (**D**) Exendin-4 (Ex-4) ΔΔAUC responses (6 hours minus acute) calculated from Fig. 1J. (**E**) Exendin-F1 (Ex-F1) ΔΔAUC responses (6 hours minus acute) calculated from Fig. 1K. (**F**) Age-matched body weight of *hGlp1r*$^{+/+}$ *versus* $^{A316T/+}$ mixed sex littermate adult mice ($n$ = 9 per genotype) on a chow diet. (**G**) OGTT responses and corresponding AUCs following 2 g/kg body weight glucose administration via oral gavage after 5 hours of fasting in a mixed sex cohort of lean, adult *hGlp1r*$^{+/+}$ *versus* $^{A316T/+}$ mice on a chow diet ($n$ = 5 per genotype). (**H**) Glucose levels after 2 hours of fasting in chow fed *hGlp1r*$^{+/+}$ *versus* $^{A316T/+}$ mixed sex adult mice ($n$ = 7 per genotype). (**I** and **J**) IPGTTs (2 g/kg body weight glucose) following acute (0 hours), or 6 hours post-intraperitoneal administration of saline vehicle (Veh) in chow-fed *hGlp1r*$^{+/+}$ *versus* $^{A316T/+}$ mixed sex adult mice. Glucose curves (I) and corresponding AUCs (J) shown; *n* = 7 per genotype. (**K** to **N**) As in (**I** and **J**) following administration of 2 nmol/kg exendin-4 (Ex-4) (**K** and **L**) or exendin-F1 (Ex-F1) (**M** and **N**). (**O** and **P**) Vehicle-corrected ΔAUC responses from (L) and (N), respectively. Data are mean ± SEM; *p<0.05; **p<0.01; ***p< 0.001; ****p<0.0001; ns, not significant by paired t-tests, one- or two-way ANOVA with Sidak's post-hoc tests.

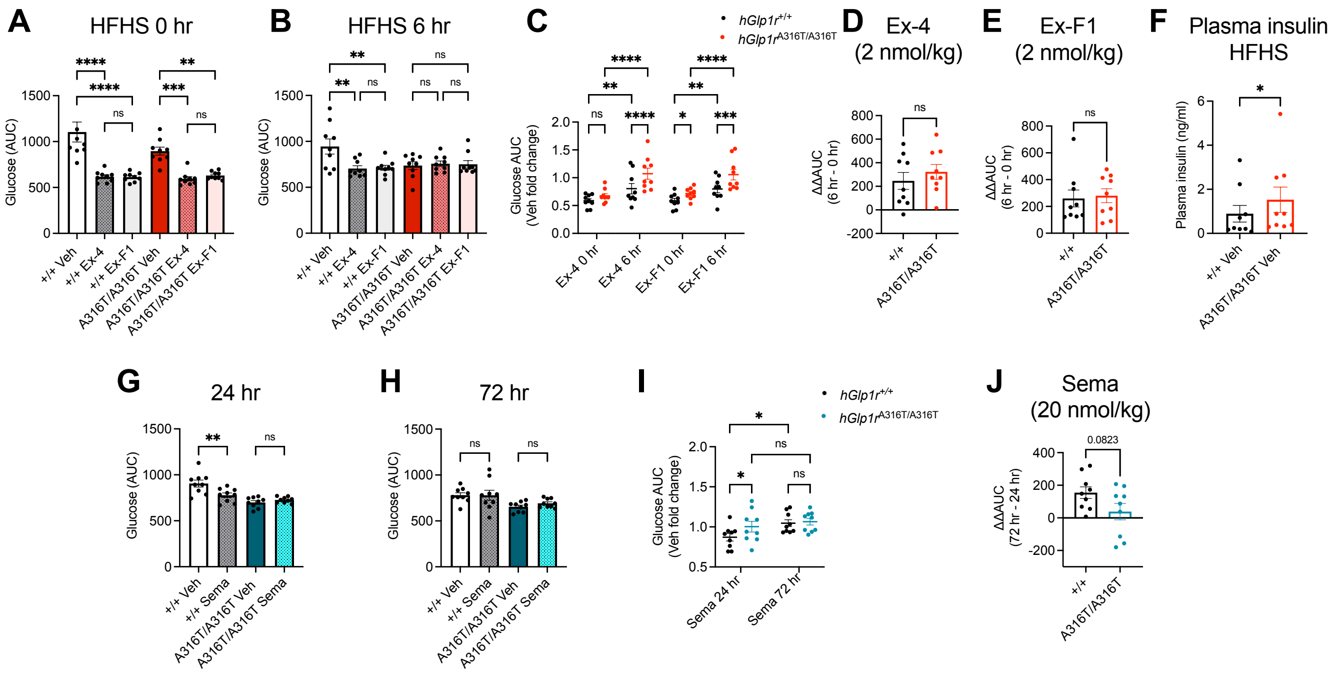


**Fig. S3.** ***In vivo*** **glucose responses of adult** ***hGlp1r***$^{\mathbf{A316T/A316T}}$ ***versus hGlp1r***$^{\mathbf{+/+}}$ **mice on a HFHS diet – extra data.** (**A**) AUCs from acute IPGTTs taken from Fig. 2F, H and J. **b**, As in (A) for IPGTTs at 6 hours post-vehicle or agonist administration. (**C**) Vehicle fold AUC responses calculated from Fig. 2F, H and J. (**D**) Exendin-4 (Ex-4) ΔΔAUC responses (6 hours minus acute) calculated from Fig. 2K. (**E**) Exendin-F1 (Ex-F1) ΔΔAUC responses (6 hours minus acute) calculated from Fig. 2L. (**F**) Acute plasma insulin levels post-vehicle (Veh) administration in HFHS-fed *hGlp1r*$^{+/+}$ *versus* $^{A316T/A316T}$ mixed sex adult mice. (**G**) AUCs from IPGTTs at 24 hours post-vehicle (Veh) or agonist administration taken from Fig. 2N and P. (**H**) As in (G) for IPGTTs at 72 hours post-vehicle (Veh) or agonist administration. (**I**) Vehicle (Veh) fold AUC responses calculated from Fig. 2N and P. (**J**) Semaglutide (Sema) ΔΔAUC responses (72 hours minus 24 hours) calculated from Fig. 2Q. Data are mean ± SEM; *p<0.05; **p<0.01; ***p< 0.001; ****p<0.0001; ns, not significant by paired or ratio paired t-tests, one- or two-way ANOVA with Sidak's post-hoc tests.

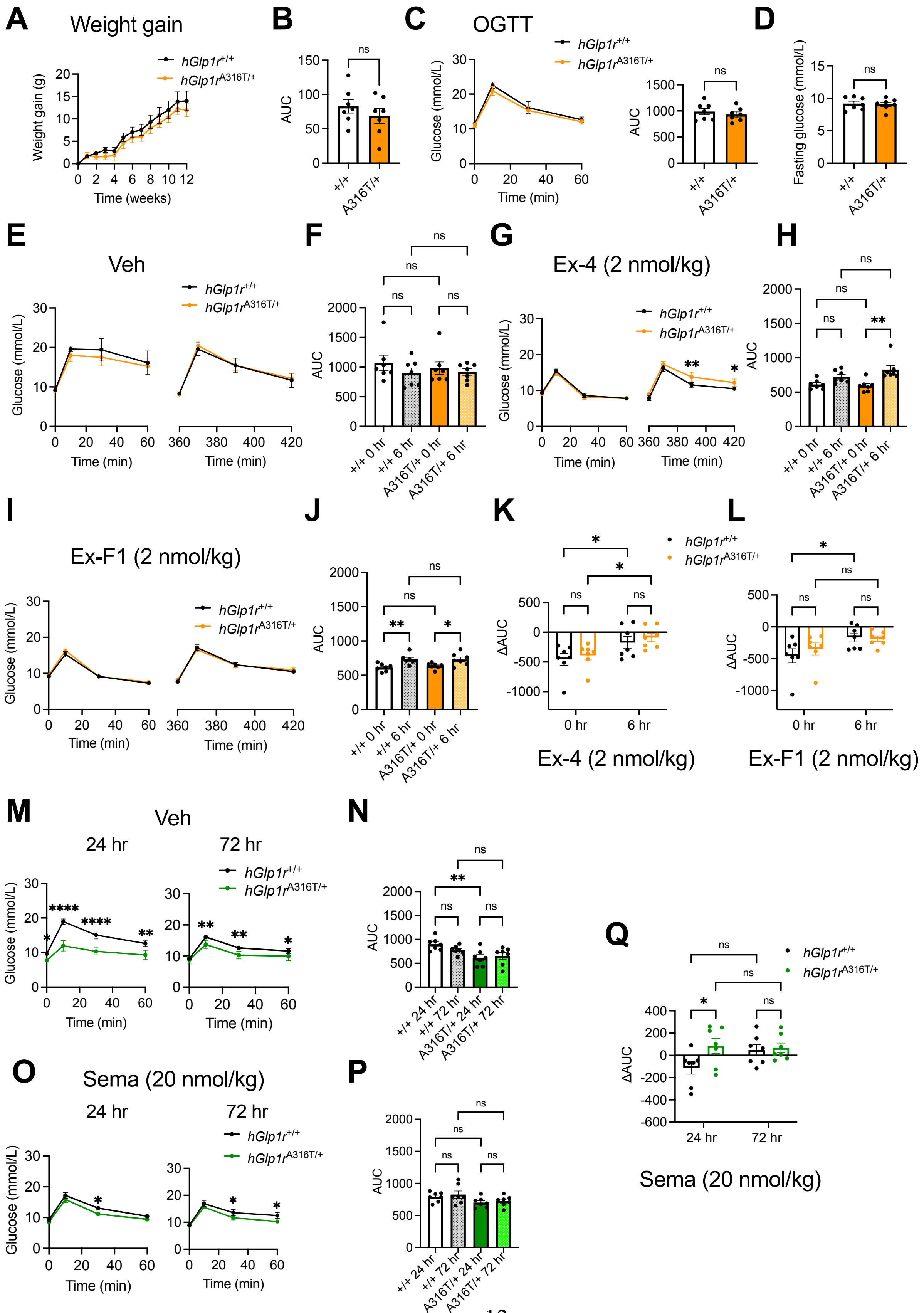

A
Weight gain
hGlp1r+/+
hGlp1rA316T/+
Weight gain (g)
Time (weeks)
B
AUC
ns
+/+
A316T/+
C
OGTT
hGlp1r+/+
hGlp1rA316T/+
Glucose (mmol/L)
Time (min)
AUC
ns
D
Fasting glucose (mmol/L)
ns
E
Veh
hGlp1r+/+
hGlp1rA316T/+
Glucose (mmol/L)
Time (min)
F
AUC
ns
+/+ 0 hr
+/+ 6 hr
A316T/+ 0 hr
A316T/+ 6 hr
G
Ex-4 (2 nmol/kg)
hGlp1r+/+
hGlp1rA316T/+
Glucose (mmol/L)
Time (min)
**
*
H
AUC
ns
**
I
Ex-F1 (2 nmol/kg)
hGlp1r+/+
hGlp1rA316T/+
Glucose (mmol/L)
Time (min)
J
AUC
ns
**
*
K
ΔAUC
hGlp1r+/+
hGlp1rA316T/+
0 hr
6 hr
Ex-4 (2 nmol/kg)
L
ΔAUC
0 hr
6 hr
Ex-F1 (2 nmol/kg)
M
Veh
24 hr
72 hr
hGlp1r+/+
hGlp1rA316T/+
Glucose (mmol/L)
Time (min)
****
**
*
N
AUC
ns
**
+/+ 24 hr
+/+ 72 hr
A316T/+ 24 hr
A316T/+ 72 hr
O
Sema (20 nmol/kg)
24 hr
72 hr
hGlp1r+/+
hGlp1rA316T/+
Glucose (mmol/L)
Time (min)
P
AUC
ns
Q
hGlp1r+/+
hGlp1rA316T/+
ΔAUC
24 hr
72 hr
Sema (20 nmol/kg)

**Fig. S4.** ***In vivo*** **glucose responses of adult** ***hGlp1r***$^{\mathbf{A316T/+}}$ ***versus hGlp1r***$^{\mathbf{+/+}}$ **mice on a HFHS diet.** (**A**) Body weight gain for *hGlp1r*$^{+/+}$ *versus* $^{A316T/+}$ mixed sex littermate adult mice (*n* = 7 per genotype) maintained on a HFHS diet for the indicated times. (**B**) AUC from (A). (**C**) OGTT responses and corresponding AUC following 2 g/kg body weight glucose administration via oral gavage after 5 hours of fasting in a mixed sex cohort of adult *hGlp1r*$^{+/+}$ *versus* $^{A316T/+}$ mice on a HFHS diet (*n* = 7 per genotype). (**D**) Glucose levels after 2 hours of fasting from mice in (A). (**E** and **F**) IPGTTs (2 g/kg body weight glucose) following acute (0 hours), or 6 hours post-intraperitoneal administration of saline vehicle (Veh) in HFHS-fed *hGlp1r*$^{+/+}$ *versus* $^{A316T/+}$ mixed sex adult mice. Glucose curves (E) and corresponding AUCs (F) shown; *n* = 7 mice per genotype. (**G** to **J**) As in (**E** and **F**) following administration of 2 nmol/kg exendin-4 (Ex-4) (**G** and **H**) or exendin-F1 (Ex-F1) (**I** and **J**). (**K** and **L**) Vehicle-corrected ΔAUC responses from (H) and (J), respectively. (**M** to **R**) Glucose curves and corresponding AUCs following vehicle (Veh) (M and N) or 20 nmol/kg semaglutide (Sema) (O and P) treatment of HFHS-fed *hGlp1r*$^{+/+}$ *versus* $^{A316T/+}$ mice; IPGTTs performed 24- and 72-hours post-agonist administration; *n* = 7 mice per genotype. (**Q**) Vehicle-corrected ΔAUC responses from (P). Data are mean ± SEM; *p<0.05; **p<0.01; ****p<0.0001; ns, not significant by paired t-tests, one- or two-way ANOVA with Sidak's post-hoc tests.

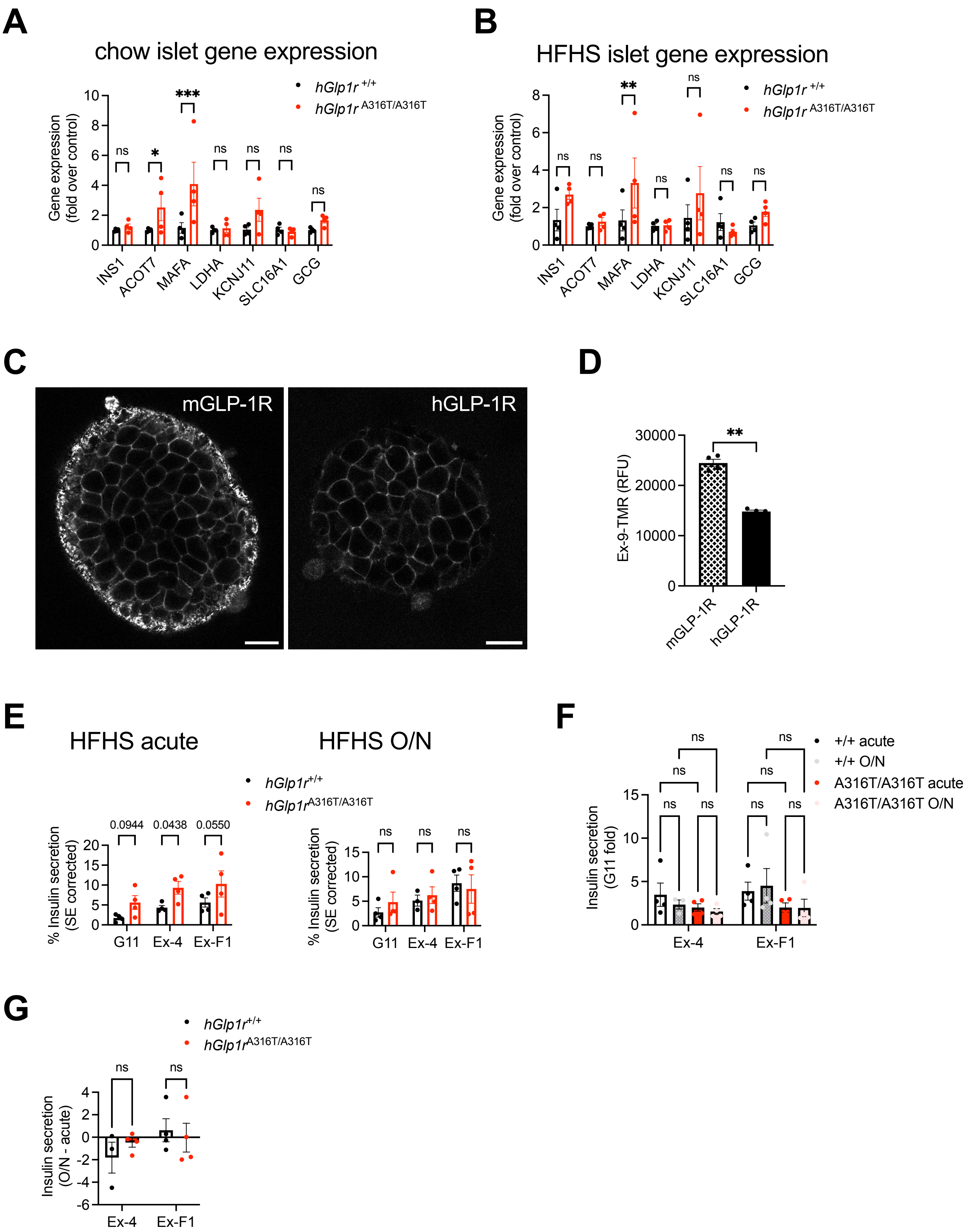
A
chow islet gene expression
hGlp1r +/+
hGlp1r A316T/A316T
Gene expression (fold over control)
INS1
ACOT7
MAFA
LDHA
KCNJ11
SLC16A1
GCG
ns
*
***
B
HFHS islet gene expression
**
C
mGLP-1R
hGLP-1R
D
Ex-9-TMR (RFU)
30000
20000
10000
0
mGLP-1R
hGLP-1R
E
HFHS acute
HFHS O/N
% Insulin secretion (SE corrected)
0.0944
0.0438
0.0550
G11
Ex-4
Ex-F1
F
+/+ acute
+/+ O/N
A316T/A316T acute
A316T/A316T O/N
Insulin secretion (G11 fold)
G
Insulin secretion (O/N - acute)

**Fig. S5. GLP-1R-induced downstream signalling in *hGlp1r*$^{A316T/A316T}$ *versus hGlp1r*$^{+/+}$ mouse islets – extra data.** (**A** and **B**) Relative gene expression of β-cell enriched and disallowed genes in islets purified from *hGlp1r*$^{A316T/A316T}$ *versus hGlp1*$^{+/+}$ mice fed a chow (A) or HFHS (B) diet; size bars, 20 μm; *n* = 4. (**C**) Representative confocal microscopy images of islets from *hGlp1r versus mGlp1r* mice treated for 10 minutes with 100 nM exendin-9-TMR. (**D**) Quantification of exendin-9-TMR (Ex-9-TMR) fluorescence intensity in islets from (C); *n* = 4. (**E**) Percentage of acute and overnight (O/N) surface expression (SE)-corrected insulin secretion in HFHS-fed *hGlp1r*$^{+/+}$ *versus* $^{A316T/A316T}$ mouse islets stimulated with 11 mM glucose (G11) alone or supplemented with 100 nM exendin-4 (Ex-4) or exendin-F1 (Ex-F1); *n* = 4. (**F**) Insulin secretion fold changes to G11 calculated from data in (E). (**G**) Overnight over acute insulin secretion responses calculated from data in (F). Data are mean ± SEM; *p<0.05; **p<0.01; ***p<0.001; ns, not significant by paired t-tests, or two-way ANOVA with Sidak's post-hoc tests.

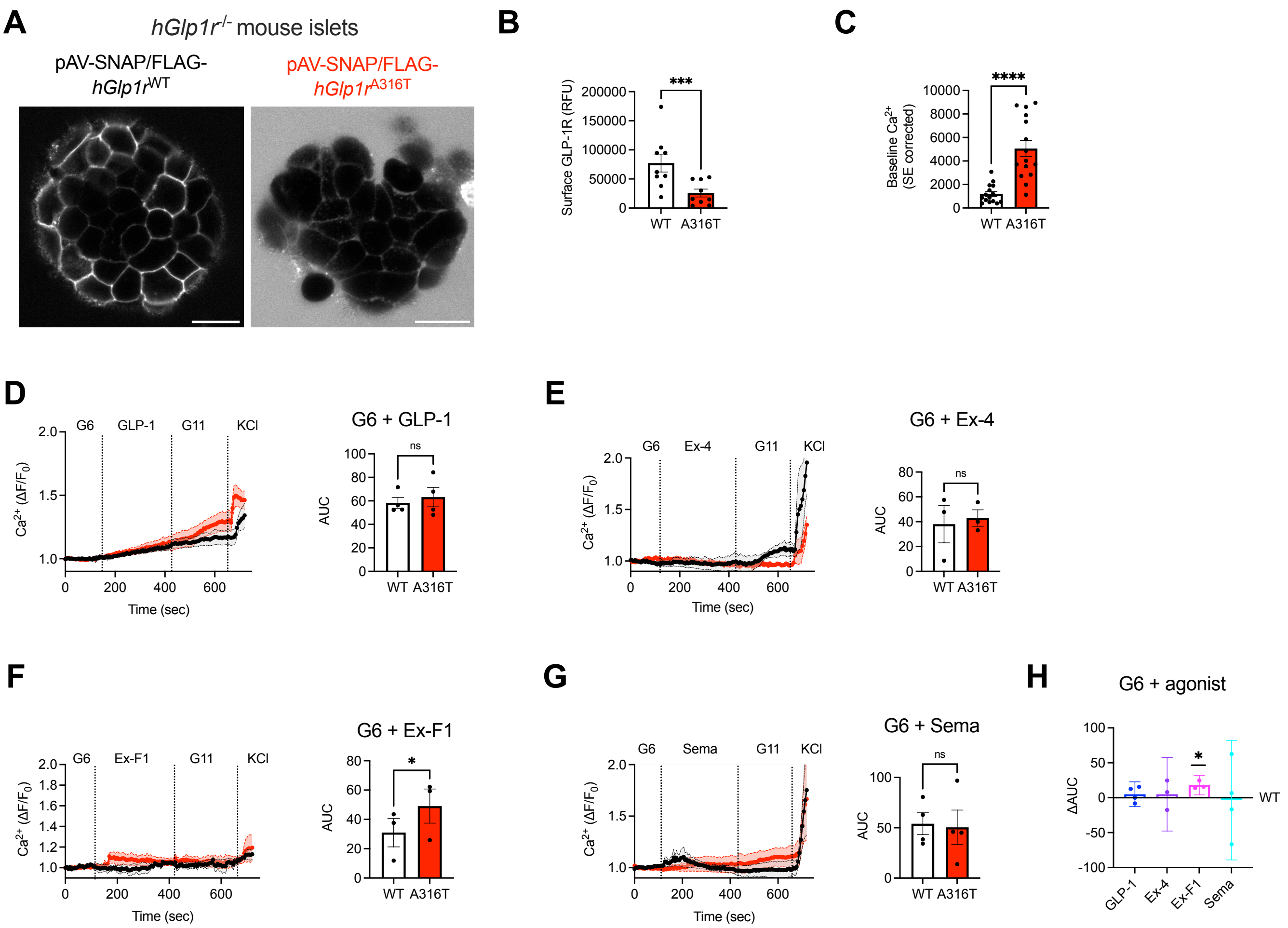


**Fig. S6. GLP-1R surface expression and calcium signalling in *hGlp1r*$^{-/-}$ mouse islets transduced with SNAP/FLAG-*hGlp1r*$^{WT}$ *versus* $^{A316T}$ adenoviruses.** (**A** and **B**) Surface expression in islets from *hGlp1r*$^{-/-}$ mice transduced with pAV-SNAP/FLAG-*hGlp1r*$^{WT}$ or $^{A316T}$ labelled with SNAP-Surface Alexa Fluor 647. Representative images (A) and quantification of surface GLP-1R (B); size bars, 20 μm; *n* = 9. (**C**) Basal $Ca^{2+}$ levels in Cal-520 AM-loaded *hGlp1r*$^{-/-}$ mouse islets transduced with pAV-SNAP/FLAG-*hGlp1r*$^{WT}$ or$^{A316T}$; *n* = 15. (**D** to **G**) $Ca^{2+}$ responses from islets from (C) stimulated with 100 nM GLP-1 (D), exendin-4 (Ex-4) (E), exendin-F1 (Ex-F1) (F), or semaglutide (Sema) (G), followed by 20 mM KCl for maximal responses; $Ca^{2+}$ traces and corresponding AUCs for agonist responses at 6 mM glucose (G6) shown; *n* = 3-4. (**H**) ΔAUC (SNAP/FLAG-*hGlp1r*$^{A316T}$ minus $^{WT}$) responses calculated from data in

(D to G). Data are mean ± SEM except for ΔAUC *hGlp1r*$^{A316T}$ minus $^{WT}$ responses which are mean ± 95% confidence interval (CI); *p<0.05; ***p<0.001; ns, not significant by paired or ratio paired t-tests.

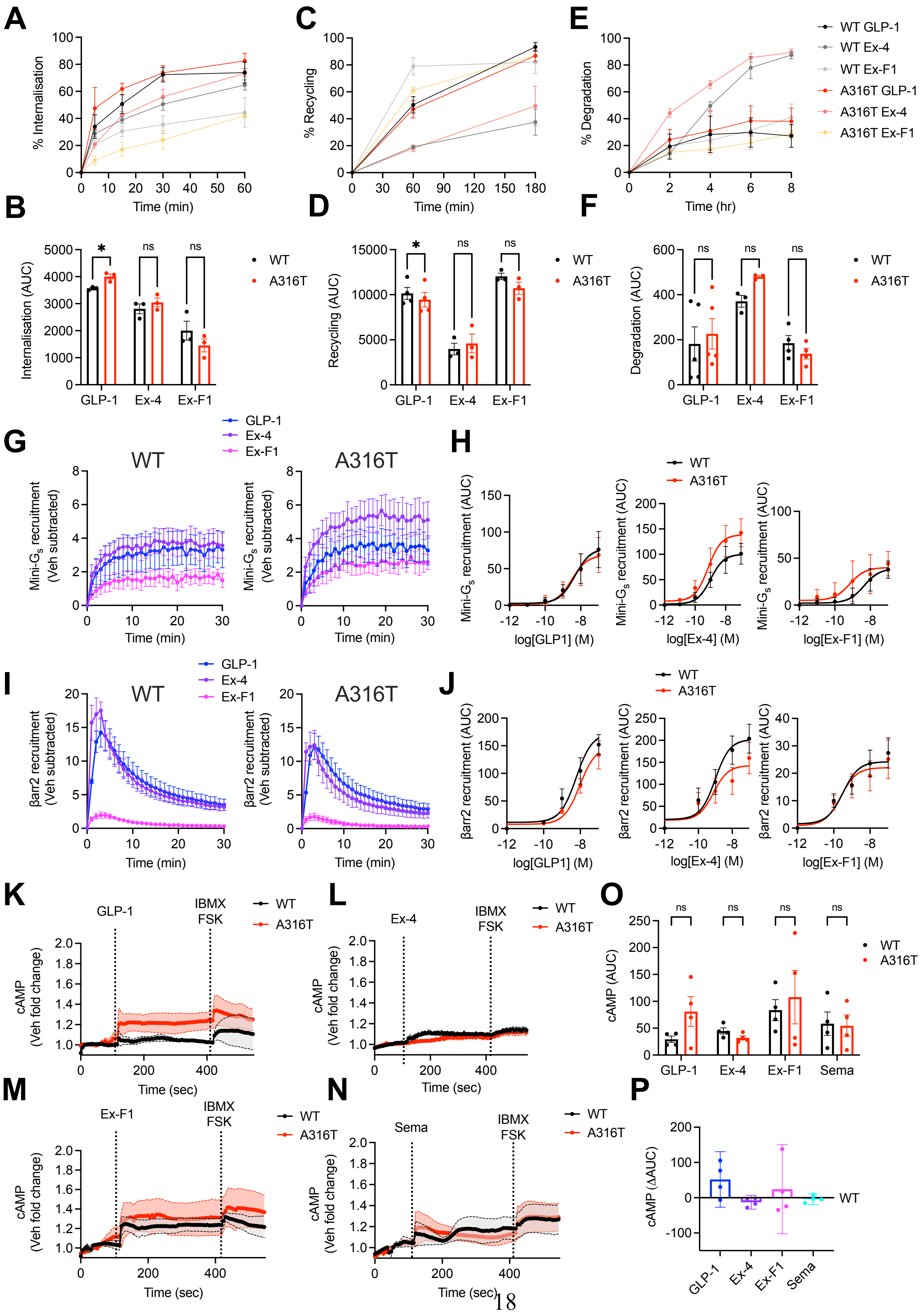
A
% Internalisation
Time (min)
C
% Recycling
Time (min)
E
% Degradation
Time (hr)
WT GLP-1
WT Ex-4
WT Ex-F1
A316T GLP-1
A316T Ex-4
A316T Ex-F1
B
Internalisation (AUC)
GLP-1
Ex-4
Ex-F1
WT
A316T
D
Recycling (AUC)
F
Degradation (AUC)
G
WT
A316T
Mini-Gs recruitment (Veh subtracted)
H
Mini-Gs recruitment (AUC)
log[GLP1] (M)
log[Ex-4] (M)
log[Ex-F1] (M)
I
βarr2 recruitment (Veh subtracted)
J
βarr2 recruitment (AUC)
K
GLP-1
IBMX FSK
cAMP (Veh fold change)
Time (sec)
L
Ex-4
O
cAMP (AUC)
Sema
M
Ex-F1
N
Sema
P
cAMP (ΔAUC)

**Fig. S7. Functional characterisation of INS-1 832/3 *hGlp1r*$^{-/-}$ SNAP/FLAG-*hGlp1r*$^{WT}$ *versus* $^{A316T}$ sublines – extra data.** (**A**) Percentage of GLP-1R internalisation over time following stimulation with the indicated agonists in INS-1 832/3 *hGlp1r*$^{-/-}$ SNAP/FLAG-*hGlp1r*$^{WT}$ *versus* $^{A316T}$ cells; *n* = 3. (**B**) Internalisation AUCs calculated from (A). (**C**) Percentage of GLP-1R recycling to the plasma membrane over time following stimulation with the indicated agonists in INS-1 832/3 *hGlp1r*$^{-/-}$ SNAP/FLAG-*hGlp1r*$^{WT}$ *versus* $^{A316T}$ cells; *n* = 3. (**D**) Recycling AUCs calculated from (C). (**E**) Percentage of GLP-1R degradation over time following stimulation with the indicated agonists in INS-1 832/3 *hGlp1r*$^{-/-}$ SNAP/FLAG-*hGlp1r*$^{WT}$ *versus* $^{A316T}$ cells; *n* = 3. (**F**) Degradation AUCs calculated from (E). (**G**) Vehicle-subtracted LgBiT-mini-$G_s$ recruitment to *hGlp1r*$^{WT}$ or $^{A316T}$-SmBiT over time in INS-1 832/3 *hGlp1r*$^{-/-}$ cells following stimulation with 100 nM of the indicated agonist; *n* = 5. (**H**) LgBiT-mini-$G_s$ recruitment to *hGlp1r*$^{WT}$ or $^{A316T}$-SmBiT AUC dose responses to the indicated agonist; *n* = 5. (**I**) Vehicle-subtracted LgBiT-β-arrestin 2 (βarr2) recruitment to *hGlp1r*$^{WT}$ or $^{A316T}$-SmBiT over time in INS-1 832/3 *hGlp1r*$^{-/-}$ cells following stimulation with 100 nM of the indicated agonist; *n* = 5. (**J**) LgBiT-βarr2 recruitment to *hGlp1r*$^{WT}$ or $^{A316T}$-SmBiT AUC dose responses to the indicated agonist; *n* = 5. (**K** to **N**) cAMP responses from cADDis transduced INS-1 832/3 *hGlp1r*$^{-/-}$ SNAP/FLAG-*hGlp1r*$^{WT}$ *versus* $^{A316T}$ cells in response to 100 nM GLP-1 (K), exendin-4 (Ex-4) (L), exendin-F1 (Ex-F1) (M), or semaglutide (Sema) (N), followed by IBMX plus forskolin (FSK) stimulation for maximal responses; *n* = 4. (**O** and **P**) cAMP AUCs (O) or ΔAUC (SNAP/FLAG-*hGlp1r*$^{A316T}$ minus $^{WT}$) (P) calculated from agonist-stimulated periods in (K to N); *n* = 4. Data are mean ± SEM except for ΔAUC *hGlp1r* $^{A316T}$ minus $^{WT}$ responses which are mean ± 95% confidence interval (CI); *p<0.05; ns, not significant by two-way ANOVA with Sidak's post-hoc tests.

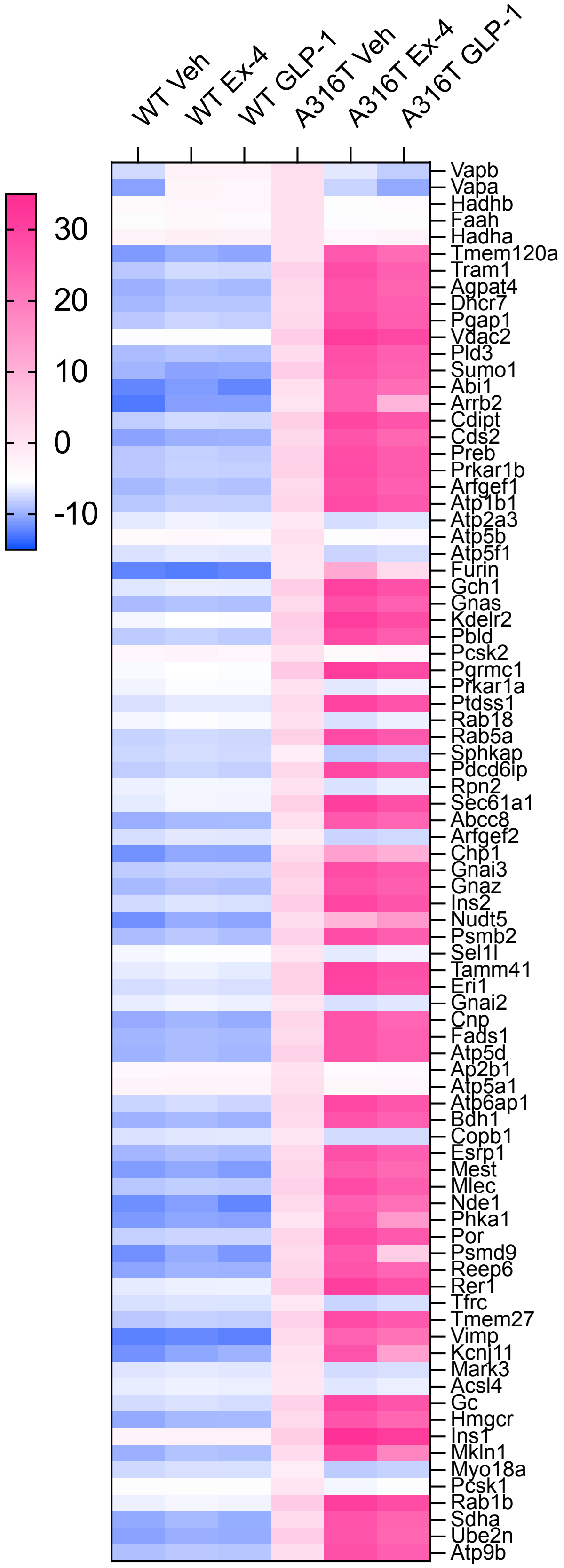
WT Veh
WT Ex-4
WT GLP-1
A316T Veh
A316T Ex-4
A316T GLP-1
30
20
10
0
-10
Vapb
Vapa
Hadhb
Faah
Hadha
Tmem120a
Tram1
Agpat4
Dhcr7
Pgap1
Vdac2
Pld3
Sumo1
Abi1
Arrb2
Cdipt
Cds2
Preb
Prkar1b
Arfgef1
Atp1b1
Atp2a3
Atp5b
Atp5f1
Furin
Gch1
Gnas
Kdelr2
Pbld
Pcsk2
Pgrmc1
Prkar1a
Ptdss1
Rab18
Rab5a
Sphkap
Pdcd6ip
Rpn2
Sec61a1
Abcc8
Arfgef2
Chp1
Gnai3
Gnaz
Ins2
Nudt5
Psmb2
Sel1l
Tamm41
Eri1
Gnai2
Cnp
Fads1
Atp5d
Ap2b1
Atp5a1
Atp6ap1
Bdh1
Copb1
Esrp1
Mest
Mlec
Nde1
Phka1
Por
Psmd9
Reep6
Rer1
Tfrc
Tmem27
Vimp
Kcnj11
Mark3
Acsl4
Gc
Hmgcr
Ins1
Mkln1
Myo18a
Pcsk1
Rab1b
Sdha
Ube2n
Atp9b

**Fig. S8. GLP-1R WT *versus* A316T interactome analysis in INS-1 832/3 *hGlp1r*$^{-/-}$ SNAP/FLAG-*hGlp1r*$^{WT}$ *versus* $^{A316T}$ cells.** Heatmap of GLP-1R interactor enrichment from LC-MS/MS analysis of INS-1 832/3 *hGlp1r*$^{-/-}$ SNAP/FLAG-*hGlp1r*$^{WT}$ *versus* $^{A316T}$ anti-FLAG co-immunoprecipitates under vehicle (Veh) or 5 minutes stimulation with 100 nM exendin-4 (Ex-4) or GLP-1; blue, decreased; pink, increased interactor enrichment; LC-MS/MS data analysed in LFQ-Analyst and normalised to hGLP-1R levels for each experimental repeat; colour scale depicting $\log_2$ fold change to WT vehicle centred around overall median value of the experiment; $n$ = 2-4.

**A**

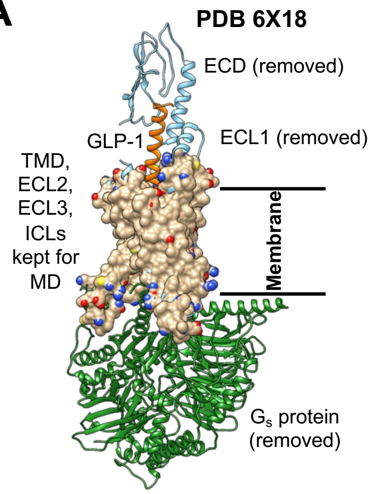


**B**

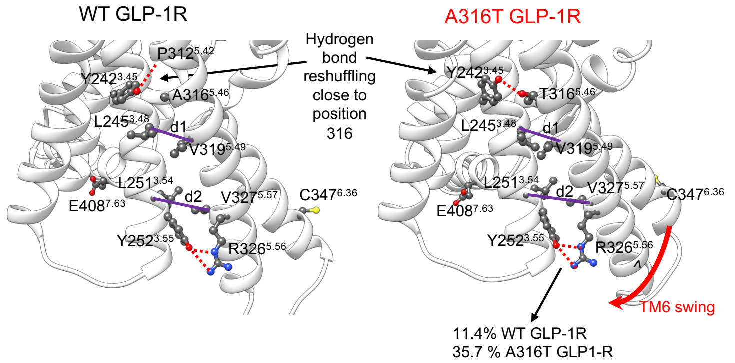


**C**

| Distance | WT GLP-1R | A316T GLP-1R |
|---|---|---|
| L245$^{3.48}$-V319$^{5.49}$ (**d1**) | 7.58 ± 0.28 Å | 8.09 ± 0.34 Å |
| L251$^{3.54}$-V327$^{5.57}$ (**d2**) | 8.91 ± 0.68 Å | 7.75 ± 0.48 Å |

**D**

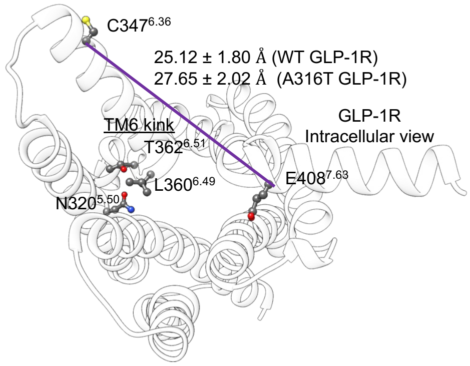


**E**

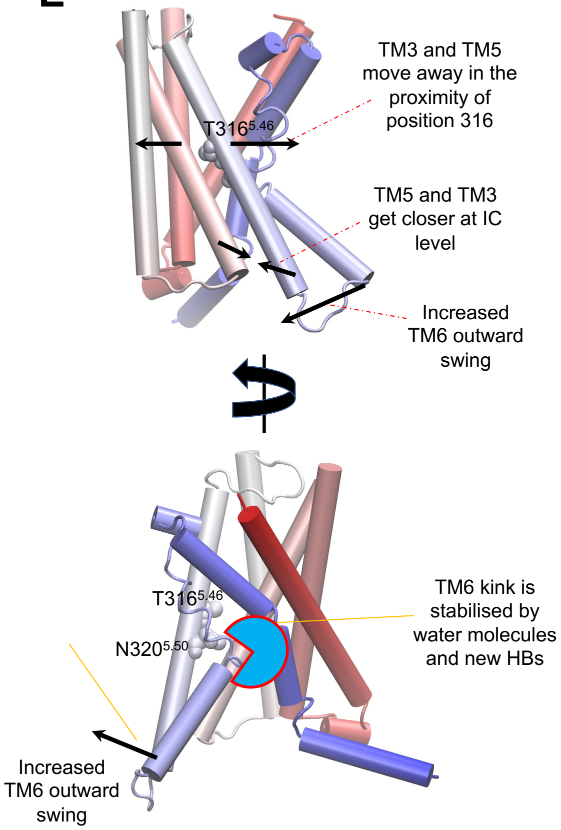

**Fig. S9. MD simulations of GLP-1R A316T TMD.** (**A**) GLP-1R structure used in MD simulations: only the TMD, ECL2, ECL3, and ICLs were retained to highlight the subtle allosteric effect exerted by A316T. (**B**) Comparison between GLP-1R WT and A316T TMD. The side chain of Y242$^{3.45}$ forms a persistent H-bond with the backbone of P312$^{5.42}$ in WT GLP-1R (left), while in A316T the side chain of T316$^{5.46}$ engages Y242$^{3.45}$ (right); as a result of this H-bond reshuffling, the TM3 - TM5 interhelical distance at the L245$^{3.48}$ - V319$^{5.49}$ level (d1) increases from 7.58 ± 0.28 Å to 8.09 ± 0.34 Å, while the TM3 - TM5 interhelical distance at the L251$^{3.54}$ - V327$^{5.57}$ level (d2) decreases from 8.91 ± 0.68 Å to 7.75 ± 0.48 Å; this is accompanied by a stabilisation of the intracellular side of TM5 due to a more persistent H-bond between Y252$^{3.55}$ and R326$^{5.56}$. (**C**) TM5 - TM6 interhelical distances, d1 and d2. (**D**) In A316T, TM6 dynamics is accentuated and the characteristic, active state opening of TM6 (TM6 swing) increases by ~2.5 Å compared to WT [25.12 ± 1.80 Å (WT GLP-1R) *versus* 27.65 ± 2.02 Å (A316T GLP-1R)]. (**E**) Proposed dynamic model for the enhanced constitutive activity of GLP-1R A316T.

**Supplementary Tables**

| **Residue 1** | **Residue 2** | **WT GLP-1R TMD** (% MD frames) | **A316T GLP-1R TMD** (% MD frames) | **WT GLP-1R: GLP-1** (% MD frames) | **A316T GLP-1R: GLP-1** (% MD frames) | **WT GLP-1R: Ex-F1:G$\alpha_s$** (% MD frames) | **A316T GLP-1R: Ex-F1:G$\alpha_s$** (% MD frames) |
|---|---|---|---|---|---|---|---|
| **Y242$^{3.45}$ (sc)** | **A/T316$^{5.46}$ (bb/sc)** | 0.0 | 85.3 | 0.0 | 41.8 | 0.0 | 37.7 |
| **Y242$^{3.45}$ (sc)** | **P312$^{5.42}$ (bb)** | 95.2 | 2.2 | 93.3 | 46.1 | 87.4 | 7.9 |
| **A/T316$^{5.46}$ (bb/sc)** | **P312$^{5.42}$ (bb)** | 0.0 | 97.2 | 0.0 | 49.6 | 0.0 | 86.9 |
| **A/T316$^{5.46}$ (bb/sc)** | **I313$^{5.43}$ (bb)** | 0.0 | 1.6 | 0.0 | 47.2 | 0.0 | 8.3 |

sc =side chain; bb=backbone

**Table S1: H-bond reshuffling close to position 316.** WT GLP-1R and A316T were simulated in 3 different configurations: the isolated TMD in the absence of any agonist, in complex only with the endogenous agonist GLP-1, and in complex with exendin-F1 (Ex-F1) and $G\alpha_s$.

| Residue 1 | Residue 2 | WT GLP-1R TMD (% MD frames) | A316T GLP-1R TMD (% MD frames) | WT GLP-1R: GLP-1 (% MD frames) | A316T GLP-1R: GLP-1 (% MD frames) | WT GLP-1R: Ex-F1:$G\alpha_s$ (% MD frames) | A316T GLP-1R: Ex-F1:$G\alpha_s$ (% MD frames) |
|---|---|---|---|---|---|---|---|
| **A/T316$^{5.46}$** | **Y241$^{3.44}$** | 12.2 | 2.0 | 3.1 | 29.9 | 11.2 | 0.8 |
| **A/T316$^{5.46}$** | **N320$^{5.50}$** | 3.9 | 1.1 | 0.6 | 5.2 | 3.4 | 1.1 |

**Table S2: Water-mediated interactions involving position 316.** WT GLP-1R and A316T were simulated in 3 different configurations: the isolated TMD in the absence of any agonist, in complex only with the endogenous agonist GLP-1, and in complex with exendin-F1 (Ex-F1) and $G\alpha_s$.

| Residue 1 | Residue 2 | WT GLP-1R TMD (% MD frames) | A316T GLP-1R TMD (% MD frames) | WT GLP-1R: GLP-1 (% MD frames) | A316T GLP-1R: GLP-1 (% MD frames) | WT GLP-1R: Ex-F1:$G\alpha_s$ (% MD frames) | A316T GLP-1R: Ex-F1:$G\alpha_s$ (% MD frames) |
|---|---|---|---|---|---|---|---|
| **N320$^{5.50}$ (sc)** | **L360$^{6.49}$ (bb)** | 68.5 | 30.6 | 79.6 | 35.6 | 65.4 | 54.8 |
| **N320 $^{5.50}$ (wb)** | **L360$^{6.49}$ (wb)** | 19.5 | 49.3 | 12.2 | 26.8 | 24.2 | 31.7 |
| **N320$^{5.50}$ (wb)** | **G361$^{6.50}$ (wb)** | 5.3 | 19.5 | 5.9 | 10.3 | 10.9 | 9.8 |

sc =side chain; bb=backbone; wb=water bridge

**Table S3: H-bond and water-mediated interactions involving N320$^{5.50}$ and TM6 kink.** WT GLP-1R and A316T were simulated in 3 different configurations: the isolated TMD in the absence of any agonist, in complex only with the endogenous agonist GLP-1, and in complex with exendin-F1 (Ex-F1) and G$\alpha_s$.

| **Residue 1** | **WT GLP-1R TMD** (% MD frames) | **A316T GLP-1R TMD** (% MD frames) | **WT GLP-1R: GLP-1** (% MD frames) | **A316T GLP-1R: GLP-1** (% MD frames) | **WT GLP-1R: Ex-F1:G$\alpha_s$** (% MD frames) | **A316T GLP-1R: Ex-F1:G$\alpha_s$** (% MD frames) |
|---|---|---|---|---|---|---|
| **A/T316$^{5.46}$** | 1.1 | 1.8 | 0.8 | 6.8 | 1.5 | 5.9 |
| **N320$^{5.50}$** | 2.6 | 4.4 | 3.8 | 15.3 | 4.4 | 11.8 |
| **L360$^{6.49}$** | 2.3 | 3.8 | 2.7 | 11.3 | 3.0 | 7.3 |
| **G361$^{6.50}$** | 2.4 | 3.1 | 9.9 | 13.2 | 6.7 | 14.9 |

**Table S4: Water molecules maximum occupancy nearby positions 316, 320, 360 and 361.** The occupancy of the water molecule with the highest residence time is reported as % of the total MD frames.

| **A316T *versus* WT GLP-1R** | | | | |
|---|---|---|---|---|
| | **Vehicle** | **GLP-1** | **Exendin-4** | **Exendin-F1** |
| **Glucose tolerance** | ↑↑ (0 hr); ↑↑ (6 hr) | - | ↓↓ (0 hr); ↓↓ (6 hr) | ↓↓ (0 hr); ≈ (6 hr) |
| **Plasma insulin** | ↑ (0 hr); ↑↑ (6 hr) | - | ↓ (0 hr); ↓↓ (6 hr) | - |
| **Surface level** | ↓↓ | - | - | - |
| **cAMP generation** | ↑ | - | ↓ | ↑ |
| **Insulin secretion** | ↑↑ (0hr); ↑↑ (6 hr) | - | ↓↓ (0 hr); ↓ (6 hr) | ↓↓ (0 hr); ↓ (6 hr) |
| **Internalisation** | ↑↑ | ↑↑ | ≈ | ≈ |
| **Recycling** | - | ↓↓ | ≈ | ≈ |
| **Degradation** | ↑↑ | ≈ | ≈ | ≈ |
| **$Ca^{2+}$ mobilisation** | ↑↑ | ≈ | ≈ | ↑↑ |

| | | | | |
|---|---|---|---|---|
| **Mini-$G_s$ recruitment** | ↑↑ | ≈ | ≈ | ↑ |
| **β-arrestin 2 recruitment** | ≈ | ≈ | ≈ | ↓ |
| **Endosomal activity** | ↑ | ↑ | ↓ | - |
| **Plasma membrane activity** | ≈ | ≈ | ≈ | - |
| **ECD opening** | ≈ | ↑↑ | - | - |
| **Lipid raft Recruitment** | ↑↑ | ≈ | ↓↓ | - |
| **Membrane diffusion** | ↓↓ | - | ≈ | - |
| **Ubiquitination** | ≈ | - | ↑↑ | - |

**Table S5: Summary of GLP-1R WT *versus* A316T effects.** ↓ reduced; ↑ increased; two arrows, significant; one arrow, tendency; ≈ unchanged; - untested.

| Arrival date | Donor Age | Sex | BMI ($kg/m^2$) |
|---|---|---|---|
| 24.11.21 | 73 | Female | 32 |
| 08.12.21 | 46 | Female | 31.9 |
| 10.12.21 | 67 | Male | 29.7 |
| 08.02.22 | 64 | Male | 26.7 |

**Table S6: Human islet donor characteristics.**

| Primer Name | Sequence |
|---|---|
| PCR forward primer | 5'- GCAGTACTGTGTGTGGCGGCCAATTACTACT- 3' |
| PCR reverse primer | 3'- GTCCATCACAAAGGCAAAGATGACCTCA-5' |
| A316T sequencing primer | 5'- GTACCTGTACACACTGCTGG-3' |

**Table S7: *hGlp1r*$^{A316T/A36T}$ mouse genotyping primers.**

| Gene | Forward (5'→ 3') | Reverse (5'→ 3') |
|---|---|---|
| *Actb* | CACTGTCGAGTCGCGTCC | TCATCCATGGCGAACTGGTG |
| *Mafa* | CTTCAGCAAGGAGGAGGTCATC | CGTAGCCGCGGTTCTTGA |
| *Acot7* | AGATGATTGAGGAGGCCGG | ACAGCGCTCCCCATTCTG |
| *LdhA* | ATGAAGGACTTGGCGGATGA | ATCTCGCCCTTGAGTTTGTCTT |
| *Slc16a1* | GCTTGGTGACCATTGTGGAAT | CCCAGTACGTGTATTTGTAGTCTCCAT |
| *Gcg* | CCAAGAGGAACCGGAACAAC | CCTTCAGCATGCCTCTCAAAT |
| *Ins1* | GCTGGTGGGCATCCAGTAA | AATGACCTGCTTGCTGATGGT |
| *Kcnj11* | CACGGCGGGATAAGTCTACCT | AATCATTTGCCCCCTTCTTGT |

**Table S8: Primers used for qPCR.**

| Imaging | GLP-1:GLP-1R(A316T):DN-G$\alpha_s$:nb35 |
|---|---|
| Magnification (*x*) | 120,000 |
| Accelerating voltage (kV) | 200 |
| Total electron dose ($e^-/Å^2$) | 50 |
| Exposure time (sec) | 7.66 |
| Frames | N/A (EER) |
| Pixel size (Å) | 0.86 |
| Detector | Falcon 4 |
| Acquisition routine | Aberration-free image shift (AFIS) fast acquisition |
| **Processing** | |
| Symmetry imposed | C1 |
| Initial particle images | 1,667,111 |
| Final particle images | 330,000 |
| Resolution (Å) @ FSC = 0.143 | 3.4 |
| **Molecular refinement** | |
| Initial model | PDB 6X18 |
| Chains | 6 |
| Non-hydrogen atoms | 18487 |
| Protein residues | 1172 |
| Waters | 0 |
| Ligands | 0 |

| Bonds (RMSD) | |
|---|---|
| Bond length (Å) | 0.009 |
| Bond angles (º) | 1.28 |
| **Validation** | |
| MolProbity score | 1.23 |
| Clashscore | 2.71 |
| Rotamer outliers (%) | 0.80 |
| CaBLAM outliers (%) | 0.09 |
| **Ramachandran statistics** | |
| Favoured (%) | 97.05 |
| Allowed (%) | 2.95 |
| Disallowed (%) | 0 |

**Table S9: Data collection, processing and refinement statistics for cryo-EM of GLP1-bound GLP-1R (A316T):DN-G$\alpha_s$:Nb35 complex.**

**Movie S1:** T316$^{5.46}$ rotameric states 1 and 2 and a side-by-side comparison of the GLP-1R environment in GLP-1R (WT):GLP-1, GLP-1R (A316T):GLP-1, and GLP-1R (A316T):exendin-F1 (Ex-F1):G$\alpha_s$ near positions 316$^{5.46}$ (A/T), N320$^{5.50}$ and TM6 kink. Y242$^{3.45}$, N320$^{5.50}$, L360$^{6.49}$, and G361$^{6.60}$ are shown as sticks, with position 316$^{5.46}$ (A/T) shown as thicker sticks. Water molecules within 3.5 Å of position 316$^{5.46}$ and N320$^{5.50}$ are shown as red spheres; H-bonds are represented by dashed lines; GLP-1R backbone is shown as transparent white ribbon; GLP-1 is in orange and Ex-F1 in green ribbons.
